# Inhibition of systemic mammalian metabolism by carnitine mimics from the gut microbiota

**DOI:** 10.1101/2025.11.11.687811

**Authors:** Katja Thümmler, Shyam Basak, Heather Hulme, Clio Dritsa, Rachel Foreman, Humaira Naseer, Eidarus Salah, Anthony Tumber, Sam Worboys, Gillian Douce, Jennifer Haggarty, Rachel Williams, Vicky Taylor, Michael J. Ormsby, Phillip D Whitfield, James S. O. McCullagh, Ian P. Salt, Richard Burchmore, Richard J. A. Goodwin, Christopher J. Schofield, Daniel M. Wall

## Abstract

**Background:** The gut microbiota and microbiome-derived metabolites are implicated in various aspects of human health. Here we sought to determine the systemic effects, and mechanism of action, of microbiome-derived carnitine analogues in germ free and conventionally colonised mice.

**Results:** Here we report the systemic localization of the microbiome-derived carnitine analogues, 3-methyl-4-(trimethylammonio)butanoate (3M-4-TMAB) and 5-aminovalerate betaine (5-AVAB), post-administration to germ free mice, with systemic carnitine depletion and mitochondrial dysregulation, reflected in altered acylcarnitine profiles due to incomplete carnitine-mediated fatty acid oxidation. Studies on the inhibitory potency of 3M-4-TMAB at the enzymatic, cellular, and organism levels indicate that, in part, this is a result of inhibition of gammabutyrobetaine hydroxylase, which catalyses the final step in carnitine biosynthesis. Systemic administration of ^13^C- labelled 3M-4-TMAB to conventionally colonised animals to further investigate the physiological relevance of this inhibition, identified a significant reduction in systemic carnitine levels due to increased excretion in urine and faeces and disruption of carnitine-mediated metabolism across ten organs.

**Conclusions:** These results highlight the physiological relevance and significance of microbiome-derived metabolites *in vivo*. The depletion of carnitine and inhibition of its function have potentially long-term impacts on both mammalian energy generation and the protective, signalling and immune regulatory effects of this critical molecule.

## Introduction

Pathways such as fatty acid oxidation (FAO) are essential for supplying mammalian cellular energy needs in the form of adenosine triphosphate (ATP) and metabolic disorders affecting FAO are often developmentally fatal, severely debilitating, or require lifelong treatment^1^. FAO is mediated by carnitine, an omnipresent molecule with roles not only in energy generation, but also signal transduction, epigenetics, tumour biology, neuroprotection and maintenance of immune and mitochondrial homeostasis^2^. Crosstalk between mammalian metabolism and the microbiota is implicated in health and disease studies and gut microbiota mediated effects on metabolism are increasingly being recognised as a modifiable means of influencing mammalian health for positive benefits^3^. Correlative studies have provided insights into potential effects mediated by the microbiota, but further mechanistic studies are critical for understanding how the gut microbiota influences mammalian health^4^.

A number of FAO inhibitors produced by the gut microbiota that mimic carnitine structure have been identified including, 5-AVAB and the *Lachnospiraceae-*derived 3M-4-TMAB and 4-(trimethylammonio)pentanoate (4-TMAP), which are found systemically in mice and which inhibit energy generation in murine white matter cells derived from the region of the brain in which they accumulate^5,6^. When subjected to mass spectrometry these metabolites have an identical mass to charge ratio, one that has been investigated or identified in studies related to several human health conditions with known or suspected metabolic input, including type 1 and type 2 diabetes, autism spectrum disorder, pre-eclampsia and cardiac hypertrophy^7-9^. With lines of communication between the gut microbiota and mitochondria now emerging, it is imperative that steps are taken to understand the consequences of this complex inter-kingdom metabolic interplay^10^. Such an understanding has the potential to provide new mechanistic insights into poorly understood diseases, including those of mitochondrial dysfunction, and general mammalian health.

With this in mind we applied spatial localisation, metabolic profiling and quantitative bioanalysis approaches to germ free (GF) and conventionally colonised animals treated with these microbiome-derived carnitine mimics, identifying metabolic signatures that could be deciphered using mechanistic enzyme assays. This novel approach has identified a clear and systemic role for these microbiome metabolites in mammalian health.

## Results

### 3M-4-TMAB and 5-AVAB localise systemically upon administration to GF mice

To investigate the distribution and effects of microbiome-derived FAO inhibitors, 5-AVAB or 3M-4-TMAB were administered to GF C57BL/6 mice for 5 days (100 mg/Kg intraperitoneally). GF mice were used as they lack a gut microbiota and thus the capability to make 5-AVAB and 3M-4-TMAB. This meant that dosing could be strictly controlled, hence distribution and effects observed could be attributed to the compounds themselves rather than native gut-microbiota-derived metabolites. Mass spectrometry imaging (MSI) indicated that 5-AVAB and 3M-4-TMAB (which have an identical *m/z* of 160.133) localised to all the organs tested including the brain, liver, heart, quadriceps muscle, and colon (Fig. 1). This observation indicates these metabolites not only localise to the liver as expected across the gut-liver axis, but also to distal organs such as the brain across the blood brain barrier, and to other organs with high energy demand such as the heart and quadriceps muscle. Small but significant increases in 5-AVAB accumulation were noted in the brain, colon, muscle and heart in comparison to 3M-4-TMAB (Fig. S1). No significant differences were observed in the liver. The localization of 5-AVAB and 3M-4-TMAB in the brain mirrored what we previously reported for 3M-4-TMAB and carnitine, with the majority of 5-AVAB and 3M-4-TMAB localised in the white matter, in particular the corpus callosum, and with more diffuse distribution in other organs (Fig. 1)^6^.

**Figure 1:**
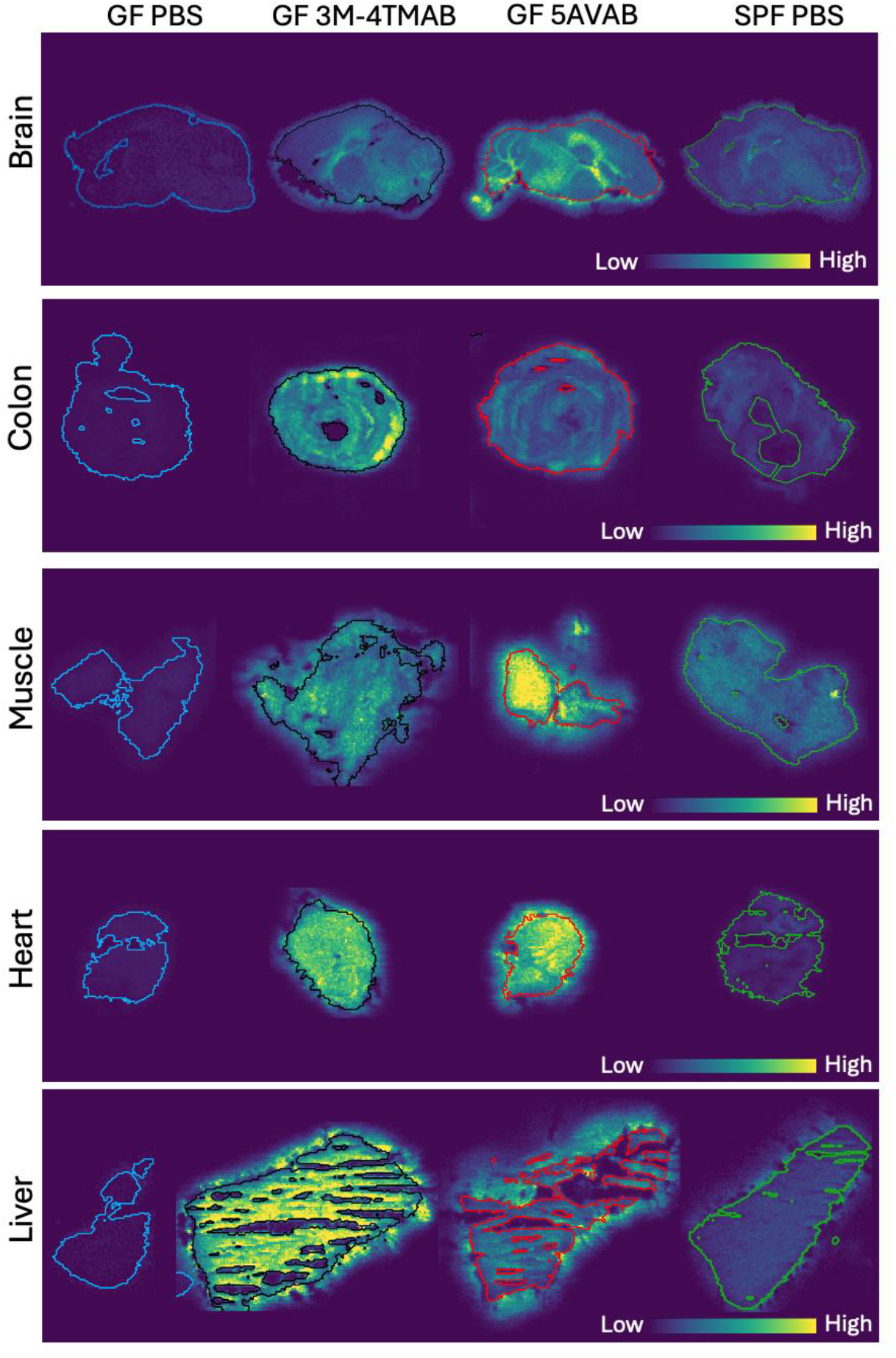
DESI-MSI images of tissues from C57BL/6 SPF or GF mice post-treatment with 3M-4-TMAB, 5-AVAB or PBS. After 5 days of treatment of GF animals with 100 mg/Kg or metabolites of interest or PBS control, tissues were isolated, cryosectioned and subjected to DESI-MSI. Representative brain, colon, muscle, heart and liver sections are shown with GF PBS treatment organs outlined in blue, GF post-3M-4-TMAB treatment outlined in navy, GF post-treatment with 5-AVAB in red, and SPF mouse organs post-treatment with control PBS in green.

### Systemic carnitine and acylcarnitines are significantly reduced by 3M-4-TMAB and 5-AVAB

To investigate the effects of 5-AVAB and 3M-4-TMAB we undertook targeted metabolomics in the form of carnitine quantitation and acylcarnitine profiling across the brain, heart, liver, muscle and colon of 3M-4-TMAB and 5-AVAB treated GF mice, alongside PBS treated control GF mice. Carnitine levels were significantly reduced across all organs tested after treatment with each metabolite (Fig. 2; Fig. S2). While significant fold reductions were observed in carnitine levels across all organs post-treatment of GF mice with 5-AVAB and 3M-4-TMAB, this reduction was comparatively lower in the brain compared to other organs with a 6-fold reduction in colonic carnitine levels post-3M-4-TMAB treatment for example, but only a 40% reduction in carnitine levels observed in the brain after treatment with the same metabolite. A conventionally colonised control group of mice that were specific pathogen free (SPF) was also included in this experiment as these mice contain a microbiota that produces native 3M-4-TMAB and 5-AVAB, and potentially other metabolites of *m/z* 160.133. A significant decrease in carnitine levels was also observed in these control mice in all organs, except brain (Fig. S2).

**Figure 2:**
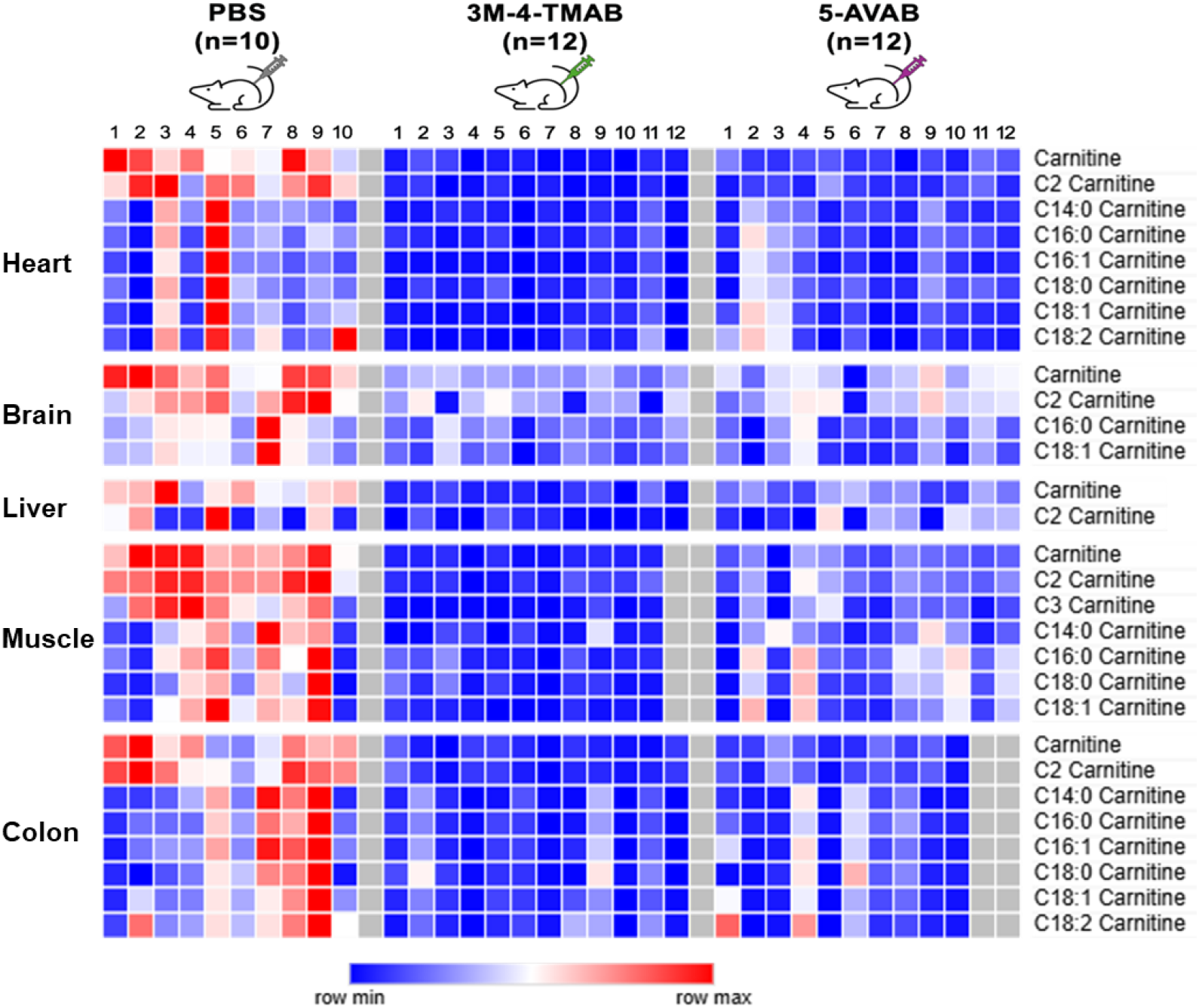
Heat maps showing relative abundance of significantly altered carnitine and acylcarnitines in C57BL/6 GF mice post-treatment with 3M-4-TMAB or 5-AVAB. Summary of molecules significantly changed in C57BL/6 GF mice post-treatment with 3M-4-TMAB and 5-AVAB with PBS treated control C57BL/6 mice shown for comparison. Abundances are shown across 5 tissues, heart, brain, liver, muscle and colon. Data underlying heatmaps are shown in Supplementary Figures S3-S10.

Acylcarnitine profiling is informative in helping to identify disorders of FAO^1^. Both long and short chain acylcarnitines were significantly reduced in all organs in GF mice following treatment with 5-AVAB and 3M-4-TMAB (Fig. 2; Fig. S3-S10). MSI showed that within each organ the presence or absence of acylcarnitines, and their detectability, varied. Acetylcarnitine (C2), the smallest of all acylcarnitines, was detected in all organs and was significantly reduced in each, except for the liver, with these reductions mirroring those seen for carnitine. Propionylcarnitine (C3) was the only odd chain acylcarnitine for which levels were significantly reduced post-3M-4-TMAB and 5-AVAB treatment; and this was only observed in the colon (Fig. S3). The majority of acylcarnitines are even chained meaning any changes will likely be reflected in their abundance in particular, but the reduction in propionylcarnitine seen here may be reflective of reduced propionyl-CoA availability for shuttling to gluconeogenesis. For longer chain acylcarnitines significant reductions were seen post-treatment with 3M-4-TMAB in: myristoylcarnitine (C14) in the heart, muscle, colon; palmitoylcarnitine (C16) in the heart, colon, muscle, brain; palmitoleoylcarnitine (C16:1) in the heart, muscle; stearoylcarnitine (C18:0) in the heart, colon; and oleoylcarnitine (C18:1) in the heart, muscle, colon, brain (Fig. S4-S10). For 5-AVAB significant reductions in acylcarnitines were less frequent and in fewer organs, that is myristoylcarnitine (C14:0) in the heart, palmitoylcarnitine (C16) in the brain and oleoylcarnitine (C18:1) in the muscle, colon, brain.

### 3M-4-TMAB is a competitive inhibitor of the BBOX enzyme that synthesizes carnitine

The significant reduction in carnitine availability observed post-5-AVAB and 3M-4-TMAB treatment in all organs tested is a potential reason underlying the reduced systemic acylcarnitine levels (Fig. 2). To test for potential inhibition of carnitine synthesis the activity of the enzyme that synthesises carnitine from its precursor gammabutyrobetaine (GBB), gammabutyrobetaine hydroxylase (BBOX), was assayed in the presence of both 5-AVAB and 3M-4-TMAB (Fig. S11). Both 5-AVAB and 3M-4-TMAB are structurally similar to mildronate and other known BBOX and FAO inhibitors^11^. The half-maximal inhibitory concentration (IC_50_) for BBOX was 26.9 +/-10.3 μM for 3M-4-TMAB, and >100 μM for 5-AVAB. This was in comparison to the known BBOX inhibitor mildronate (which is competitive substrate) which had an IC_50_ of 62 μM under our conditions^12,13^. This was indicative of 3M-4-TMAB being a more potent inhibitor of BBOX, and thus carnitine synthesis, than mildronate.

FAO inhibition by 3M-4-TMAB and 5-AVAB was investigated in more detail through acylcarnitine profiling *in vitro* using the HepG2 liver cell line, a cell line which synthesises carnitine endogenously through BBOX^14^. For 3M-4-TMAB, disruption of mitochondrial function as evidenced through altered acylcarnitine profiles, occurred at concentrations as low as 25 μM (Fig. S12). However, for 5-AVAB, significant acylcarnitine changes were only detected for one of five acylcarnitines tested and only at higher concentrations than noted with 3M-4-TMAB. For mildronate no inhibition was noted in either acylcarnitine levels or overall FAO (Fig. S12 and Fig. S13). As HepG2s endogenously synthesise carnitine, it may be that this overcame the relatively weak inhibition of BBOX by 5-AVAB. Given the similarity in structure to the known carnitine palmitoyltransferase 1 (CPT1) and carnitine palmitoyltransferase 2 (CPT2) inhibitor aminocarnitine (3-amino-4-trimethylaminobutanoate), it was speculated that 3M-4-TMAB may exert additional inhibitory effects via targeting of these enzymes. However, the decrease in acylcarnitine levels seen with 3M-4-TMAB differed to aminocarnitine treatment in our HepG2 assay where increases were noted in a number acylcarnitines, although these were not significant (Fig. S12). Increased acylcarnitines are indicative of decreased CPT enzyme activity and contrast with our findings with 3M-4-TMAB, indicating it likely does not inhibit CPT1 or CPT2. However, inhibition of carnitine synthesis by 3M-4-TMAB may not be limited to inhibition of the BBOX enzyme alone as a significant decrease in GBB synthesis, the precursor of carnitine, was also noted post-treatment of cells with 3M-4-TMAB (Fig. S14)^15^.

### ^13^C-3M-4-TMAB administration induced systemic metabolic changes and carnitine excretion

To model the interplay between the host, the microbiota and 3M-4-TMAB under more physiologically relevant conditions *in vivo*, ^13^C-labelled 3M-4-TMAB was administered to conventionally colonised C57BL/6 mice. This novel approach provides the metabolite with a unique *m/z* of 163.14336 which facilitated its tracking *in vivo* via MSI, the first time a microbiome-derived metabolite has been re-introduced *in vivo* and systemically localized in this way (Fig. S15). ^13^C-3M-4-TMAB was administered to mice for 5 days and its location and effects tracked. The observed distribution matched what was seen previously with unlabelled 3M-4-TMAB, with ^13^C-3M-4-TMAB present in the white matter of the brain and diffusely present in other organs, including the liver, heart, quadriceps muscle, colon, testes, kidney, lung, ileum and spleen (Fig. 3). Carnitine and acetylcarnitine levels were significantly decreased across all 10 organs tested, while an array of acylcarnitines, both long and short chain, were significantly altered across the different organs (Fig. 3, Fig. S16-S20). The most common significantly altered acylcarnitines were butyrylcarnitine (C4) across 8 of 10 organs, C16 carnitine in 5 organs, and propionylcarnitine (C3) in 4 organs. Significant changes were also detected in undecanoylcarnitine (C11), myristoylcarnitine (C14), palmitoylcarnitine (C16:0), stearoylcarnitine (C18) and oleoylcarnitine (C18:1) in either one or two organs. All organs had significant changes in at least 2, and up to 6, acylcarnitines as well as carnitine itself. The quadriceps muscle was the least affected organ with significant changes in only carnitine and acetylcarnitine.

**Figure 3:**
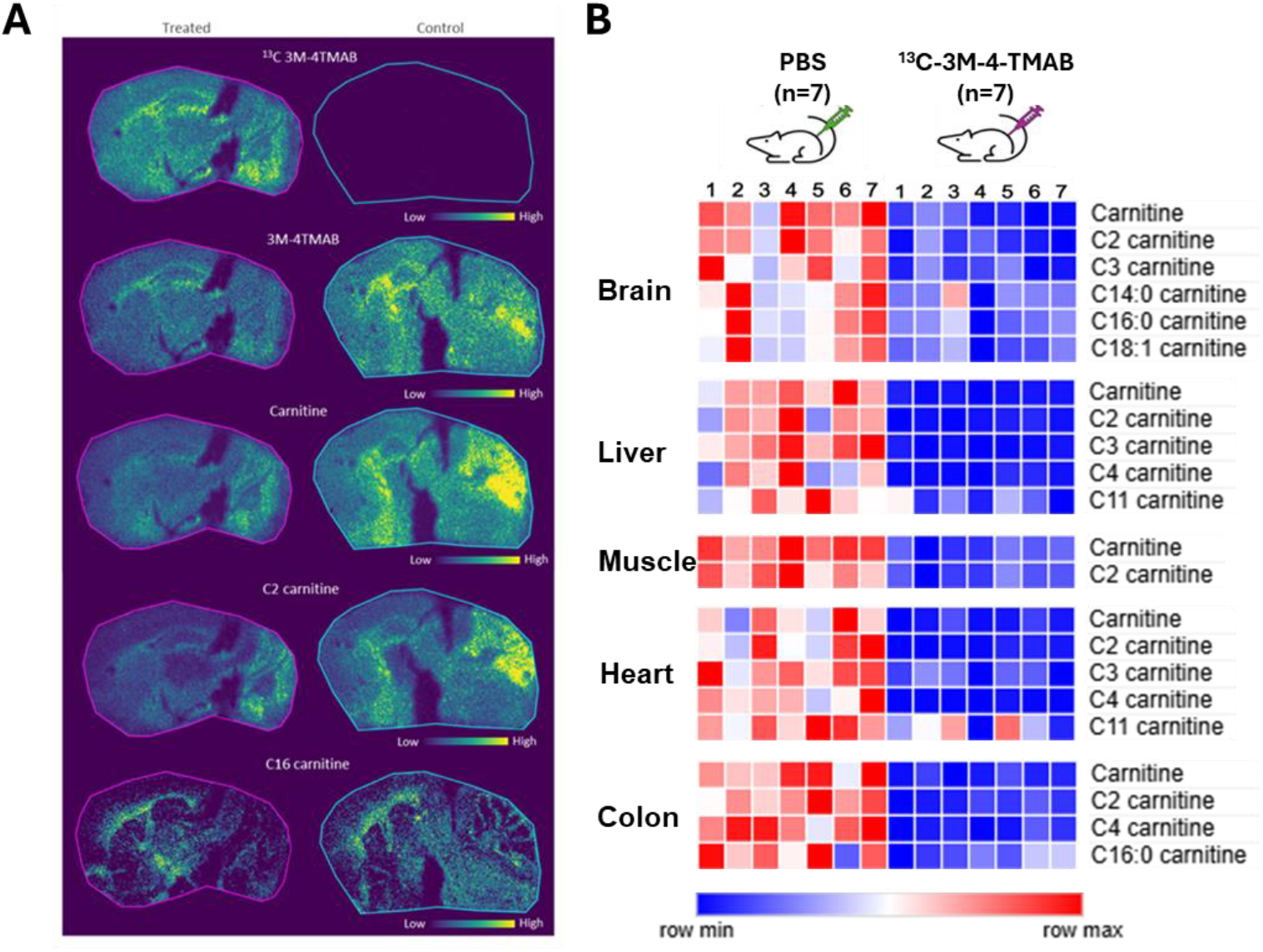
^13^C-3M-4-TMAB, carnitine and acylcarnitine levels in tissues of C57BL/6 SPF mice post-treatment with ^13^C-3M-4-TMAB. (**A**) After 5 days of treatment of animals with 100 mg/Kg of ^13^C-3M-4-TMAB, brains were isolated, cryosectioned and subjected to MSI. Representative brain sections show the distribution of ^13^C-3M-4-TMAB, carnitine and acylcarnitines. (**B**) Heat maps showing relative abundance of ^13^C-3M-4-TMAB and significantly altered carnitine and acylcarnitines in C57BL/6 SPF mice post-treatment. Abundances are shown across 5 tissues; heart, brain, liver, muscle and colon.

A decrease in systemic carnitine levels was observed in individual organs post-treatment and in the serum where carnitine concentration decreased significantly over time (Fig. 3, Fig. 4). We tested excretion of carnitine in response to ^13^C-3M-4-TMAB treatment and noted significantly increased carnitine concentration in urine and to a lesser extent in the faeces post-treatment (Fig. 4). Therefore, the systemic decrease in the carnitine and acylcarnitine pools could be explained by the increased retention of these bacterial metabolites at the expense of native carnitine in these tissues, with carnitine excreted primarily in urine.

**Figure 4:**
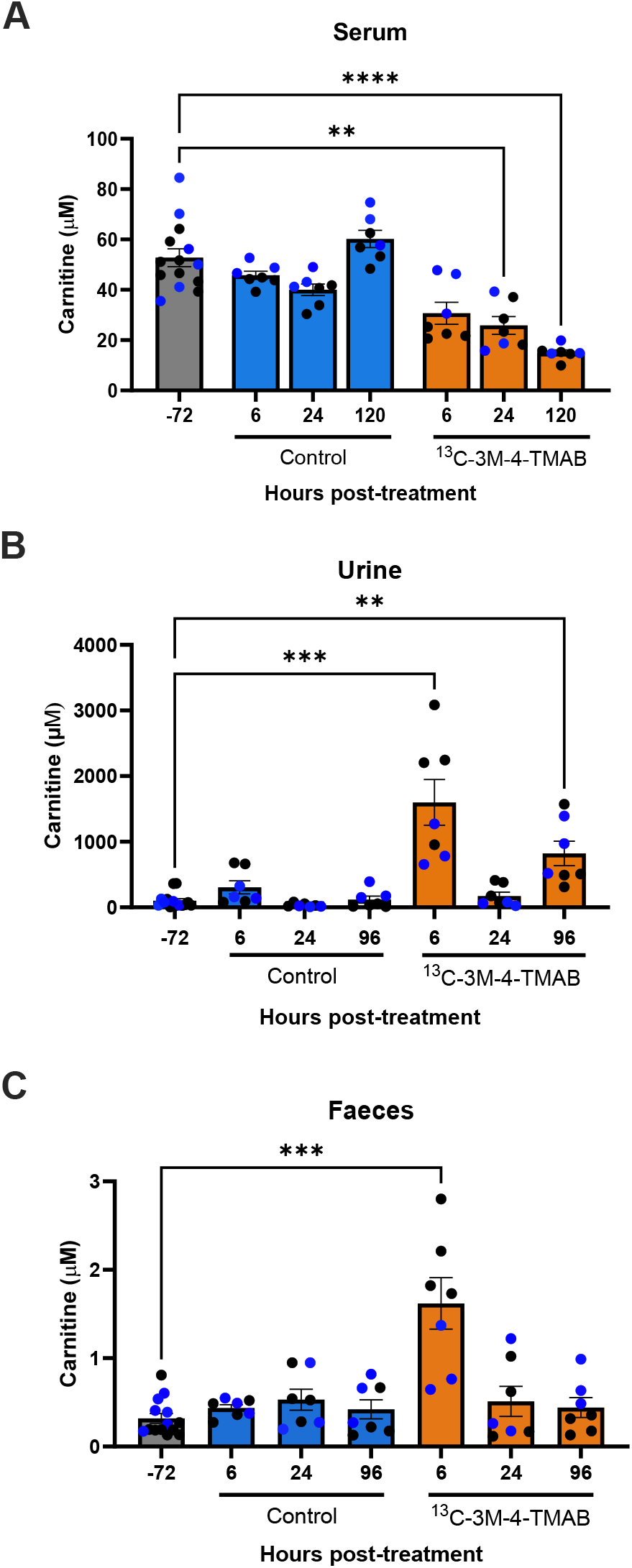
^13^C-3M-4-TMAB treatment of C57BL/6 SPF mice resulted in lower circulating carnitine in the serum and increased carnitine excretion in the urine and faeces. Absolute concentration of carnitine in the serum, urine and faeces 72 hours prior to treatment and over 5 days of ^13^C-3M-4-TMAB administration to C57BL/6 SPF mice (100 mg/Kg). Graphs show mean +/-SEM. Blue dots represent male mice and black dots, female mice. Data are presented as mean ± SEM. Data were analyzed using the Kruskal–Wallis test followed by Dunn’s multiple comparisons test. Asterisks denote significant differences with ** = *p*<0.01, and **** =*p*<0.0001.

Together these data point towards these microbial metabolites exerting significant influence on mammalian physiology in both the long and short terms. Their effects are seen systemically *in vivo*, penetrating and influencing every organ tested and with defined mechanisms of action identified here, targeting carnitine synthesis and inducing its increased excretion. The significance of the microbiome-mammalian crosstalk through such metabolites may, we believe, be critical to understand mammalian health and disease.

## Discussion

It is recognised that the gut microbiota play an important role in mammalian health but mechanisms remain elusive. While many studies focus on the role of diet, nutrient extraction, and the role of the microbiota in modulating and adapting the immune response, less attention has been paid to the role of microbial metabolites in this interkingdom communication. The ability of these metabolites to influence mammalian physiology is likely an underappreciated and underexplored aspect of microbiome influence on health and disease. A number of microbial metabolites have been identified that directly target FAO in the mitochondria with each mimicking the structure of carnitine, or its precursor GBB, and likely inhibiting carnitine function through their interactions with enzymes involved in its synthesis or function^5,6^. These metabolites are worthy of further study for reasons including; their capacity to interfere with essential mammalian energy generation mechanisms, data linking their specific *m/z* with human disease or animal models of disease, and a number of diseases of mitochondrial dysfunction being of unknown cause^5-7,9,16-18^.

We carried out studies on two microbial carnitine mimics, 3M-4-TMAB and 5-AVAB, determining their systemic effects both in the absence of a microbiome, and in the case of 3M-4-TMAB, in the presence of a gut microbiota. It was clear that 3M-4-TMAB and 5-AVAB have the capacity to penetrate all tissues, being found in all organs tested, and in the case of the brain being localised to the white matter. The distribution results, although spatially similar, indicated that 5-AVAB is found at significantly higher levels in the heart, colon, muscle and brain in GF animals in comparison to 3M-4-TMAB. Conversely, despite its lower abundance, it was noted that across all organs 3M-4-TMAB exerted a more significant inhibitory effect on mammalian metabolism, directly inhibiting FAO resulting in significant shifts in acylcarnitine abundances.

The differing metabolite structures clearly have a role to play, with 3M-4-TMAB being a more efficient inhibitor of carnitine synthesis, likely at least in part through inhibition of the BBOX enzyme which catalyses the final step in carnitine biosynthesis, hydroxylation of GBB. Similarly to the known BBOX inhibitor mildronate, it is likely that both 3M-4-TMAB and 5-AVAB have additional mechanisms of action which interfere with carnitine function, including competing with carnitine or acylcarnitine uptake into cells or tissues through their own uptake through channels such as organic cation transporter 2 (OCTN2) that transports carnitine^19^. This possibility is highlighted by our findings with HepG2 cells which synthesise carnitine endogenously, with the inhibitory effect of 5-AVAB being diminished in contrast to 3M-4-TMAB^14^. This observation may be indicative of the inhibitory effects of 3M-4-TMAB and 5-AVAB being more pronounced outside tissues such as the liver, kidney and central nervous system which are capable of endogenously synthesizing carnitine and overcoming the inhibitory effects of these metabolites^20^. In tissues such as muscle where BBOX activity has not been detected, there is likely a dependence on the systemic carnitine pool to replenish local needs, but it is difficult to determine whether the inhibitory effects seen here in muscle are due to this systemic depletion of carnitine, or more local inhibitory effects.

Carnitine or acylcarnitine deficiency has been linked to diseases across multiple organs, with carnitine known to play important roles in apoptosis prevention, epigenetic regulation, maintenance of immune and mitochondrial homeostasis and signal transduction^2,21-23^. Conversely the decreases noted in acylcarnitine levels here could have beneficial effects given that acylcarnitine accumulation has known pro-inflammatory effects and acylcarnitine accumulation has been shown to have detrimental effects in metabolic disorders such as type 2 diabetes (T2D)^24,25^. Interestingly, treatment with metformin, a commonly used therapeutic in T2D patients, results in a significant increase in serum concentrations of a molecule of unknown identity, but with an identical mass to charge ratio to that of 3M-4-TMAB and 5-AVAB^26^. Whether increases in molecules of this particular mass to charge ratio have a role in disease outcomes is unclear, but FAO defects including accumulation of a variety of acylcarnitines have been clearly linked to T2D and are biomarkers of defective FAO in T2D^27^. This raises the possibility that gut microbiome derived metabolites could mediate some of the positive effects of metformin for which multiple mechanisms of action have been proposed^28^.

The role of carnitine can vary between organs, playing specific context and location-dependent roles including those independent of metabolism and FAO. Accumulation in the white matter of the brain of 5-AVAB and 3M-4-TMAB may counteract carnitine function where it has been positively associated with brain size and development in pre-term infants, while carnitine deficiency and FAO defects have been linked to autism, as has 5-AVAB^7,29-31^. Additionally, the significant decrease in acetylcarnitine noted post-treatment with 5-AVAB and 3M-4-TMAB is likely to have multiple neurological effects given the roles of acetylcarnitine in promoting acetylcholine synthesis, protection of the developing brain, facilitating neurotransmitter release and synaptic plasticity^32,33^. In the liver carnitine reduces ammonia toxicity resulting in its use as a treatment in diseases such as hepatic encephalopathy^34^. Carnitine reabsorption in the liver also plays a critical role in controlling systemic levels of the molecule and therefore the accumulation to high levels of 3M-4-TMAB in this organ may have wider implications for systemic carnitine levels. Indeed, significant increases in carnitine excretion in the urine were noted post-^13^C-3M-4-TMAB treatment and the role of inhibition of carnitine reabsorption in the liver in this excretion is worthy of further investigation.

BBOX depletion selectively inhibits growth of triple-negative breast cancer cells and has recently been identified as a cancer target. These cells rely heavily on FAO for energy generation and often overexpress BBOX to supply sufficient carnitine to maintain their FAO-dependent energy needs^35^. Therefore, gut microbiome metabolites such as 3M-4-TMAB, that directly inhibit BBOX, may exert some therapeutic benefits in the context of some cancers and indeed other diseases. FAO inhibitors are used therapeutically for angina as they shift the demand for energy precursors from fatty acids to glucose, reducing oxygen demand, and thus stress^36^. Mildronate, also known as Meldonium, has been used as a metabolic modulator in the context of performance enhancement, shifting energy demands in the body to more efficient use of glucose during periods of intense exercise and recovery^37^. The emergence of gut microbiota-derived functional mimics of such metabolic modulators will likely lead to more focus on the gut microbiota in the context of elite sport where use of Meldonium is already banned.

Together the data presented here indicate a far wider ranging role for these microbiome-derived metabolites than previously described. The loss of carnitine through excretion and inhibition of its synthesis results in a systemic deficit in this important mediator of energy generation. Whether the systemic distribution of these metabolites has negative consequences due to FAO inhibition, or whether they play an important role in balancing fatty acid and other energy source use, is still to be determined. However, the results presented here are the first to show a direct influence of microbial metabolites on systemic mammalian physiology. This has longer term implications on how we view communication between the gut microbiota and both healthy mammalian physiology and disease.

## Material and Methods

### Animal work

Seven-to 8-week-old male C57BL/6J GF and SPF mice were sourced from the University of Manchester, Gnotobiotic Facility. Both GF and SPF mice were fed the same pelleted diet, which was sterilised by irradiation with 50 kGy. Mice were treated daily for 5 days via intraperitoneal (i.p.) injection on alternating flanks with 100 mg/Kg 3M-4-TMAB, or 5-AVAB in 100 μL phosphate buffered saline (PBS) or PBS alone. For treatment with ^13^C-labelled 3M-4TMAB, seven-to eight-week-old male and female C57BL/6J mice from Charles River were housed under a 12-hour light/dark cycle under pathogen-free conditions with *ad libitum* access to food and water. Mice were injected i.p. on alternating flanks with 100 mg/kg [^13^C]-labelled 3M-4-TMAB in 100 μL PBS, or PBS alone for control mice, daily for 5 days and blood, urine and faecal samples taken 72 hours prior to treatment and at 6, 24, 96 and 120 hours post-treatment. All experimental treatments were carried out between 8-10 am.

### Tissue processing

Mice were culled 24 hours after the final i.p. injection and organs dissected immediately, colons and ileums were co-embedded in 2.5% hydroxypropyl-methylcellulose (HPMC) and 7.5 % polyvinylpyrrolidone (PVP) hydrogel (both from Merck) and snap frozen in dry ice-chilled isopropanol followed by isopentane^38^. All other organs were first snap frozen in iso-pentane on dry ice and later embedded in HPMC/PVP hydrogel and frozen in isopropanol on dry-ice and washed with dry-ice chilled isopentane. Ten micrometre thick tissue sections were cut at -16ºC using a cryostat microtome (Thermo Scientific) and thaw mounted on Superfrost glass slides (Thermo Scientific) or on conductive indium tin oxide (ITO) slides (Bruker Daltonik) for MALDI MSI.

### Matrix-assisted laser desorption ionization (MALDI)-MSI

MALDI matrix, 2,5-Dihydroxybenzoic acid (DHB) was dissolved in 50% acetonitrile, 50% water and 0.1% trifluoracetic acid to a concentration of 37.5 mg/ml. DHB was deposited across the tissue samples with an HTX TM-sprayer (HTX Technologies, LLC) with 8 passes, a flow rate of 80 μl/min, a nozzle temperature of 75°C, velocity of 1200 mm/minute and track spacing of 3 mm. MALDI-MSI was performed using a timTOF Flex (Bruker Daltonics) in positive ion mode, with a spatial resolution of 50 μm, 200 laser shots per position and *m/z* range of 50-1000. A collision energy of 10 eV a Collision RF of 600 Vpp, a transfer rime of 55 μs and a pre pulse storage of 5 μs was used. Laser power was optimised for the experiment.

### Desorption electrospray ionization (DESI)-MSI

DESI-MSI was performed using a Q-Exactive mass spectrometer (Thermo Scientific, Bremen, Germany) equipped with an automated 2D-DESI ion source (Prosolia Inc., Indianapolis, IN, USA). The machine was operated in positive ion mode, with a mass range of *m/z* 80-900, with a nominal mass resolution of 70,000. The injection time was fixed to 150 milliseconds (ms) resulting in a scan rate of 3.8 pixels/s. The spatial resolution used was 70 μm. A home-built Swagelok DESI sprayer was operated with a mixture of 95%_v/v_ methanol:5%_v/v_ water delivered with a flow rate of 1.5 μL/min and nebulised with nitrogen at a backpressure of 6 bar. The data files in .raw format were converted into .mzML files using ProteoWizard msConvert (version 3.0.4043) and subsequently compiled to an .imzML file (imzML converter version 1.3). All subsequent data processing was performed in SCiLS Lab (version 2024, Bruker Daltonik, Bremen, Germany). Identification of metabolites of interest was performed by comparison of the measured accurate mass to theoretical.

### Quantitative bioanalysis of mouse serum, urine and faeces

Mouse blood was collected by tail vein puncture prior to start of the treatment, during treatment and postmortem. Serum was then collected after coagulation for 1 hour at room temperature and centrifugation (10,000 g, 15 minutes, 4ºC). Faecal samples were weighed and diluted to 200 mg/mL in 0.2% _v/v_ formic acid in acetonitrile:water (10:90) and subjected to rapid vortex mixing. The homogenised samples were centrifuged for 3 minutes at 3500 g and the supernatant was taken for analysis.

Intermediate solutions of ^13^C-3M-4-TMAB and carnitine were diluted to working concentrations (0.2% _v/v_ formic acid in acetonitrile:water (10:90)) and used to make calibration standards and quality controls (QC) in either PBS or control C57B/6J mouse serum, urine and faeces (BioIVT). Calibration curves were prepared in PBS for all analytes, at two relevant linear ranges for accurate quantitation (low range; 2 – 2000 nM and high range; 0.2 – 100 μM). QCs were either prepared in 10% mouse serum (PBS) with non-endogenous ^13^C-3M-4-TMAB, and by dilution of each matrix at three levels (1:25, 1:50, 1:100) to ensure robust quantitation of the endogenous carnitine.

For extraction 10 μL calibration, QC and diluted study samples (in PBS) were aliquoted into a LoBind 96-well plate, 10 μL internal standard working solution (100 nM carnitine d9 in 0.2% _v/v_ formic acid in acetonitrile:water (10:90)) was added and the plate was mixed for 1 minute at 1000 rpm. To each sample, 180 μL of 0.2% _v/v_ formic acid in acetonitrile:water (90:10) was added before being mixed for 2 minutes at 1000 rpm and centrifuged (3500 g, 5 minutes). An aliquot of 50 μL was transferred to clean plate, diluted with either 50 μL 0.2%_v/v_ formic acid in 10 mM ammonium formate (aq) (low range) or 200 μL 0.2% _v/v_ formic acid in acetonitrile:water (50:50) (high range) for analysis.

### Liquid Chromatography Tandem Mass Spectrometry (LC-MS/MS)

All experimental procedures were performed using an I-Class Acquity (Waters, USA) LC system coupled to an AB SCIEX 5000 triple quadrupole mass spectrometer (Sciex, USA). Chromatographic data and regression were processed using Sciex Analyst (version 1.7.2); any additional data analyses were performed in Microsoft Excel. The extracted sample (5 μL) was injected onto a Waters Acquity BEH-Amide column (1.7μM, 2.1 x 50mm) set to 30°C, with starting mobile phases set to 45% _v/v_ A (0.2% _v/v_ formic acid in 10 mM ammonium formate(aq)) and 55% _v/v_ B (0.2% _v/v_ formic acid in 5 mM ammonium formate: acetonitrile 10:90) and a flow rate of 0.6 mL/min. The analytes were separated over a 2-minute gradient from 55 to 25% _v/v_ B, followed by a flush of 5% _v/v_ B for 1 minute before returning to initial conditions, resulting in an overall runtime of 3 minutes. The sample manager wash solvent was phase A, the sample purge was phase B, and seal wash solvent was 50% _v/v_ acetonitrile (aq).

Turbo Spray ionisation was applied in positive mode within the heated mass spectrometer source (400°C), with an ionspray voltage of 5 kV, curtain gas of 25 psi and a high collision gas setting. The target analytes and internal standard were detected using multiple reaction monitoring (45 ms dwell time) and using a collision energy of 35 V and declusting potential of 90 V, specific Q1 and Q3 ions were targeted for each analyte; 3M-4-TMAB, Q1=160, Q3=58.6; ^13^C-3M-4-TMAB, Q1=163,Q3=55.1; carnitine, Q1=162, Q3=85.0; carnitine d9 (internal standard), Q1=171, Q3=69.0.

### Statistical analysis

Statistical analyses were performed using GraphPad Prism 10 (GraphPad Software Inc.). Depending on the experimental design, comparisons were made using t-tests or one-way ANOVA (with post hoc tests and sample sizes detailed in the figure legends or main text). When data did not meet the assumptions of normality required for ANOVA, the nonparametric Kruskal–Wallis test was used instead. A *p*-value of <0.05 was considered statistically significant.

## Declarations

### Ethics approval and consent to participate

All procedures were approved by University of Glasgow and University of Manchester ethics committees under U.K. Home Office Project licence PP1122278.

### Consent for publication

Not applicable.

### Funding

This work was supported by UKRI Biotechnology and Biological Sciences Research Council (BBSRC) grant numbers BB/K008005/1 to R.B., R.G., and DMW and BB/V001876/1 to R.B., R.G., C.J.S. and DMW. C.D. was funded through a University of Glasgow MVLS doctoral training programme PhD studentship. C.J.S. thanks Cancer Research UK. The Manchester Gnotobiotic Facility was established with the support of the Wellcome Trust (097820/Z/11/B) using founder mice obtained from the Clean Mouse Facility, University of Bern.

### Availability of data and materials

All data needed to evaluate the conclusions in the paper are present in the paper and/or the Supplementary Materials. Raw data related to this paper are freely available in the University of Glasgow online repository https://doi.org/10.5525/gla.researchdata.1965.

## Acknowledgements

Not applicable.

## Author contributions statement

K.T., S.B., H.H., R.F., H.N., S.W., R.W., V.T., M.J.O., P.D.W., J.S.O.M., I.P.S., R.B., R.J.A.G., C.J.S., and D.M.W. planned the studies. K.T., S.B., H.H., C.D., R.F., H.N., E.S., A.T., S.W., G.D., J.H., R.W. conducted the experiments. K.T., S.B., H.H., R.F., H.N., R.W., P.D.W., J.S.O.M., I.P.S., R.B., R.J.A.G., and D.M.W interpreted the studies. D.M.W. wrote the first draft of the paper. All authors reviewed, edited, and approved the paper. R.J.A.G., R.B., C.J.S. and D.M.W. obtained funding.

## Competing interests statement

The authors declare that there are no competing interests.

## Notes

https://doi.org/10.5525/gla.researchdata.1965

### Competing Interest Statement

The authors have declared no competing interest.

## References and Notes

1. Merritt JL, 2nd, MacLeod E, Jurecka A, Hainline B. Clinical manifestations and management of fatty acid oxidation disorders. Rev Endocr Metab Disord. Dec 2020;21(4):479–493. doi:10.1007/s11154-020-09568-3

2. Xiang F, Zhang Z, Xie J, et al. Comprehensive review of the expanding roles of the carnitine pool in metabolic physiology: beyond fatty acid oxidation. J Transl Med. Mar 14 2025;23(1):324. doi:10.1186/s12967-025-06341-5

3. Aron-Wisnewsky J, Warmbrunn MV, Nieuwdorp M, Clement K. Metabolism and Metabolic Disorders and the Microbiome: The Intestinal Microbiota Associated With Obesity, Lipid Metabolism, and Metabolic Health-Pathophysiology and Therapeutic Strategies. Gastroenterology. Jan 2021;160(2):573–599. doi:10.1053/j.gastro.2020.10.057

4. Fan Y, Pedersen O. Gut microbiota in human metabolic health and disease. Nat Rev Microbiol. Jan 2021;19(1):55–71. doi:10.1038/s41579-020-0433-9

5. Karkkainen O, Tuomainen T, Koistinen V, et al. Whole grain intake associated molecule 5-aminovaleric acid betaine decreases beta-oxidation of fatty acids in mouse cardiomyocytes. Sci Rep. Aug 29 2018;8(1):13036. doi:10.1038/s41598-018-31484-5

6. Hulme H, Meikle LM, Strittmatter N, et al. Microbiome-derived carnitine mimics as previously unknown mediators of gut-brain axis communication. Sci Adv. Mar 2020;6(11):eaax6328. doi:10.1126/sciadv.aax6328

7. Ottosson F, Russo F, Abrahamsson A, et al. Unraveling the metabolomic architecture of autism in a large Danish population-based cohort. BMC Med. Jul 19 2024;22(1):302. doi:10.1186/s12916-024-03516-7

8. Wang Z, Yang S, Liu L, et al. The gut microbiota-derived metabolite indole-3-propionic acid enhances leptin sensitivity by targeting STAT3 against diet-induced obesity. Clin Transl Med. Dec 2024;14(12):e70053. doi:10.1002/ctm2.70053

9. Zhao M, Wei H, Li C, et al. Gut microbiota production of trimethyl-5-aminovaleric acid reduces fatty acid oxidation and accelerates cardiac hypertrophy. Nat Commun. Apr 1 2022;13(1):1757. doi:10.1038/s41467-022-29060-7

10. Yardeni T, Tanes CE, Bittinger K, et al. Host mitochondria influence gut microbiome diversity: A role for ROS. Sci Signal. Jul 2 2019;12(588)doi:10.1126/scisignal.aaw3159

11. Corner TP, Tumber A, Salah E, et al. Derivatives of the Clinically Used HIF Prolyl Hydroxylase Inhibitor Desidustat Are Efficient Inhibitors of Human gamma-Butyrobetaine Hydroxylase. J Med Chem. May 8 2025;68(9):9777–9798. doi:10.1021/acs.jmedchem.5c00586

12. Henry L, Leung IK, Claridge TD, Schofield CJ. gamma-Butyrobetaine hydroxylase catalyses a Stevens type rearrangement. Bioorg Med Chem Lett. Aug 1 2012;22(15):4975–8. doi:10.1016/j.bmcl.2012.06.024

13. Leung IK, Krojer TJ, Kochan GT, et al. Structural and mechanistic studies on gamma-butyrobetaine hydroxylase. Chem Biol. Dec 22 2010;17(12):1316–24. doi:10.1016/j.chembiol.2010.09.016

14. Zhan Y, Dong X, Yang M, et al. Gamma-butyrobetaine hydroxylase (BBOX1) exerts suppressive effects on HepG2 hepatoblastoma cells. Med Oncol. Sep 27 2024;41(11):253. doi:10.1007/s12032-024-02496-1

15. Chegary M, Te Brinke H, Doolaard M, et al. Characterization of L-aminocarnitine, an inhibitor of fatty acid oxidation. Mol Genet Metab. Apr 2008;93(4):403–10. doi:10.1016/j.ymgme.2007.11.001

16. Tsutsui H, Maeda T, Min JZ, et al. Biomarker discovery in biological specimens (plasma, hair, liver and kidney) of diabetic mice based upon metabolite profiling using ultra-performance liquid chromatography with electrospray ionization time-of-flight mass spectrometry. Clin Chim Acta. May 12 2011;412(11-12):861–72. doi:10.1016/j.cca.2010.12.023

17. Haukka JK, Sandholm N, Forsblom C, Cobb JE, Groop PH, Ferrannini E. Metabolomic Profile Predicts Development of Microalbuminuria in Individuals with Type 1 Diabetes. Sci Rep. Sep 14 2018;8(1):13853. doi:10.1038/s41598-018-32085-y

18. Jaaskelainen T, Karkkainen O, Jokkala J, et al. A Non-Targeted LC-MS Profiling Reveals Elevated Levels of Carnitine Precursors and Trimethylated Compounds in the Cord Plasma of Pre-Eclamptic Infants. Sci Rep. Oct 2 2018;8(1):14616. doi:10.1038/s41598-018-32804-5

19. Longo N, Frigeni M, Pasquali M. Carnitine transport and fatty acid oxidation. Biochim Biophys Acta. Oct 2016;1863(10):2422–35. doi:10.1016/j.bbamcr.2016.01.023

20. Rebouche CJ, Engel AG. Tissue distribution of carnitine biosynthetic enzymes in man. Biochim Biophys Acta. Jun 5 1980;630(1):22–9. doi:10.1016/0304-4165(80)90133-6

21. Madiraju P, Pande SV, Prentki M, Madiraju SR. Mitochondrial acetylcarnitine provides acetyl groups for nuclear histone acetylation. Epigenetics. Aug 16 2009;4(6):399–403. doi:10.4161/epi.4.6.9767

22. Hassan N, Rashad M, Elleithy E, Sabry Z, Ali G, Elmosalamy S. L-Carnitine alleviates hepatic and renal mitochondrial-dependent apoptotic progression induced by letrozole in female rats through modulation of Nrf-2, Cyt c and CASP-3 signaling. Drug Chem Toxicol. Mar 2023;46(2):357–368. doi:10.1080/01480545.2022.2039180

23. Huang H, Liu N, Guo H, et al. L-carnitine is an endogenous HDAC inhibitor selectively inhibiting cancer cell growth in vivo and in vitro. PLoS One. 2012;7(11):e49062. doi:10.1371/journal.pone.0049062

24. McCoin CS, Knotts TA, Adams SH. Acylcarnitines--old actors auditioning for new roles in metabolic physiology. Nat Rev Endocrinol. Oct 2015;11(10):617–25. doi:10.1038/nrendo.2015.129

25. Liepinsh E, Makrecka-Kuka M, Makarova E, et al. Acute and long-term administration of palmitoylcarnitine induces muscle-specific insulin resistance in mice. Biofactors. Sep 10 2017;43(5):718–730. doi:10.1002/biof.1378

26. Adam J, Brandmaier S, Leonhardt J, et al. Metformin Effect on Nontargeted Metabolite Profiles in Patients With Type 2 Diabetes and in Multiple Murine Tissues. Diabetes. Dec 2016;65(12):3776–3785. doi:10.2337/db16-0512

27. Fernandez-Duval G, Razquin C, Wang F, et al. A multi-metabolite signature robustly predicts long-term mortality in the PREDIMED trial and several US cohorts. Metabolism. Mar 17 2025:156195. doi:10.1016/j.metabol.2025.156195

28. Foretz M, Guigas B, Viollet B. Metformin: update on mechanisms of action and repurposing potential. Nat Rev Endocrinol. Aug 2023;19(8):460–476. doi:10.1038/s41574-023-00833-4

29. Manninen S, Silvennoinen S, Bendel P, Lankinen M, Schwab US, Sankilampi U. Carnitine Intake and Serum Levels Associate Positively with Postnatal Growth and Brain Size at Term in Very Preterm Infants. Nutrients. Nov 9 2022;14(22)doi:10.3390/nu14224725

30. Bankaitis VA, Xie Z. The neural stem cell/carnitine malnutrition hypothesis: new prospects for effective reduction of autism risk? J Biol Chem. Dec 13 2019;294(50):19424–19435. doi:10.1074/jbc.AW119.008137

31. Xie Z, Jones A, Deeney JT, Hur SK, Bankaitis VA. Inborn Errors of Long-Chain Fatty Acid beta-Oxidation Link Neural Stem Cell Self-Renewal to Autism. Cell Rep. Feb 9 2016;14(5):991–999. doi:10.1016/j.celrep.2016.01.004

32. Smeland OB, Meisingset TW, Borges K, Sonnewald U. Chronic acetyl-L-carnitine alters brain energy metabolism and increases noradrenaline and serotonin content in healthy mice. Neurochem Int. Jul 2012;61(1):100–7. doi:10.1016/j.neuint.2012.04.008

33. Traina G, Bernardi R, Rizzo M, Calvani M, Durante M, Brunelli M. Acetyl-L-carnitine up-regulates expression of voltage-dependent anion channel in the rat brain. Neurochem Int. Jun 2006;48(8):673–8. doi:10.1016/j.neuint.2005.11.005

34. Tani J, Morishita A, Sakamoto T, et al. L-carnitine reduces hospital admissions in patients with hepatic encephalopathy. Eur J Gastroenterol Hepatol. Feb 1 2021;32(2):288–293. doi:10.1097/MEG.0000000000001748

35. Liao C, Zhang Y, Fan C, et al. Identification of BBOX1 as a Therapeutic Target in Triple-Negative Breast Cancer. Cancer Discov. Nov 2020;10(11):1706–1721. doi:10.1158/2159-8290.CD-20-0288

36. Sesti C, Simkhovich BZ, Kalvinsh I, Kloner RA. Mildronate, a novel fatty acid oxidation inhibitor and antianginal agent, reduces myocardial infarct size without affecting hemodynamics. J Cardiovasc Pharmacol. Mar 2006;47(3):493–9. doi:10.1097/01.fjc.0000211732.76668.d2

37. Lippi G, Mattiuzzi C. Misuse of the metabolic modulator meldonium in sports. J Sport Health Sci. Mar 2017;6(1):49–51. doi:10.1016/j.jshs.2016.06.008

38. Dannhorn A, Kazanc E, Ling S, et al. Universal Sample Preparation Unlocking Multimodal Molecular Tissue Imaging. Anal Chem. Aug 18 2020;92(16):11080–11088. doi:10.1021/acs.analchem.0c00826

39. Tars K, Leitans J, Kazaks A, et al. Targeting carnitine biosynthesis: discovery of new inhibitors against gamma-butyrobetaine hydroxylase. J Med Chem. Mar 27 2014;57(6):2213–36. doi:10.1021/jm401603e

40. Gasparik V, Greney H, Schann S, et al. Synthesis and biological evaluation of 2-aryliminopyrrolidines as selective ligands for I1 imidazoline receptors: discovery of new sympatho-inhibitory hypotensive agents with potential beneficial effects in metabolic syndrome. J Med Chem. Jan 22 2015;58(2):878–87. doi:10.1021/jm501456p

41. Royer F, Felpin FX, Doris E. Synthesis of gamma-amino acids by rearrangement of alpha-cyanocyclopropanone hydrates: application to the regioselective labeling of amino acids. J Org Chem. Sep 21 2001;66(19):6487–9. doi:10.1021/jo0104227

42. Reihill JA, Ewart MA, Salt IP. The role of AMP-activated protein kinase in the functional effects of vascular endothelial growth factor-A and -B in human aortic endothelial cells. Vasc Cell. Apr 20 2011;3:9. doi:10.1186/2045-824X-3-9

43. Folch J, Lees M, Sloane Stanley GH. A simple method for the isolation and purification of total lipides from animal tissues. J Biol Chem. May 1957;226(1):497–509.

