## Supplementary Material for "Inhibition of systemic mammalian metabolism by carnitine mimics from the gut microbiota"

Short title: Gut microbial metabolites and mammalian health

Thümmler *et al*.

**Supplementary Material**

**Synthesis of metabolites**

**1. General synthesis information**

Reagents were from commercial sources (Sigma-Aldrich, Inc.; Fluorochem Ltd; Tokyo Chemical Industries, BLDpharm) and were used as received. Anhydrous solvents were kept under an atmosphere of nitrogen. HPLC grade solvents (Sigma-Aldrich Inc.) were used for purifications, reaction workups and extractions.

Nuclear magnetic resonance (NMR) spectroscopy was performed using a Bruker Avance III 400 HD Nanobay NMR machine with a 9.4T magnet or Bruker Avance III 500 HD NMR machine with a 11.75T magnet or Bruker NEO 600 with broadband helium cryoprobe equipped with a 14.1T magnet. 3-(Trimethylsilyl)propionic-2,2,3,3-d₄ acid sodium salt (TSP) or sodium trimethylsilylpropanesulfonate (DSS) are used as internal standards for quantitative NMR analysis when D₂O was used as the solvent. Chemical shifts for ¹H NMR are reported in parts per million (ppm) downfield from tetramethylsilane (TMS) when CDCl₃ was the solvent, or from TSP/DSS when D₂O was used. Shifts are referenced to the residual proton signal of CDCl₃ (δ 7.26 ppm) or the –Si(CH₃)₃ signal of TSP/DSS (δ 0.00 ppm). For ¹³C NMR, chemical shifts are reported relative to the solvent signal (CDCl₃: δ 77.16 ppm) or to the –Si(CH₃)₃ peak of TSP/DSS (δ 0.00 ppm). NMR data are reported as follows: chemical shift, multiplicity (s: singlet, d: doublet, t: triplet, q: quartet, m: multiplet), coupling constant (*J*, Hz; accurate to 0.1 Hz), and integration. NMR spectra are shown in the Supporting Information (Fig. S22-S30).

**2. General procedure for the treatment with Amberlite IRN78 (OH) ion-exchange resin**

A solution of ester or acid salt of betaine compound in water (0.25 mL/mmol) was allowed to soak into Amberlite IRN78 (OH) ion-exchange resin column (4 mL/mmol). After 0.5 h, the column slowly was eluted with water. The eluate was evaporated, the residue was azeotropically dried with 2-propanol followed by drying *in vacuo* to obtain the desired betaine compound^39^.

**3. Synthetic procedures and analytical data**

**Synthesis of 3M-4TMAB**

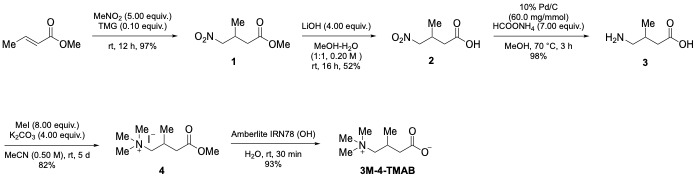

**Methyl 3-methyl-4-nitrobutanoate (1)**

**2** (15.0 g, 93.3 mmol, yield: 97%) was synthesised as yellowish liquid following reported method.^2^

^1^H NMR (400 MHz, 300 K, CDCl_3_) δ 4.47 (dd, *J* = 12.1, 6.3 Hz, 1H), 4.34 (dd, *J* = 12.1, 7.1 Hz, 1H), 3.69 (s, 3H), 2.85 – 2.70 (m, 1H), 2.50 – 2.31 (m, 2H), 1.09 (d, *J* = 6.8 Hz, 3H) ppm; ^13^C NMR (101 MHz, 300 K, CDCl_3_) δ 171.8, 80.3, 52.0, 37.8, 29.5, 17.4 ppm.

The spectroscopic data are consistent with those previously reported^40^.

**3-Methyl-4-nitrobutanoic acid (2)**

A 500 mL round-bottomed flask equipped with a magnetic stirrer bar was charged with **1** (5.62 g, 34.9 mmol) and MeOH-H_2_O (1:1, 180 mL). Lithium hydroxide hydrate (4.41 g, 105 mmol) was added to the mixture portion wise. The reaction mixture was then stirred for 18 h at rt.; MeOH was evaporated *in vacuo* and the reaction mixture was diluted with water (50 mL). The aqueous portion was washed with EtOAc (2 X 40 mL), then acidified to pH 2–3 by adding 2 N HCl at 0 °C and extracted with EtOAc (3 X 150 mL). The organic extracts were combined, washed with brine, dried over MgSO_4_, filtered, and concentrated *in vacuo*. Purification by column chromatography using a Biotage Selekt purification machine (wavelengths monitored: 200 – 400 nm) (100 g Sfär Silica Duo; 80 mL/min using 100% (v/v) cyclohexane (+ 0.1% (v/v) formic acid) (2 CV), followed by a linear gradient (25 CV): 0%→100% (v/v) EtOAc (+ 0.1% (v/v) formic acid) in cyclohexane (+ 0.1% (v/v) formic acid)) gave **2** as a yellow gummy liquid (2.68 g, 18.2 mmol, 52%).

^1^H NMR (400 MHz, 300 K, CDCl_3_) δ 4.48 (dd, *J* = 12.2, 6.3 Hz, 1H), 4.36 (dd, *J* = 12.2, 7.0 Hz, 1H), 2.87 – 2.72 (m, 1H), 2.53 (dd, *J* = 16.6, 6.5 Hz, 1H), 2.43 (dd, *J* = 16.6, 6.9 Hz, 1H), 1.13 (d, *J* = 6.8 Hz, 3H) ppm; ^13^C NMR (101 MHz, 300 K, CDCl_3_) δ 177.8, 80.1, 37.7, 29.3, 17.4 ppm.

**4-Amino-3-methylbutanoic acid (3)**

A 250 mL round-bottomed flask equipped with a magnetic stirrer bar was charged with **2** (2.71 g, 18.4 mmol) and MeOH (110 mL). Pd/C (10% w/w, 1.10 g) and ammonium formate (8.14 g, 129 mmol) were carefully added portion wise. The reaction mixture was heated to 70 °C with a reflux condenser and stirred for 3 h. Then the reaction mixture was cooled and filtered through Celite, and filtrate was concentrated and dried *in vacuo* to give a sticky gum, which was washed with diethyl ether to obtain **3** (2.12 g, NMR purity: >99%, 18.1 mmol, 98%) as a white solid.

^1^H NMR (400 MHz, 300 K, D_2_O) δ 2.98 (dd, *J* = 12.8, 6.0 Hz, 1H), 2.87 (dd, *J* = 12.8, 7.1 Hz, 1H), 2.29 (dd, *J* = 13.3, 5.6 Hz, 1H), 2.25 – 2.09 (m, 2H), 1.01 (d, *J* = 6.5 Hz, 3H) ppm; ^13^C NMR (101 MHz, 300 K, D_2_O) δ 180.8, 44.9, 42.5, 29.3, 16.7 ppm.

The spectroscopic data are consistent with those previously reported^41^.

**4-Methoxy-*N*,*N*,*N*,2-tetramethyl-4-oxobutan-1-aminium iodide (4)**

A 100 mL sealed tube equipped with a magnetic stirrer bar was charged with **3** (2.10 g, 17.9 mmol), potassium carbonate (9.90 g, 71.6 mmol) and acetonitrile (36 mL). Iodomethane (8.9 mL, 143 mmol) was added to the mixture. The resultant mixture was stirred at rt for 5 d at room temperature. Then volatiles were removed *in vacuo*. The solid was washed with EtOAc (20 mL). Then the solid was suspended in CHCl_3_ (75 mL) and filtered. The filtrate was concentrated and dried to afford the compound **4** as an off white solid (4.44 mg, NMR purity: >99%, 14.7 mmol, yield: 82%).

^1^H NMR (400 MHz, 300 K, D_2_O) δ 3.72 (s, 3H), 3.41 (dd, *J* = 13.7, 2.3 Hz, 1H), 3.36 – 3.27 (m, 1H), 3.16 (s, 9H), 2.63 – 2.47 (m, 3H), 1.18 (d, *J* = 6.5 Hz, 3H) ppm; ^13^C NMR (101 MHz, 300 K, D2O) δ 174.5, 71.8 (t, *J* = 2.9 Hz), 53.5 (t, *J* = 4.1 Hz), 52.4, 40.1, 25.3, 20.1 ppm.

**3-Methyl-4-(trimethylammonio)butanoate (3M-4TMAB)**

General procedure for the treatment with Amberlite IRN78 (OH) ion-exchange resin was followed using **5** (4.44 g, NMR purity: >99%, 14.7 mmol) to afford 3M-4-TMAB (2.46 g, NMR purity: 88%, 13.6 mmol, yield: 93%) as an off white solid.

^1^H NMR (600 MHz, 300 K, D_2_O, TSP) δ 3.40 – 3.24 (m, 2H), 3.16 (s, 9H), 2.51 – 2.38 (m, 1H), 2.33 (dd, *J* = 14.5, 6.7 Hz, 1H), 2.22 (ddd, *J* = 14.5, 7.8, 0.8 Hz, 1H), 1.17 (d, *J* = 6.8 Hz, 3H) ppm; ^13^C NMR (151 MHz, 300 K, D_2_O, TSP) δ 183.0, 75.2 (t, *J* = 2.7 Hz), 56.2 (t, *J* = 4.1 Hz), 47.3, 29.5, 23.0 ppm.

The spectroscopic data are consistent with those previously reported^6^.

**Synthesis of ^13^C-labelled 3M-4TMAB**

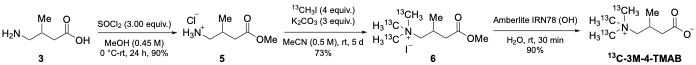

**4-Methoxy-2-methyl-4-oxobutan-1-aminium chloride (5)**

A 10 mL round-bottomed flask equipped with a magnetic stirrer bar was charged with amino acid (**3**, 286 mg, 2.00 mmol, 82% pure) and MeOH (4.00 mL) under nitrogen atmosphere. The mixture was cooled in an ice bath and thionyl chloride (438 μL, 6.00 mmol) was added dropwise. Then the ice bath was removed, and the resultant mixture was stirred at rt for 24 h; volatiles were then removed *in vacuo*. The reaction mixture was washed with diethyl ether and acetone mixture (10 mL, 5:1) and the residue was dried under high vacuum over night to afford the compound **5** as brown semi solid (304 mg, NMR purity: >99%, 1.80 mmol, yield: 90%).

^1^H NMR (600 MHz, 300 K, D_2_O, DSS) δ 3.71 (s, 3H), 3.05 (dd, *J* = 13.0, 5.9 Hz, 1H), 2.89 (dd, *J* = 13.0, 8.0 Hz, 1H), 2.51 (dd, *J* = 16.0, 6.4 Hz, 1H), 2.41 (dd, *J* = 15.9, 7.3 Hz, 1H), 2.35 – 2.25 (m, 1H), 1.04 (d, *J* = 7.2 Hz, 3H) ppm; ^13^C NMR (151 MHz, 300 K, D_2_O, DSS) δ 177.7, 55.0, 47.0, 40.9, 31.0, 19.1 ppm.

**4-Methoxy-2-methyl-*N*,*N*,*N*-tri(methyl-^13^*C*)-4-oxobutan-1-aminium iodide (6)**

A 10 mL microwave vial equipped with a magnetic stirrer bar was charged with **5** (212 mg, 1.27 mmol), potassium carbonate (524 mg, 3.79 mmol) and acetonitrile (2.50 mL). 2 M solution of iodomethane-^13^C in in tert-butyl methyl ether (2.54 mL, 5.08 mmol) was added to the mixture. The resultant mixture was stirred at rt for 5 days at room temperature; the volatiles were then removed *in vacuo*. The resultant solid was washed with EtOAc (3 mL). Then the solid was suspended in CHCl_3_ (10 mL) and filtered. The filtrate was concentrated and dried to afford the compound **6** as an off white solid (214 mg, NMR purity: >99%, 0.70 mmol, yield: 55%).

^1^H NMR (500 MHz, D_2_O, TSP) δ 3.73 (s, 3H), 3.47 – 3.27 (m, 2H), 3.17 (dt, *J*(^1^H-^13^C) = 144.3, 3.5 Hz, 9H), 2.63 – 2.50 (m, 3H), 1.19 (d, *J* = 6.6 Hz, 3H) ppm; ^13^C NMR (151 MHz, 300 K, D_2_O, TSP) δ 177.3, 74.6 (*t*, J = 3.3 Hz), 56.2 (t, *J* = 3.8 Hz), 55.2, 42.9, 28.1, 22.8 ppm.

**3-Methyl-4-(tri(methyl-^13^*C*)ammonio)butanoate (^13^C-3M-4-TMAB)**

General procedure for the treatment with Amberlite IRN78 (OH) ion-exchange resin was followed using **6** (210 mg, NMR purity: >99%, 0.69 mmol) to afford ^13^C-3M-4-TMAB (107 mg, NMR purity: 94%, 0.62 mmol, yield: 90%) as an off white solid.

^1^H NMR (600 MHz, 300 K, D_2_O, TSP) δ 3.39 – 3.24 (m, 2H), 3.04 (dt, *J*(^1^H-^13^C) = 136.3, 3.5 Hz, 9H), 2.49 – 2.39 (m, 1H), 2.33 (dd, *J* = 14.5, 6.7 Hz, 1H), 2.22 (dd, *J* = 14.5, 7.8 Hz, 1H), 1.17 (d, *J* = 6.7 Hz, 3H) ppm; ^13^C NMR (151 MHz, 300 K, D_2_O, TSP) δ 183.1, 75.2 (t, *J* = 2.7 Hz), 56.2 (t, *J* = 3.8 Hz), 47.3, 29.5, 23.0 ppm.

The spectroscopic data are consistent with those previously reported data for 3M-4-TMAB^6^.

**Synthesis of 5-AVAB**

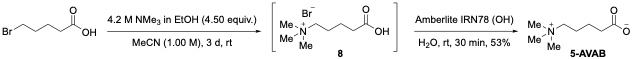

**5-(Trimethylammonio)pentanoate (5-AVAB)**

**5-AVAB** (1.23 g, NMR purity: >99%, 7.72 mmol, yield: 53%) was synthesised as a white solid following reported method.^1^

^1^H NMR (600 MHz, 300 K, D_2_O, TSP) δ 3.39 – 3.29 (m, 2H), 3.11 (s, 9H), 2.26 (td, *J* = 7.3, 1.6 Hz, 2H), 1.86 – 1.76 (m, 2H), 1.68 – 1.57 (m, 2H) ppm; ^13^C NMR (151 MHz, 300 K, D_2_O, TSP) δ 185.4, 69.2 (t, *J* = 2.9 Hz), 55.7 (t, *J* = 4.2 Hz), 39.6, 25.4, 25.0 ppm.

The spectroscopic data are consistent with those previously reported^39^.

**FAO inhibition assay**

HepG2 cells were cultured in Eagle’s Minimum Essential Media (Merck) supplemented with 2 mM L-glutamine (Merck), 1% penicillin/streptomycin (Merck) and 10% foetal bovine serum (FBS, Invitrogen). Cells were fed twice a week, sub-cultured at 80% confluency and maintained at 37°C, 5% CO₂. HepG2 cells (1 x10^5^/well) were seeded into a 12-well plate and next day transferred into low serum media (3% FBS) and pretreated with 75 mM 3M-4TMAB, and the known fatty acid oxidation inhibitors mildronate (Merck) and aminocarnitine (Santa Cruz) for 72 hours^15,36^. Thereafter, cells were washed extensively with Earle’s HEPES buffer (5.3 mM KCl, 116 mM NaCl, 0.8 mM MgSO4, 1 mM NaH2PO4, 1.8 mM CaCl2, 20 mM HEPES-NaOH, pH 7.4) and incubated with [^3^H]-palmitic acid (2 mCi/mL, 110 mM), carnitine (50 μM), fatty acid–free BSA (0.5 mg/mL) and 75 μM 3M-4TMAB, mildronate, aminocarnitine or etomoxir (Merck) in Earle’s HEPES buffer for 35 mins at 37 ºC. ^3^H_2_0 produced from [^3^H]-palmitate was separated and quantified as described previously^42^.

**Carnitine and GBB quantification**

HepG2 cells (0.5 x 10^6^ per well) were incubated in 6 well plates in EMEM with 10% FBS overnight and next day transferred into reduced serum media (3% FBS) and treated with 75 μM 3M-4-TMAB, 4-TMAP, mildronate or aminocarnitine for 72 h. Cells were washed twice with ice cold PBS, trypsinised and cell pellets resuspended in ice cold 80% methanol. To aid complete cell lysis, cells were frozen at -150ºC for 1 hour; after thawing cells suspensions were sonicated for 10 minutes, centrifuged (13,000 g 30 min, 4ºC) and supernatants collected and analysed by liquid chromatography mass spectrometry (LCMS). Samples were analysed using an Atlantis Premier BEH Z-HILIC column (1.7 µm, 2.1 × 150 mm, Waters, Wilmslow, UK) with an Ultimate 3000 RSLC system (Thermo Fisher Scientific, Hemel Hempstead, UK) coupled to an Orbitrap Exploris™ 240 mass spectrometer (Thermo Scientific, Bremen, Germany) operated in positive ion mode with heated electrospray ionization (H-ESI). Mobile phase A consisted of 100% acetonitrile, and mobile phase B was 5 mM ammonium formate in Type-1 water. Chromatographic separation was achieved using the following gradient at a flow rate of 0.4 mL/min: 0–2 min, 5% B; 6 min, 15% B; 14 min, 60% B; 15 min, 60% B; 16 min, 5% B; 30 min, 5% B. The column was maintained at 50 °C, with the autosampler set to 4 °C, and 5 µL of sample was injected per run.

MS data were acquired using a top 10 data-dependent MS/MS (ddMS2) method. Source parameters were as follows: spray voltage, +3.5 kV; sheath, auxiliary, and sweep gas flow rates of 25, 8, and 0 (arbitrary units), respectively; ion transfer tube and vaporizer temperatures were both set to 300 °C. Full scan spectra were collected at a resolution of 60,000 over an m/z range of 60–900, with two microscans, an AGC target of 5 × 10⁶, and a maximum injection time of 120 ms. ddMS2 spectra were acquired at a resolution of 15,000 using a 2 m/z isolation window, normalized HCD collision energy of 35%, two microscans, an AGC target of 1 × 10⁵, and a maximum injection time of 80 ms. The raw data were analysed using Progenesis QI (Waters, Wilmslow, UK) subsequently the carnitine and GBB peaks were annotated by matching retention time, accurate m/z (± 5 ppm), and fragmentation pattern scoring against authentic standards run with the same method to achieve a level 1 identification.

**Acylcarnitine profiling**

80% confluent monolayer of HepG2 cells (12 well plates) were treated with different concentrations (25-75 μM) of 3M-4TMAB, 4-TMAP, 5-AVAB, mildronate or aminocarnitine for 48 hours in low serum EMEM media (3% FBS). Cells were then washed twice with ice cold PBS, trypsinised and cell pellets resuspended in 0.2 mL water and 1 mL methanol. Folch extraction and LC/MS analysis were performed by MVLS Shared Research Facility (University of Glasgow). Lipids from HepG2 cells were extracted according to the method of Folch *et al*^43^. In brief, cellular lysates were extracted with 6 ml chloroform/methanol (2/1, v/v and C14:0-*d_9_* (2 pmol), 4 pmol C16:0-*d_3_* (4 pmol) and C18:0-*d_3_* (2 pmol) acylcarnitines were added as internal standards. The mixture was then left to stand at 4°C for 1 h. The samples were partitioned by the addition of 1.3 ml 0.1 M KCl and the mixture was centrifuged to facilitate phase separation. The lower chloroform layer was evaporated to dryness under nitrogen gas and reconstituted in methanol containing 5 mM ammonium formate. Acylcarnitines were analysed by LC-MS using a Thermo Exactive Orbitrap mass spectrometer equipped with a heated electrospray ionization (HESI) probe and interfaced with a Dionex UltiMate 3000 RSLC system (Thermo Fisher Scientific, Hemel Hempstead, UK). Samples (10 µl) were injected onto a Thermo Hypersil Gold C18 column (2.1 mm by 100 mm; 1.9 μm) maintained at 50°C. Mobile phase A consisted of water containing 10 mM ammonium formate and 0.1% (v/v) formic acid. Mobile phase B consisted of a 90:10 mixture of isopropanol:acetonitrile containing 10 mM ammonium formate and 0.1% (v/v) formic acid. The initial conditions for analysis were 65% mobile phase A, 35% mobile phase B and the percentage of mobile phase B was increased from 35 to 65% over 4 minutes, followed by 65% to 100% over 15 minutes, with a hold for 2 minutes before re-equilibration to the starting conditions over 6 minutes. The flow rate was 400 μl/min and samples were analysed in positive ion mode over the mass-to-charge ratio (*m/z*) range of 250 to 2,000 at a resolution of 100,000. The signals corresponding to the accurate *m/z* values for [M+H]^+^ ions of acylcarnitine molecular species were extracted from raw LC-MS data sets with the mass error set to 5 parts per million (ppm). Quantification was achieved by relating the peak area of acylcarnitine species to the respective internal standard. The concentrations of individual acylcarnitines were normalised to cell number.

**Supplementary Figures**

**Figure S1**

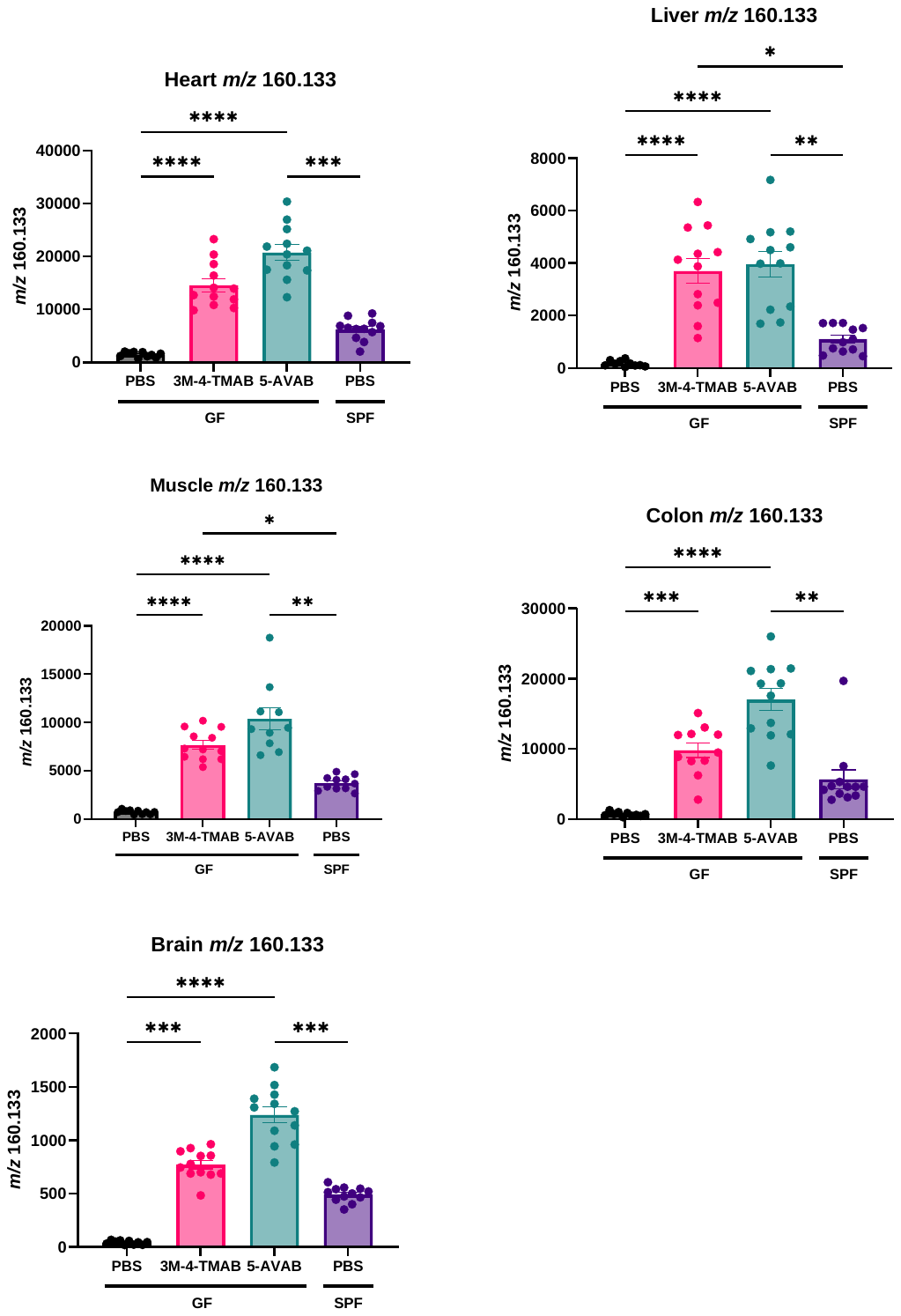

**Figure S1: Relative abundance of *m/z* 1601.133 across sections from organs from C57BL/6 GF mice treated with metabolites or PBS and C57BL/6 SPF mice treated with PBS.** Tissues were isolated from mice, cryosectioned and imaged via DESI-MSI for the presence of *m/z* 160.133, the known *m/z* 3M-4-TMAB and 5-AVAB. GF mice were treated with PBS (control) or 100 mg/Kg metabolite (3M-4-TMAB or 5-AVAB) for 5 days via i.p. injection. SPF control mice were treated with PBS. Data are presented as mean ± SEM. Data were analyzed using the Kruskal–Wallis test followed by Dunn's multiple comparisons test. Asterisks denote significant differences with * = *p*<0.05, ** = *p*<0.01, *** = *p*<0.001, and **** =*p*<0.0001.

**Figure S2**

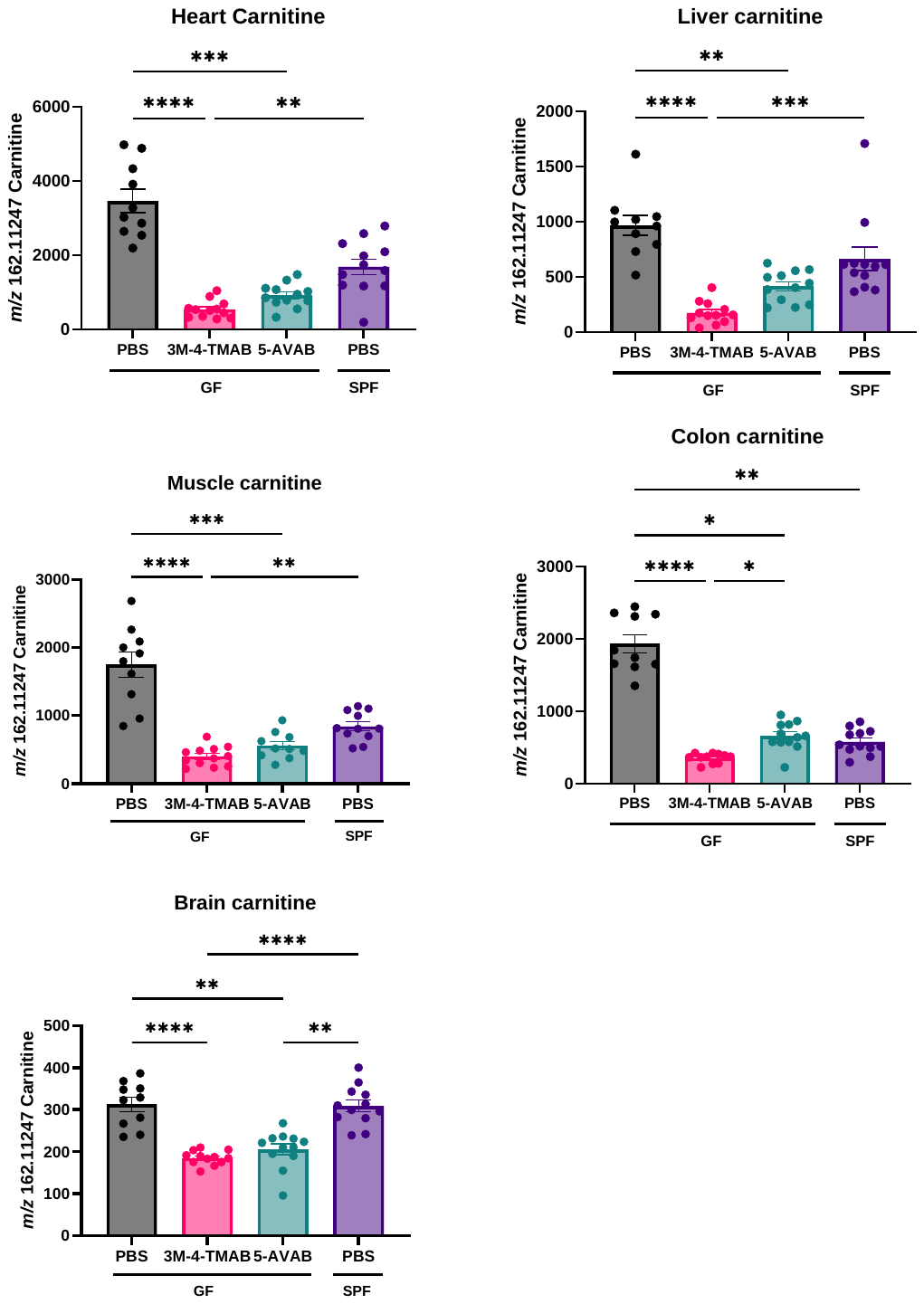

**Figure S2: Relative abundance of carnitine across sections from organs from C57BL/6 GF mice treated with metabolites or PBS and C57BL/6 SPF mice treated with PBS.** Tissues were isolated from mice, cryosectioned and imaged via DESI-MSI for the presence of carnitine. GF mice were treated with PBS (control) or 100 mg/Kg metabolite (3M-4-TMAB or 5-AVAB) for 5 days via i.p. injection. SPF control mice were treated with PBS. Data are presented as mean ± SEM. Data were analyzed using the Kruskal–Wallis test followed by Dunn's multiple comparisons test. Asterisks denote significant differences with * = *p*<0.05, ** = *p*<0.01, *** = *p*<0.001, and **** =*p*<0.0001.

**Figure S3**

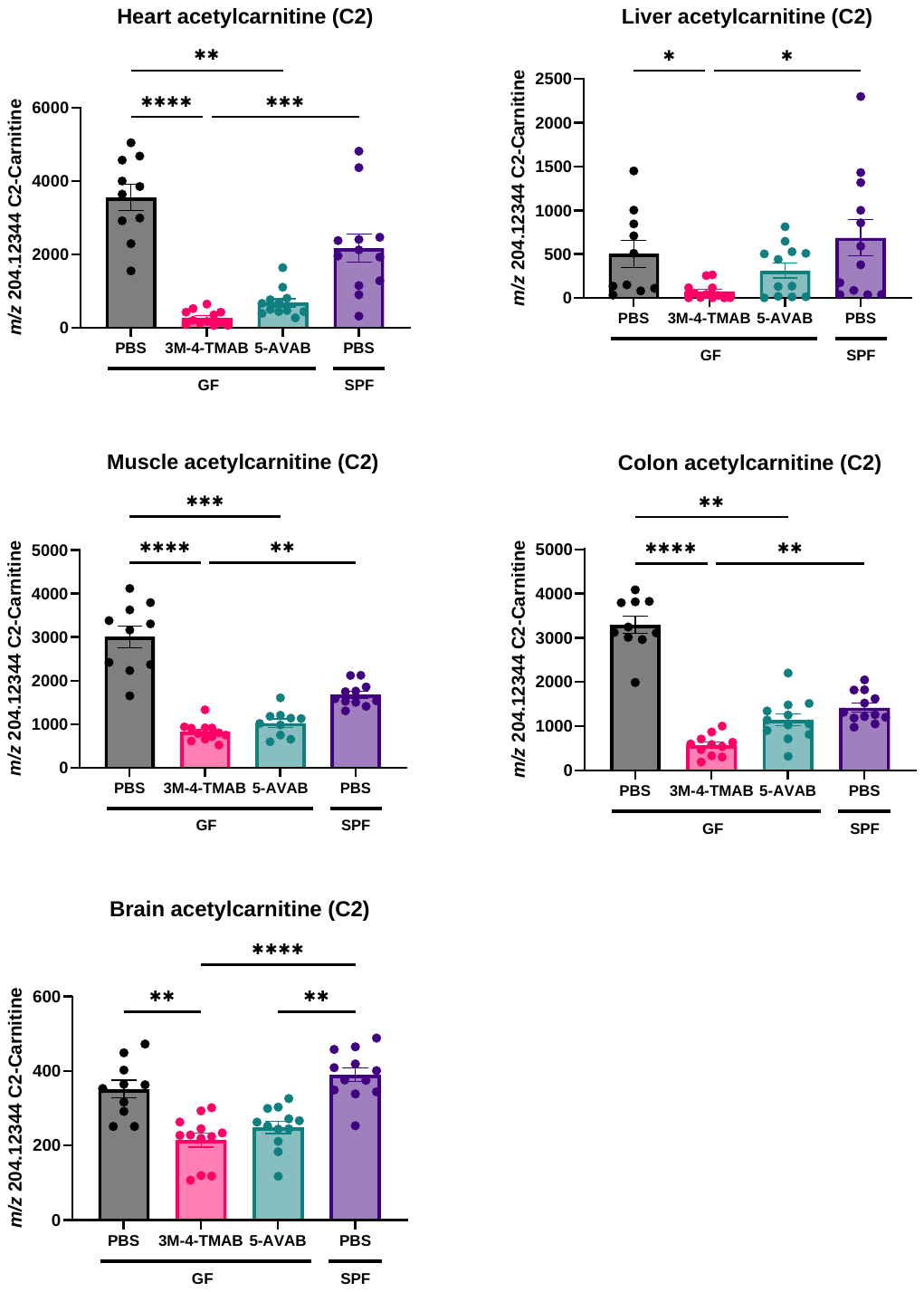

**Figure S3: Relative abundance of acetylcarnitine (C2) across sections from organs from C57BL/6 GF mice treated with metabolites or PBS or C57BL/6 SPF mice treated with PBS.** Tissues were isolated from mice, cryosectioned and imaged via DESI-MSI for the presence of acetylcarnitine. GF mice were treated with PBS (control) or 100 mg/Kg metabolite (3M-4-TMAB or 5-AVAB) for 5 days via i.p. injection. SPF control mice were treated with PBS. Data are presented as mean ± SEM. Data were analyzed using the Kruskal–Wallis test followed by Dunn's multiple comparisons test. Asterisks denote significant differences with * = *p*<0.05, ** = *p*<0.01, *** = *p*<0.001, and **** =*p*<0.0001.

**Figure S4**

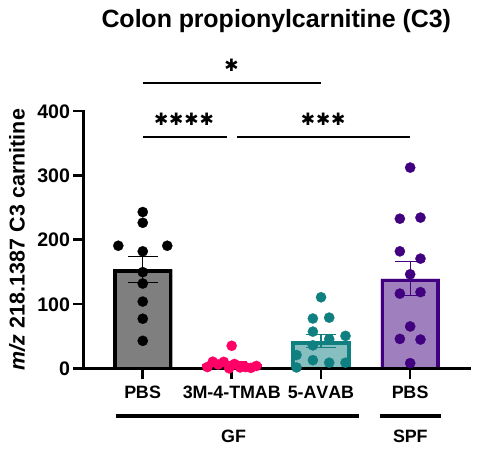

**Figure S4: Relative abundance of propionylcarnitine (C3) in the colon of C57BL/6 GF mice treated with metabolites or PBS and C57BL/6 SPF mice treated with PBS.** Colon tissue was isolated from mice, cryosectioned and imaged via DESI-MSI for the presence of propionylcarnitine. GF mice were treated with PBS (control) or 100 mg/Kg metabolite (3M-4-TMAB or 5-AVAB) for 5 days via i.p. injection. SPF control mice were treated with PBS. Data are presented as mean ± SEM. Data were analyzed using the Kruskal–Wallis test followed by Dunn's multiple comparisons test. Asterisks denote significant differences with * = *p*<0.05, ** = *p*<0.01, *** = *p*<0.001, and **** =*p*<0.0001.

**Figure S5**

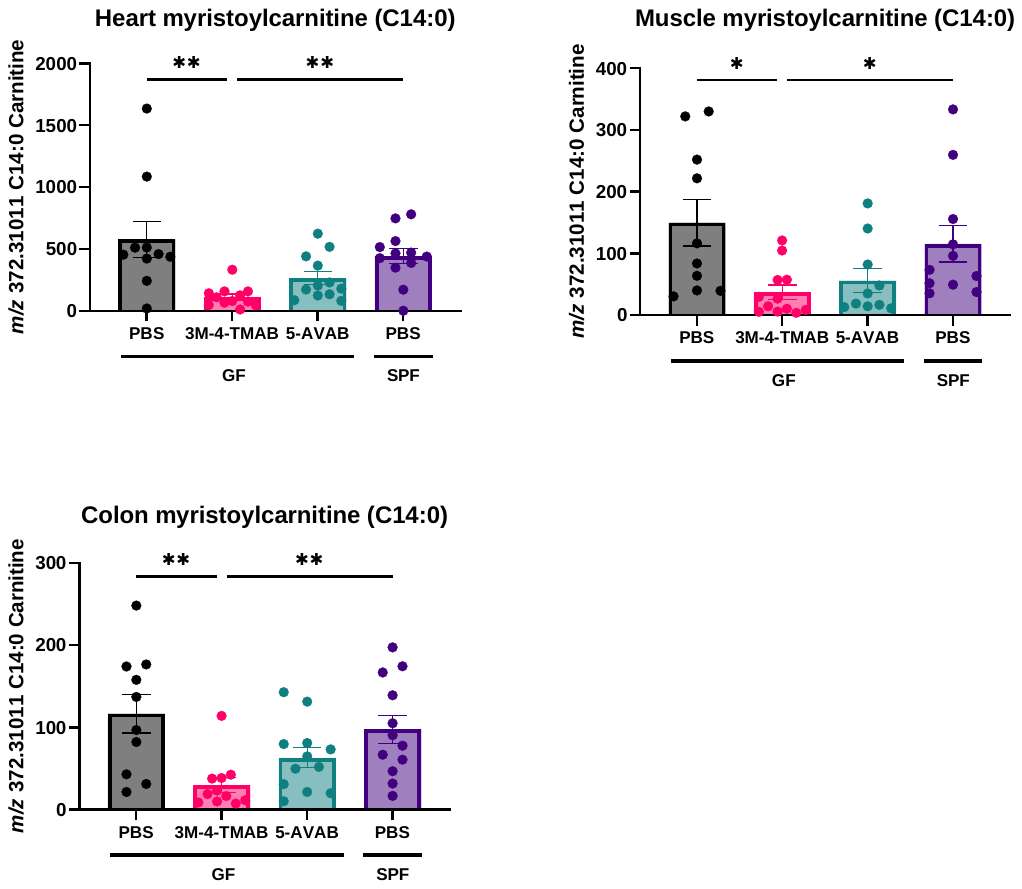

**Figure S5: Relative abundance of myristoylcarnitine (C14:0) across sections from organs from C57BL/6 GF mice treated with metabolites or PBS and C57BL/6 SPF mice treated with PBS.** Tissues were isolated from mice, cryosectioned and imaged via DESI-MSI for the presence of myristoylcarnitine. GF mice were treated with PBS (control) or 100 mg/Kg metabolite (3M-4-TMAB or 5-AVAB) for 5 days via i.p. injection. SPF control mice were treated with PBS. Data are presented as mean ± SEM. Data were analyzed using the Kruskal–Wallis test followed by Dunn's multiple comparisons test. Asterisks denote significant differences with * = *p*<0.05, ** = *p*<0.01, *** = *p*<0.001, and **** =*p*<0.0001.

**Figure S6**

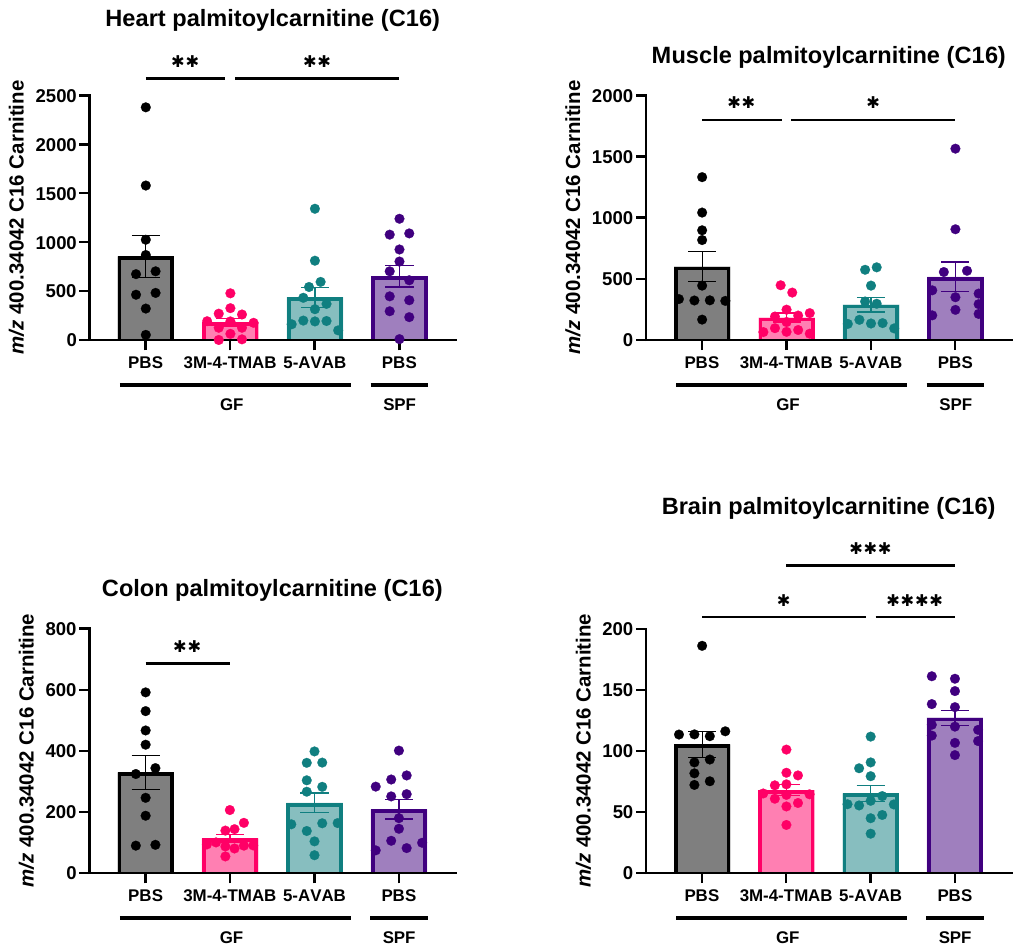

**Figure S6: Relative abundance of palmitoylcarnitine (C16) across sections from organs from C57BL/6 GF mice treated with metabolites or PBS and C57BL/6 SPF mice treated with PBS.** Tissues were isolated from mice, cryosectioned and imaged via DESI-MSI for the presence of palmitoylcarnitine. GF mice were treated with PBS (control) or 100 mg/Kg metabolite (3M-4-TMAB or 5-AVAB) for 5 days via i.p. injection. SPF control mice were treated with PBS. Data are presented as mean ± SEM. Data were analyzed using the Kruskal–Wallis test followed by Dunn's multiple comparisons test. Asterisks denote significant differences with * = *p*<0.05, ** = *p*<0.01, *** = *p*<0.001, and **** =*p*<0.0001.

**Figure S7**

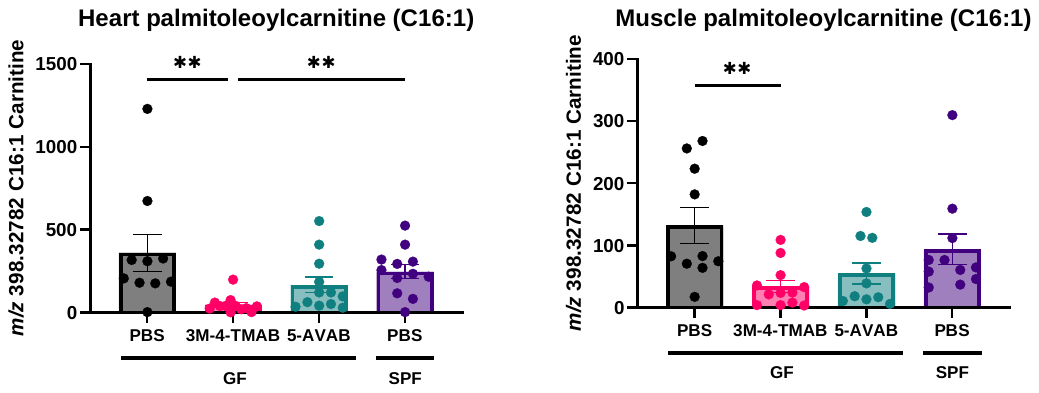

**Figure S7: Relative abundance of palmitoyleoylcarnitine (C16:1) across sections from organs from C57BL/6 GF mice treated with metabolites or PBS and C57BL/6 SPF mice treated with PBS.** Tissues were isolated from mice, cryosectioned and imaged via DESI-MSI for the presence of palmitoyleoylcarnitine. GF mice were treated with PBS (control) or 100 mg/Kg metabolite (3M-4-TMAB or 5-AVAB) for 5 days via i.p. injection. SPF control mice were treated with PBS. Data are presented as mean ± SEM. Data were analyzed using the Kruskal–Wallis test followed by Dunn's multiple comparisons test. Asterisks denote significant differences with ** = *p*<0.01.

**Figure S8**

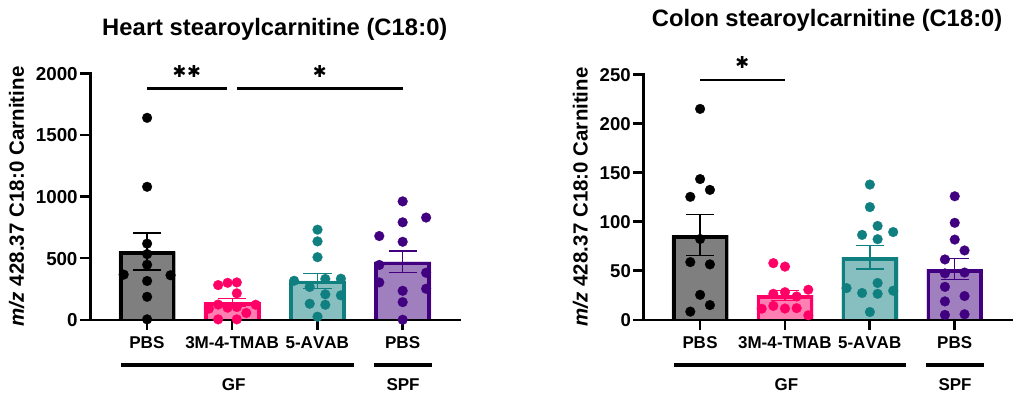

**Figure S8: Relative abundance of stearoylcarnitine (C18:0) across sections from organs from C57BL/6 GF mice treated with metabolites or PBS and C57BL/6 SPF mice treated with PBS.** Tissues were isolated from mice, cryosectioned and imaged via DESI-MSI for the presence of stearoylcarnitine. GF mice were treated with PBS (control) or 100 mg/Kg metabolite (3M-4-TMAB or 5-AVAB) for 5 days via i.p. injection. SPF control mice were treated with PBS. Data are presented as mean ± SEM. Data were analyzed using the Kruskal–Wallis test followed by Dunn's multiple comparisons test. Asterisks denote significant differences with * = *p*<0.05.

**Figure S9**

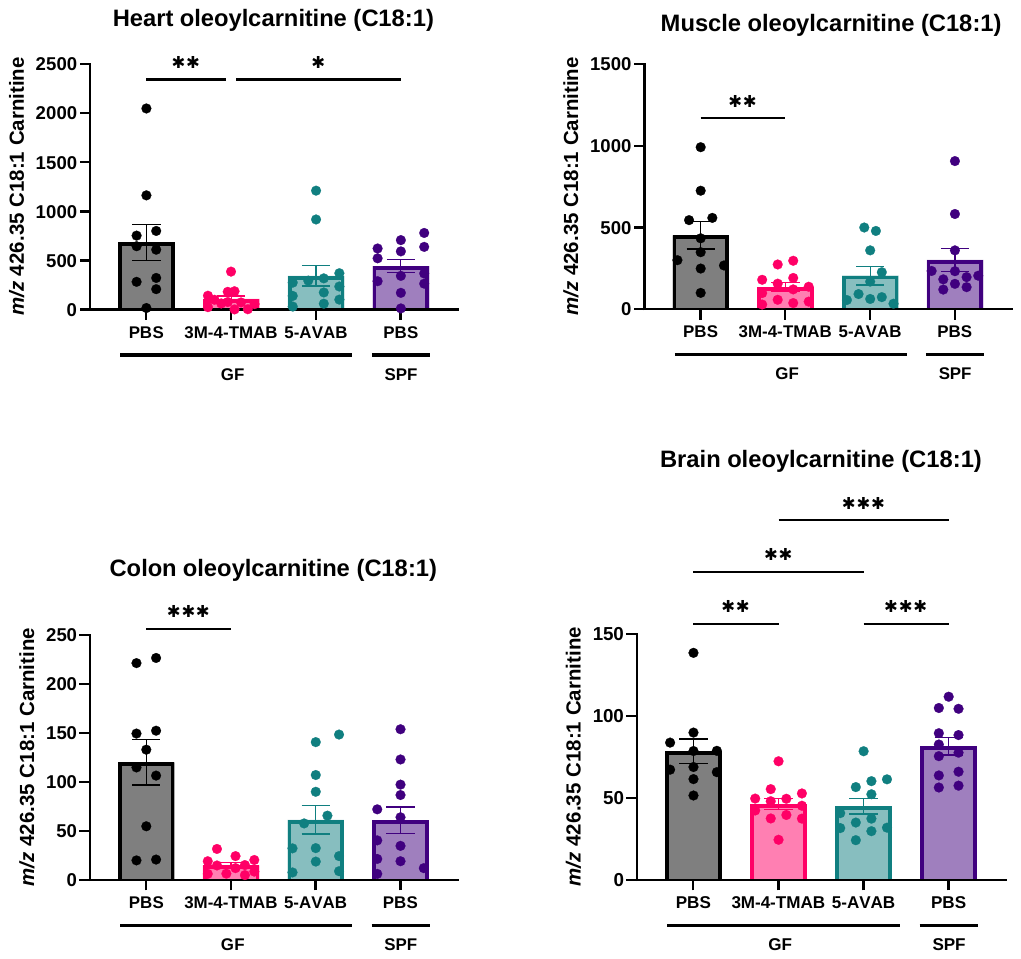

**Figure S9: Relative abundance of oleoylcarnitine (C18:1) across sections from organs from C57BL/6 GF mice treated with metabolites or PBS and C57BL/6 SPF mice treated with PBS.** Tissues were isolated from mice, cryosectioned and imaged via DESI-MSI for the presence of oleoylcarnitine. GF mice were treated with PBS (control) or 100 mg/Kg metabolite (3M-4-TMAB or 5-AVAB) for 5 days via i.p. injection. SPF control mice were treated with PBS. Data are presented as mean ± SEM Data were analyzed using the Kruskal–Wallis test followed by Dunn's multiple comparisons test. Asterisks denote significant differences with * = *p*<0.05, ** = *p*<0.01, *** = *p*<0.001, and **** =*p*<0.0001.

**Figure S10**

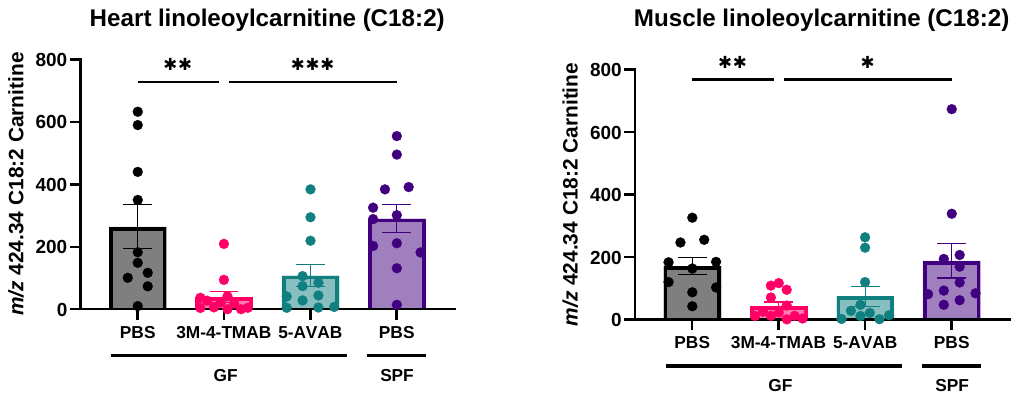

**Figure S10: Relative abundance of linoleoylcarnitine (C18:2) across heart and muscle sections from organs from C57BL/6 GF mice treated with metabolites or PBS and C57BL/6 SPF mice treated with PBS.** Tissues were isolated from mice, cryosectioned and imaged via DESI-MSI for the presence of lineoylcarnitine. GF mice were treated with PBS (control) or 100 mg/Kg metabolite (3M-4-TMAB or 5-AVAB) for 5 days via i.p. injection. SPF control mice were treated with PBS. Data are presented as mean ± SEM. Data were analyzed using the Kruskal–Wallis test followed by Dunn's multiple comparisons test. Asterisks denote significant differences with * = *p*<0.05, ** = *p*<0.01 and *** = *p*<0.001.

**Figure S11**

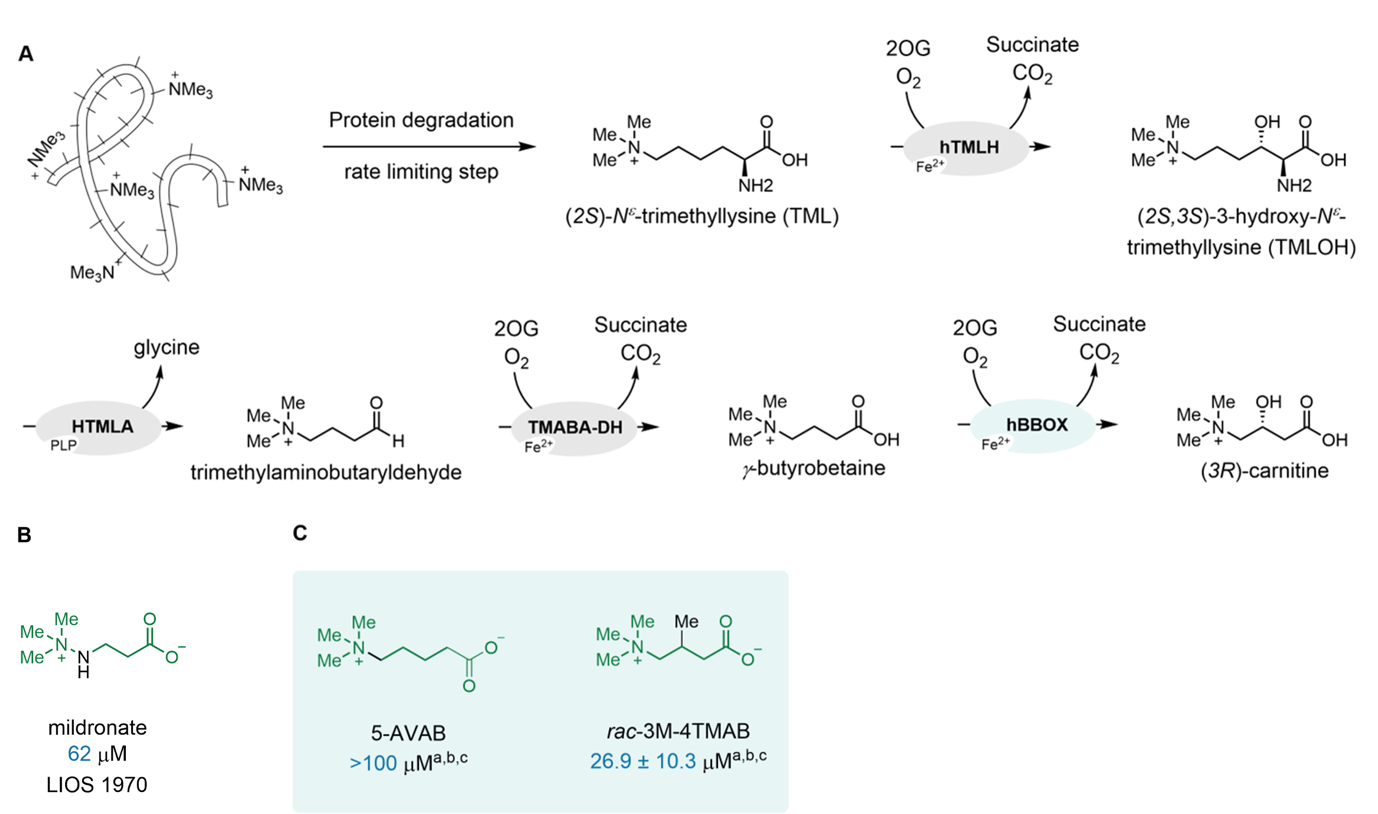

**Figure S11. Carnitine biosynthesis route via protein degradation and eventual conversion of γ-butyrobetaine to carnitine via γ-butyrobetaine hydroxylase (BBOX).** (**A**) Biosynthesis of carnitine TMLH: (2S)-N^ε^-trimethyllysine hydroxylase; HTMLA: (2S,3S)-3-hydroxy-N^ε^-trimethyllysine aldolase; TMABA-DH: trimethylaminobutaryldehyde dehydrogenase; BBOX: γ-butyrobetaine hydroxylase. (**B**) IC_50_ of mildronate a BBOX inhibitor that mimics γ-butyrobetaine and (**C**), Inhibition of BBOX by 5-AVAB and 3M-4TMAB. Mean average of two independent runs, each composed of technical duplicates (n = 2; mean ± SD). IC_50_ values obtained using a SPE-MS assay using 0.05 μM human BBOX, 400 μM 2-oxoglutarate and 25 μM GBB. Note it is possible that in addition to BBOX, 5-AVAB and 3M-4TMAB inhibit earlier steps in carnitine biosynthesis and/or impact on carnitine uptake / export.

**Figure S12**

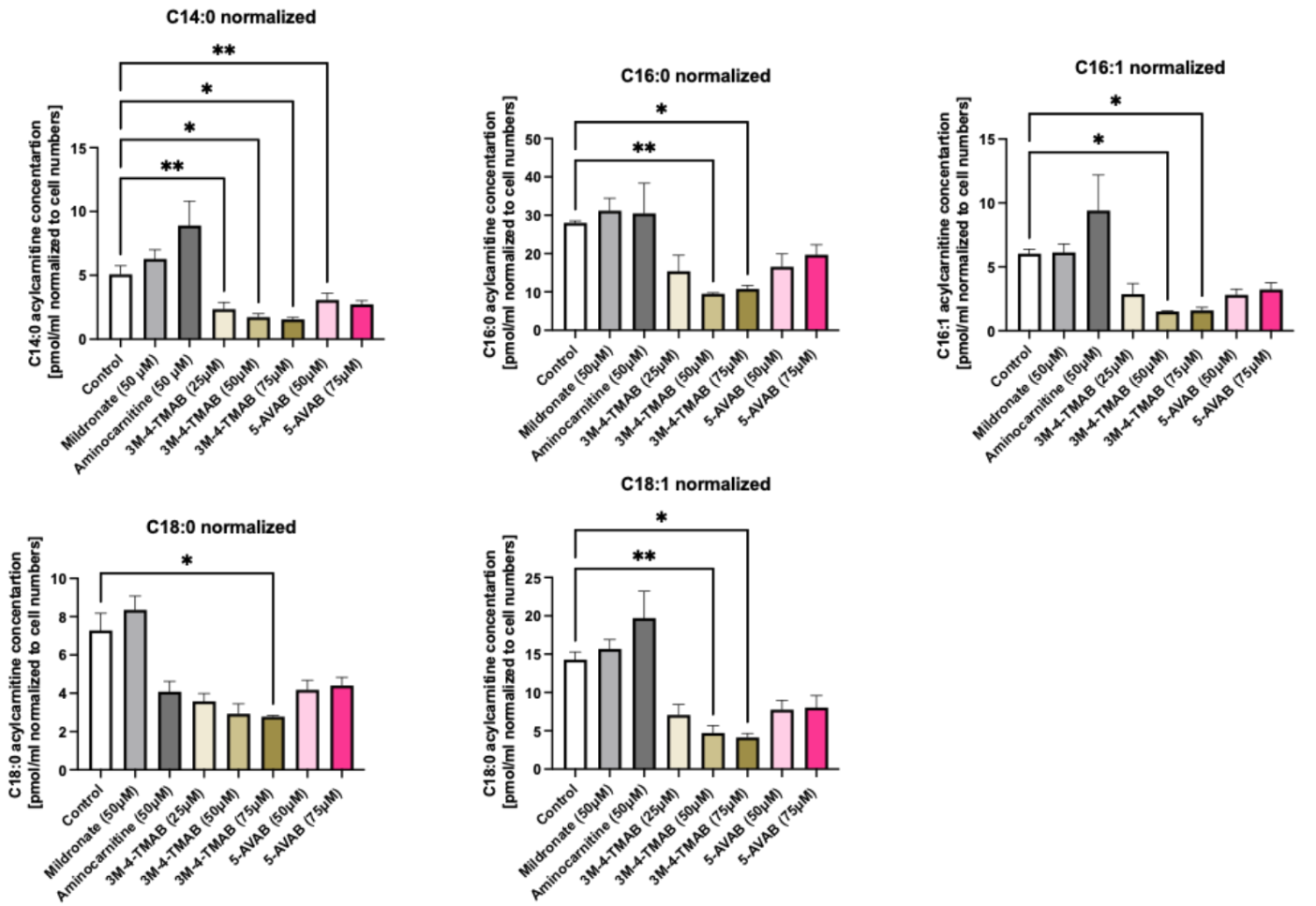

**Figure S12: Acylcarnitine profiling of HepG2 liver cells post-treatment with 3M-4-TMAB, 5-AVAB and control FAO inhibitors.** HepG2 cells were treated with increasing concentrations of the metabolites of interest and the known FAO inhibitors aminocarnitine and mildronate for 48 hours before lipids were extracted and analyzed by LC-MS alongside internal standards. Data are displayed as mean ± SEM and show 3 biological replicates. Statistical significance was determined by a one-way ANOVA followed by Dunnett’s multiple comparison test. Asterisks denote significant differences with * = *p*<0.05 and ** = *p*<0.01.

**Figure S13**

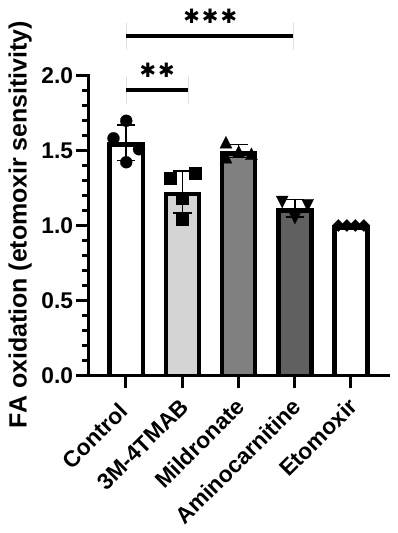

**Figure S13: FAO inhibition relative to that of known irreversible FAO inhibitor etomoxir.** HepG2 cells were treated with 75 μM of 3M-4-TMAB, 4-TMAP, mildronate or aminocarnitine for 72 h before a final 35 min incubation with [^3^H]-palmitic acid and 75 μM of the same inhibitor, alongside an etomoxir control. ^3^H_2_0 produced from [^3^H]-palmitate was quantified as an indicator of FAO. Statistical significance was determined by a one-way ANOVA followed by Dunnett’s multiple comparison test. Asterisks denote significant differences with ** = *p*<0.01 and *** = *p*<0.001.

**Figure S14**

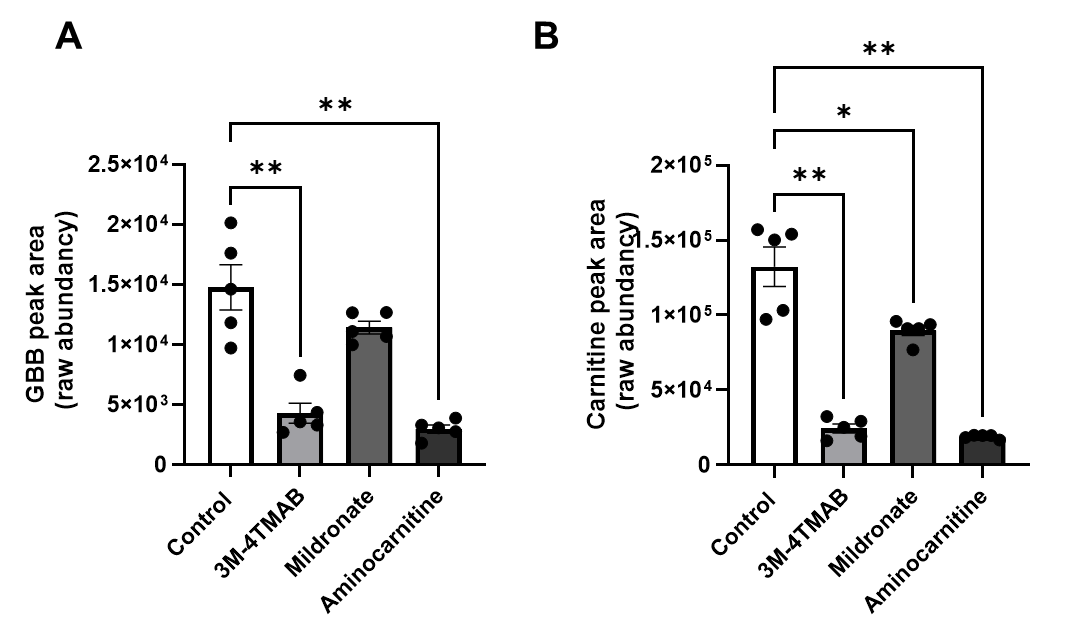

**Figure S14: Carnitine and GBB levels in HepG2 cells in the presence of inhibitor.** HepG2 cells were treated with 75 μM 3M-4-TMAB, 4-TMAP, mildronate or aminocarnitine for 72 h and levels of (**A**) GBB and (**B**) carnitine measured via LCMS. Statistical significance was determined by a one-way ANOVA followed by Dunnett’s multiple comparison test. Asterisks denote significant differences with * = *p*<0.05 and ** = *p*<0.01.

**Figure S15**

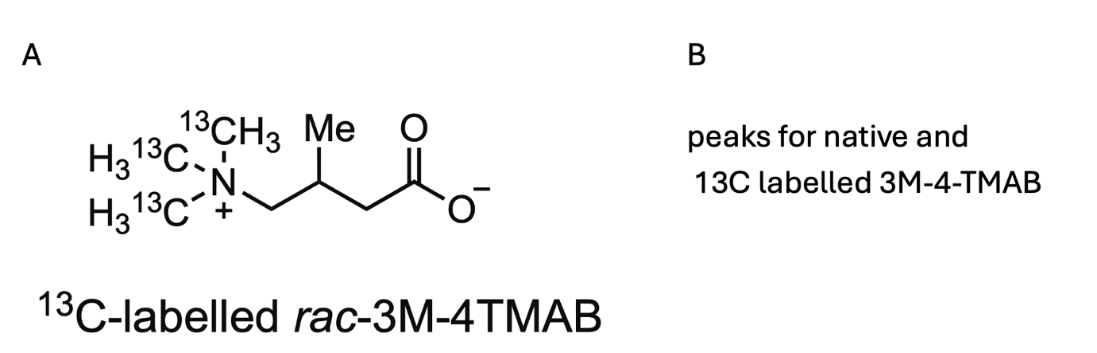

**Figure S15: Structure of ^13^C-labelled 3M-4-TMAB.** ^13^C-3M-4-TMAB structure showing the location of the labelled carbon atoms in the amine group. NMR data for labelled 3M-4-TMAB is shown in Figure S28.

**Figure S16**

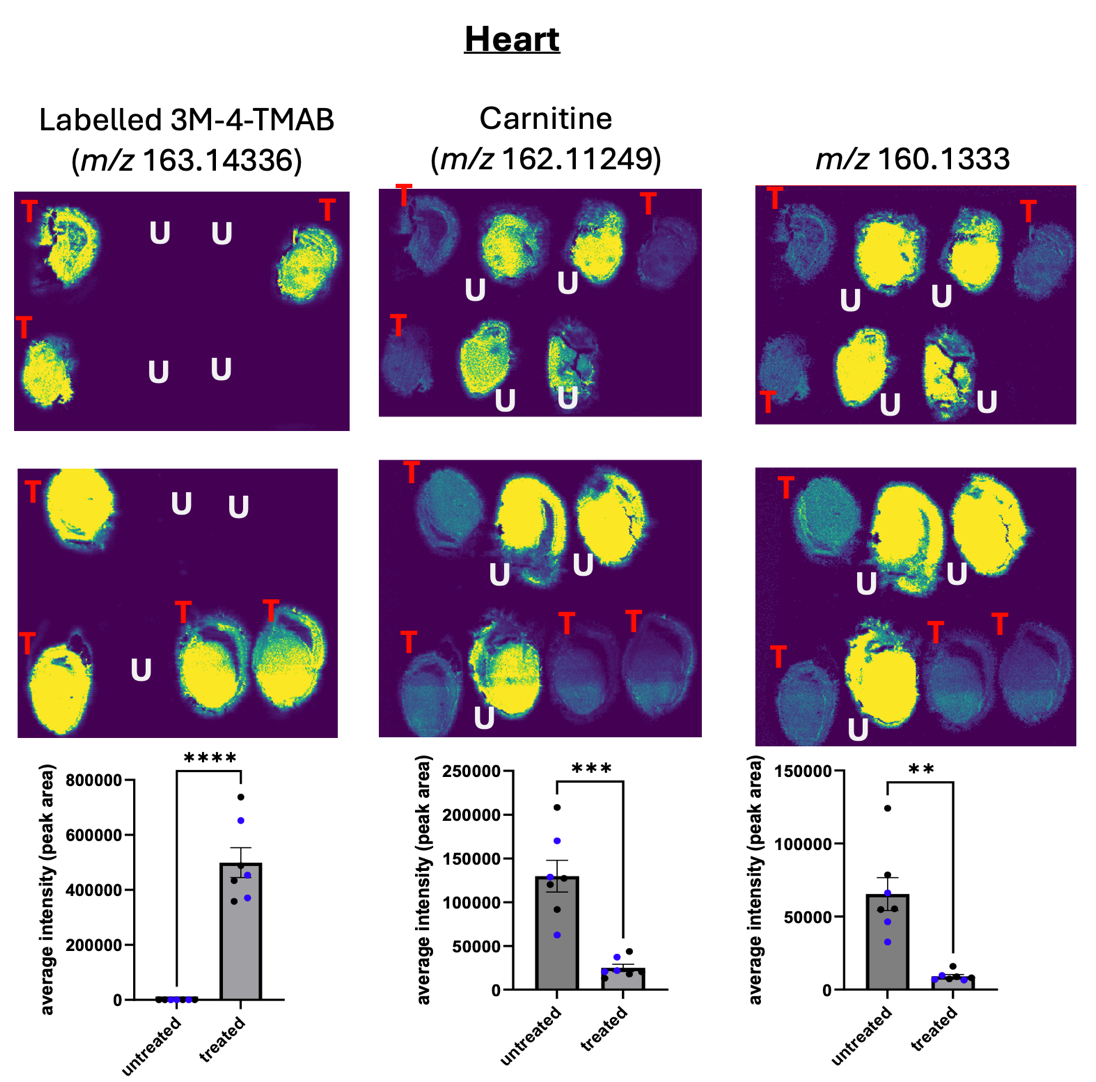

**Figure S16: DESI-MSI images of heart tissue from C57BL/6 SPF mice post-treatment with ^13^C-3M-4-TMAB.** After 5 days of treatment of animals with 100 mg/Kg of ^13^C-3M-4-TMAB hearts were isolated, cryosectioned and subjected to DESI-MSI to determine levels of ^13^C-labelled 3M-4-TMAB, *m/z* 160.133 levels and carnitine levels. Relative intensities are shown beneath each set of images. Blue dots represent male animals and black dots represent female animals.

**Figure S17**

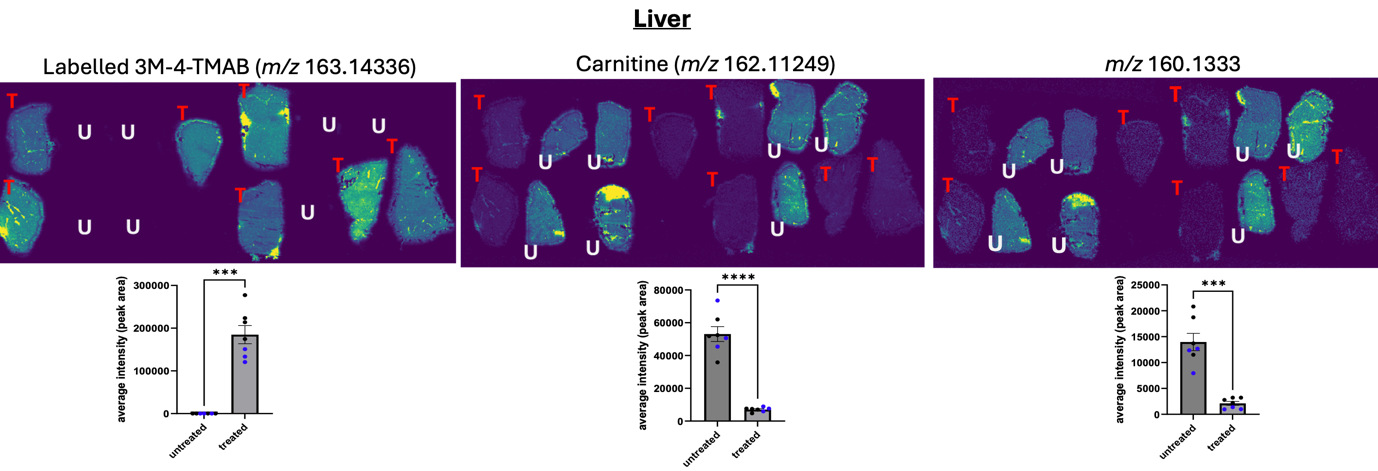

**Figure S17: DESI-MSI images of liver tissue from C57BL/6 SPF mice post-treatment with ^13^C-3M-4-TMAB.** After 5 days of treatment of animals with 100 mg/Kg of ^13^C-3M-4-TMAB livers were isolated, cryosectioned and subjected to DESI-MSI to determine levels of ^13^C-labelled 3M-4-TMAB, *m/z* 160.133 levels and carnitine levels. Relative intensities are shown beneath each set of images. Blue dots represent male animals and black dots represent female animals.

**Figure S18**

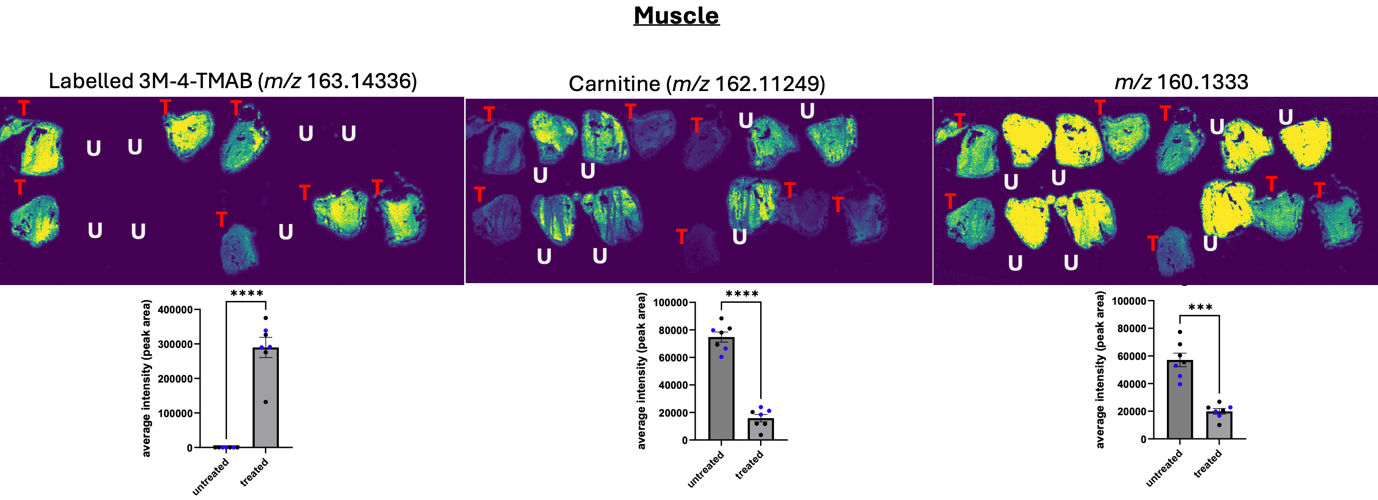

**Figure S18: DESI-MSI images of muscle tissue from C57BL/6 SPF mice post-treatment with ^13^C-3M-4-TMAB.** After 5 days of treatment of animals with 100 mg/Kg of ^13^C-3M-4-TMAB, the muscle tissue was isolated, cryosectioned and subjected to DESI-MSI to determine levels of ^13^C-labelled 3M-4-TMAB, *m/z* 160.133 levels and carnitine levels. Relative intensities are shown beneath each set of images. Blue dots represent male animals and black dots represent female animals.

**Figure S19**

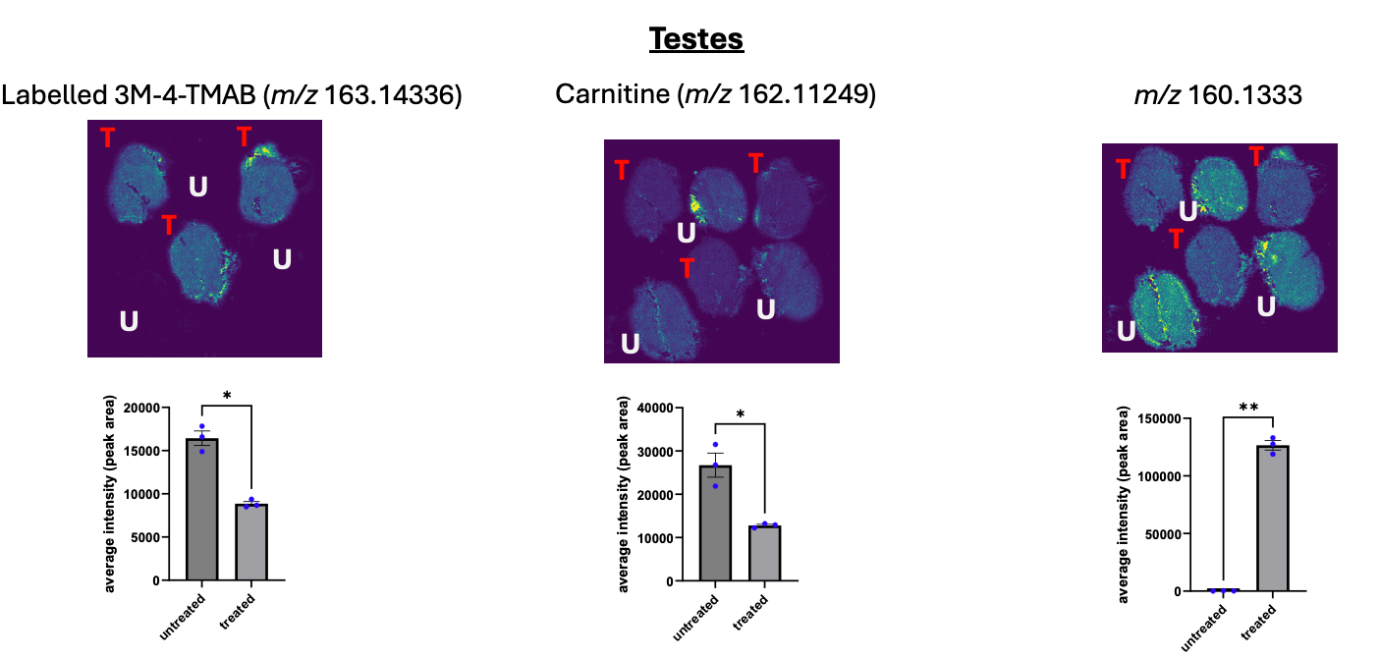

**Figure S19: DESI-MSI images of testes from C57BL/6 SPF mice post-treatment with ^13^C-3M-4-TMAB.** After 5 days of treatment of animals with 100 mg/Kg of ^13^C-3M-4-TMAB testes were removed, cryosectioned and subjected to DESI-MSI to determine levels of ^13^C-labelled 3M-4-TMAB, *m/z* 160.133 levels and carnitine levels. Relative intensities are shown beneath each set of images.

**Figure S20**

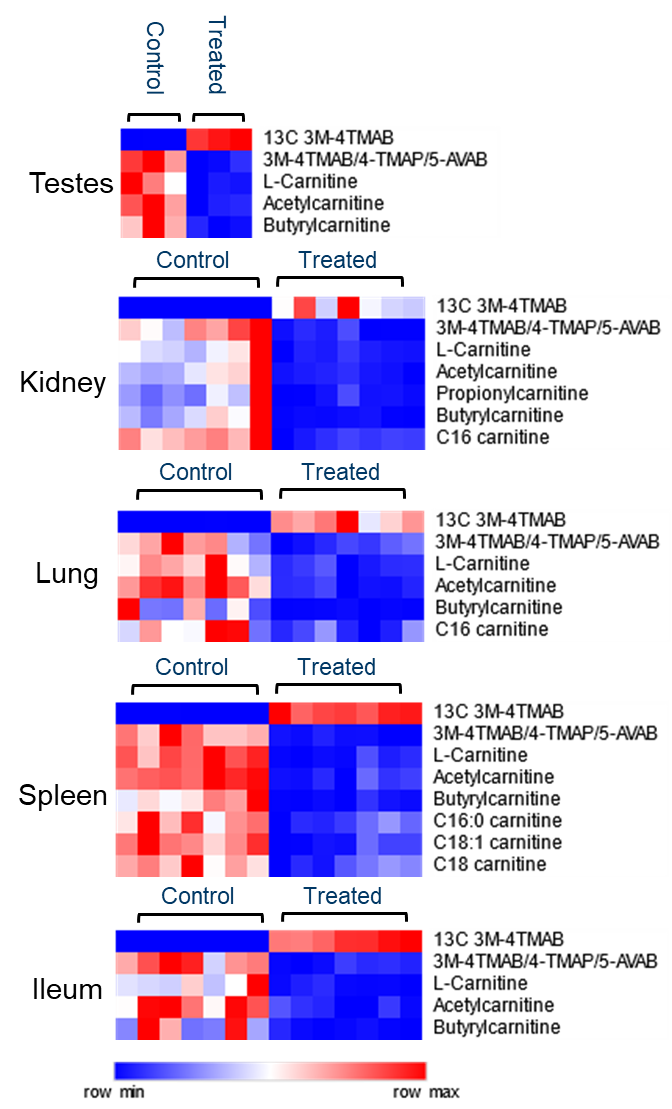

**Figure S20: ^13^C-3M-4-TMAB, carnitine and acylcarnitine levels in tissues of C57BL/6 SPF mice post-treatment with ^13^C-3M-4-TMAB.** After 5 days of treatment of animals with 100 mg/Kg of ^13^C-3M-4-TMAB, tissues were isolated, cryosectioned and subjected to DESI-MSI. Heat maps are shown with relative abundance of ^13^C-3M-4-TMAB and significantly altered carnitine and acylcarnitines across tissues in C57BL/6 SPF mice post-treatment.

**Figure S21 ^1^H and ^13^C spectra of synthesised compounds**

**Methyl 3-methyl-4-nitrobutanoate (2)**

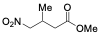

^1^H NMR (400 MHz, 300 K, CDCl_3_):

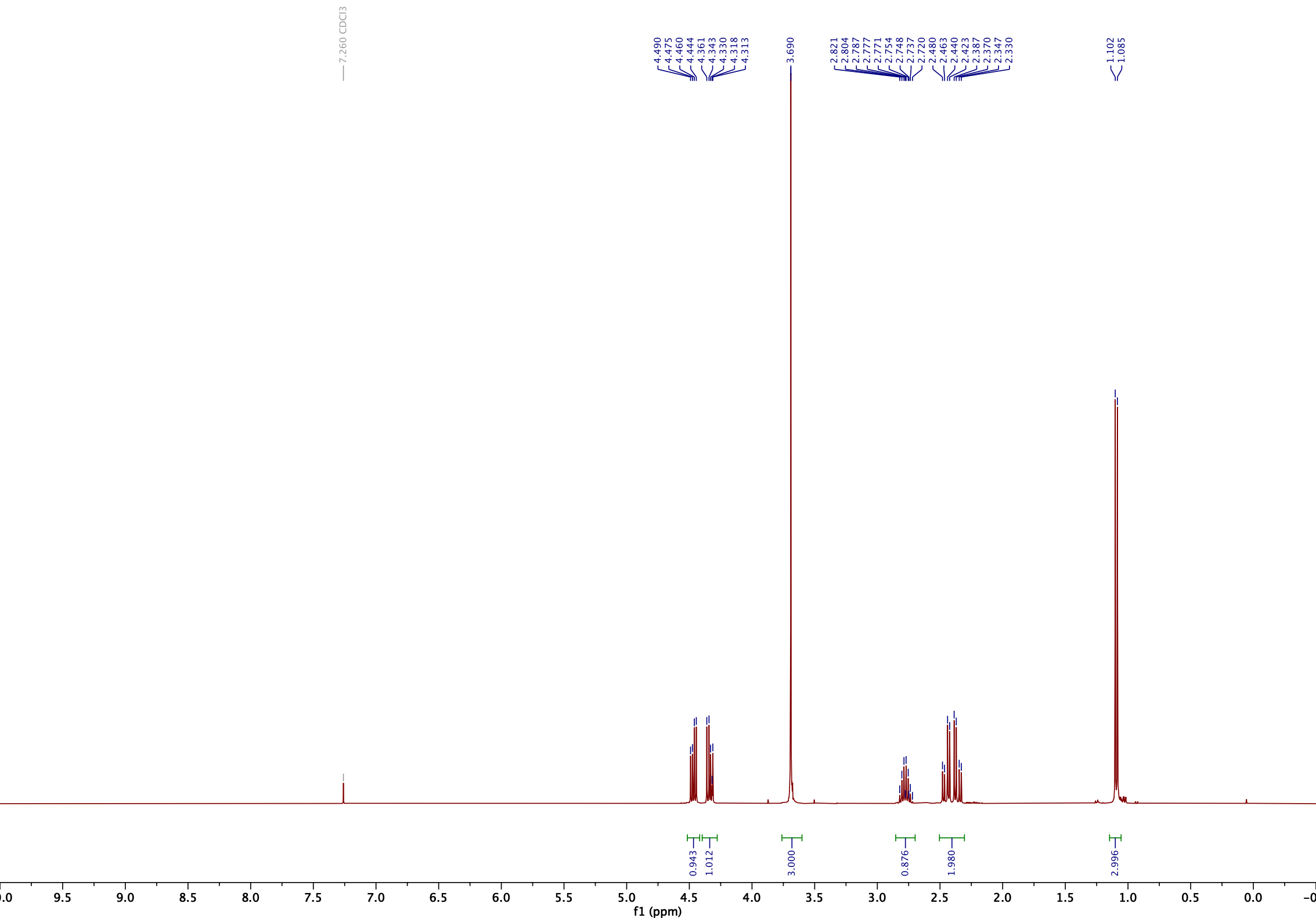

^13^C NMR (101 MHz, 300 K, CDCl_3_):

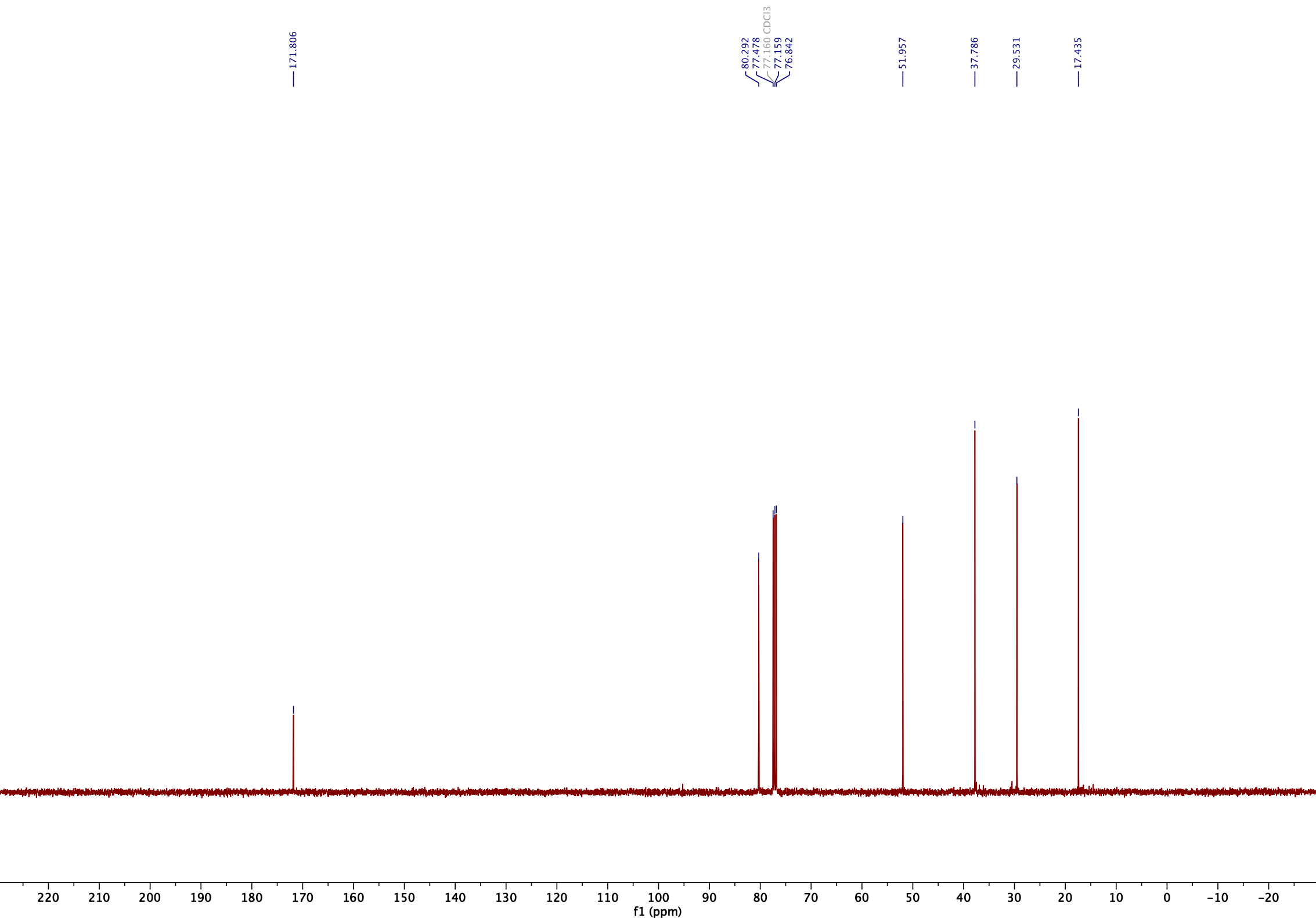

**Figure S22**

**3-Methyl-4-nitrobutanoic acid (2)**

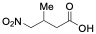

^1^H NMR (400 MHz, 300 K, CDCl_3_):

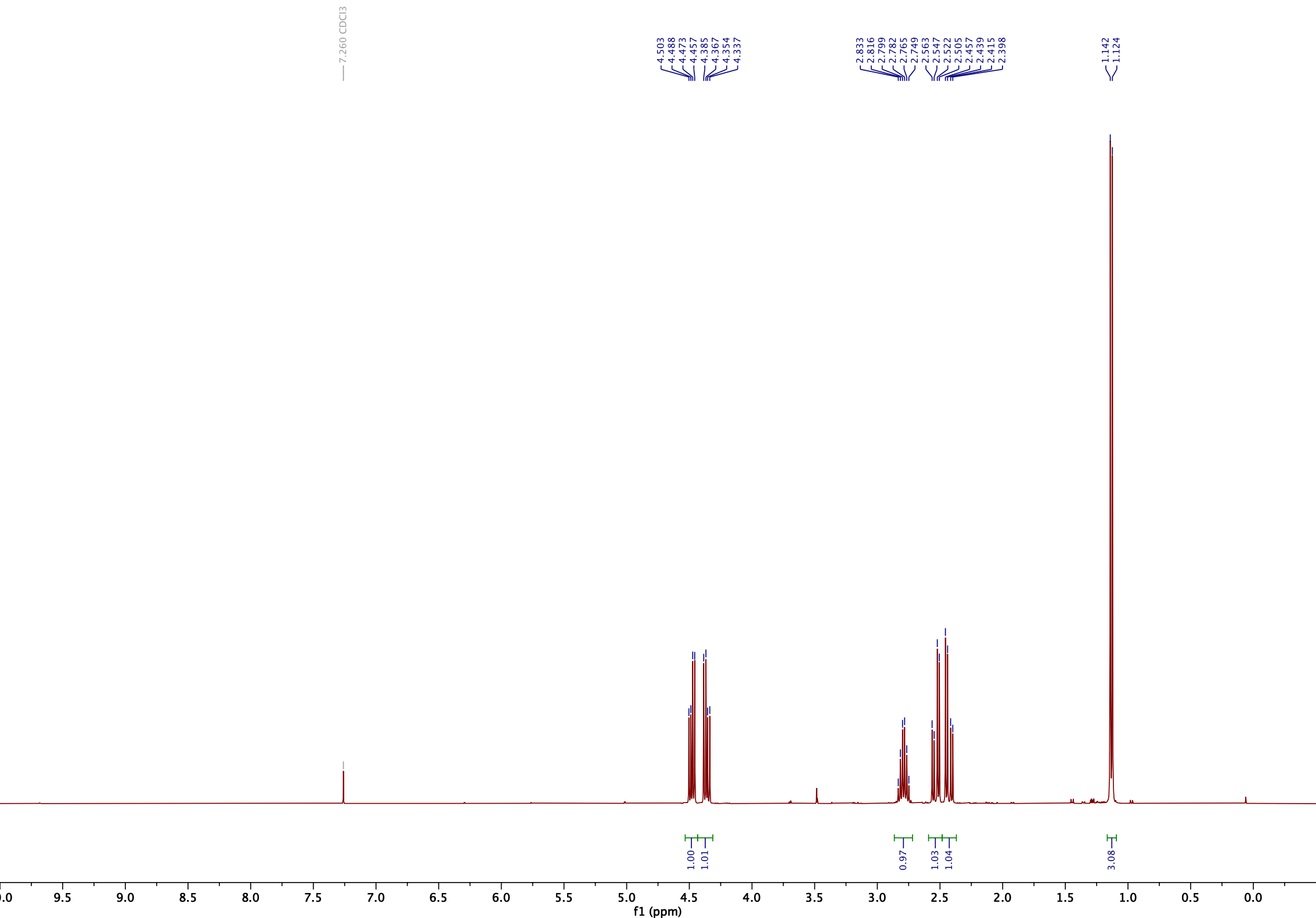

^13^C NMR (101 MHz, 300 K, CDCl_3_):

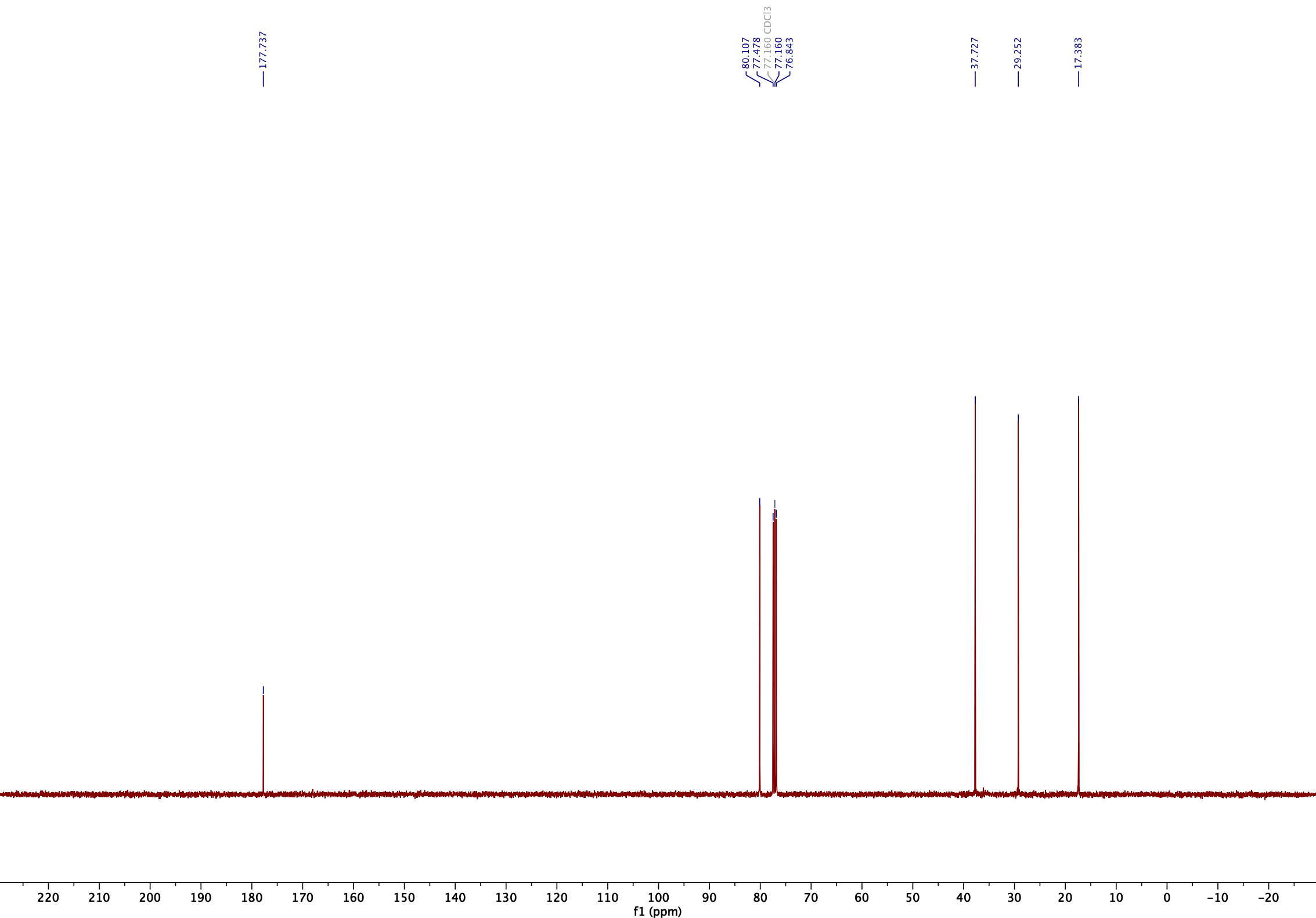

**Figure S23**

**4-Amino-3-methylbutanoic acid (3)**

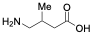

^1^H NMR (400 MHz, 300 K, D_2_O):

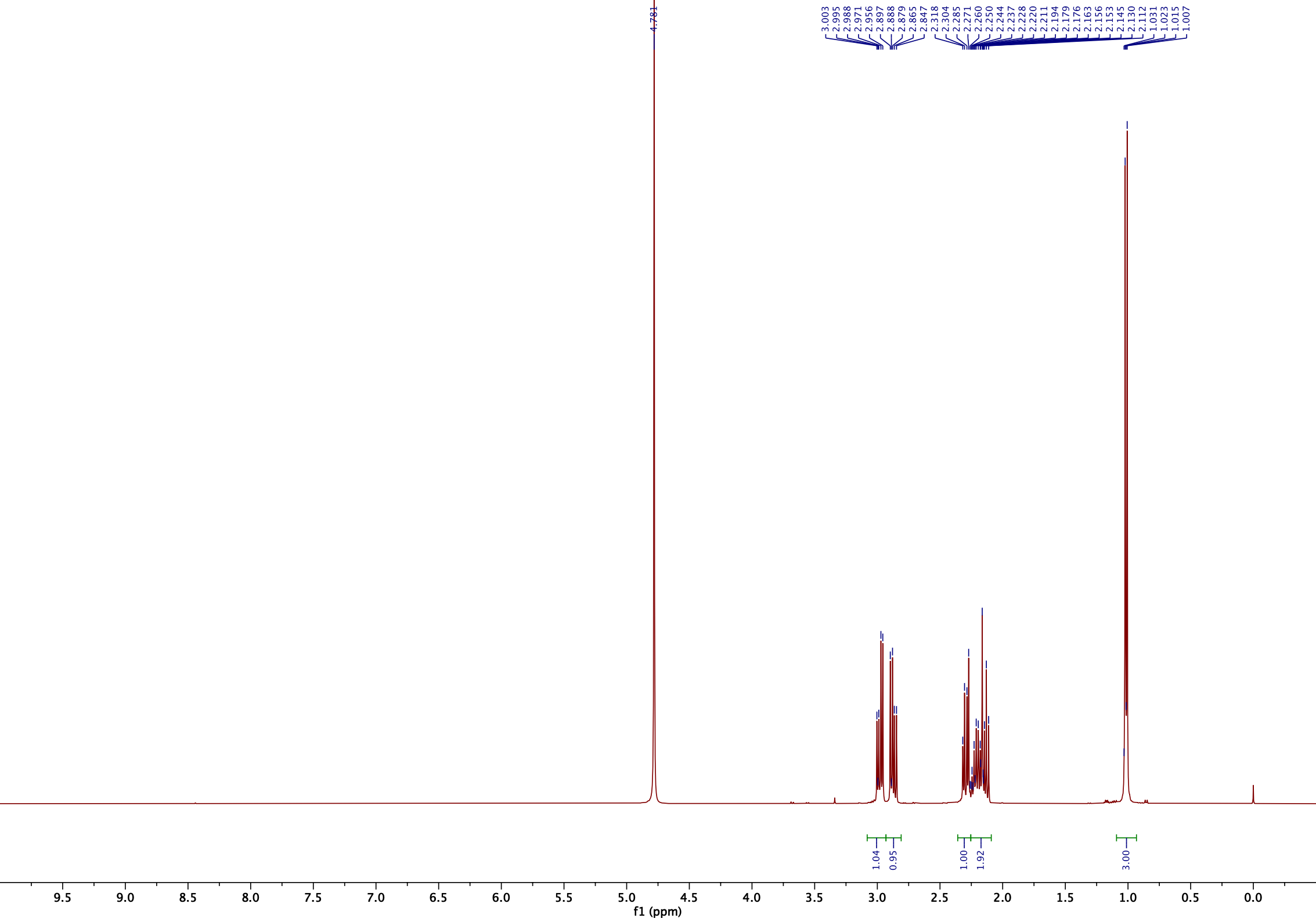

^13^C NMR (101 MHz, 300 K, D_2_O):

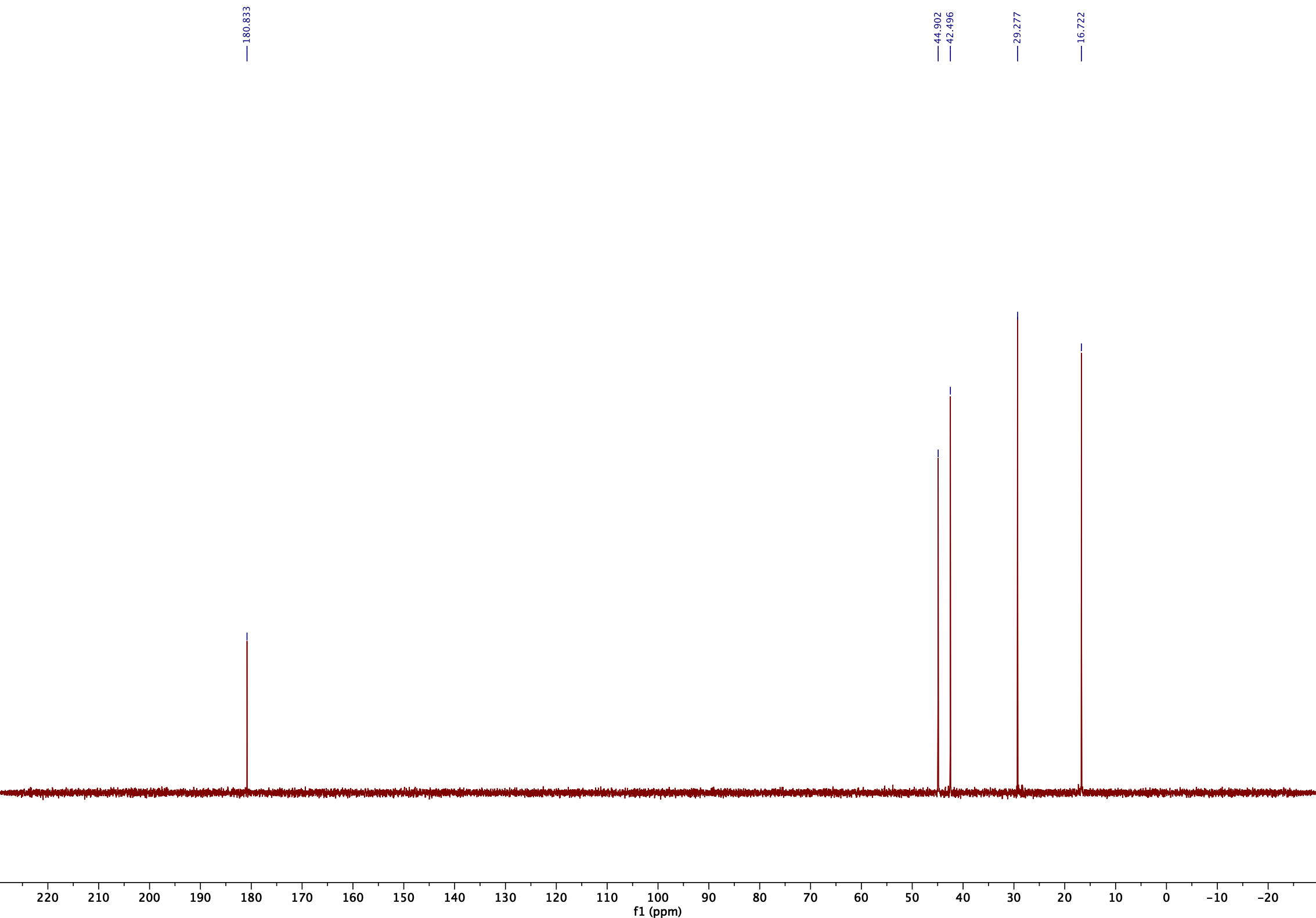

**Figure S24**

**4-Methoxy-*N*,*N*,*N*,2-tetramethyl-4-oxobutan-1-aminium iodide (4)**

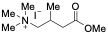

^1^H NMR (400 MHz, 300 K, D_2_O):

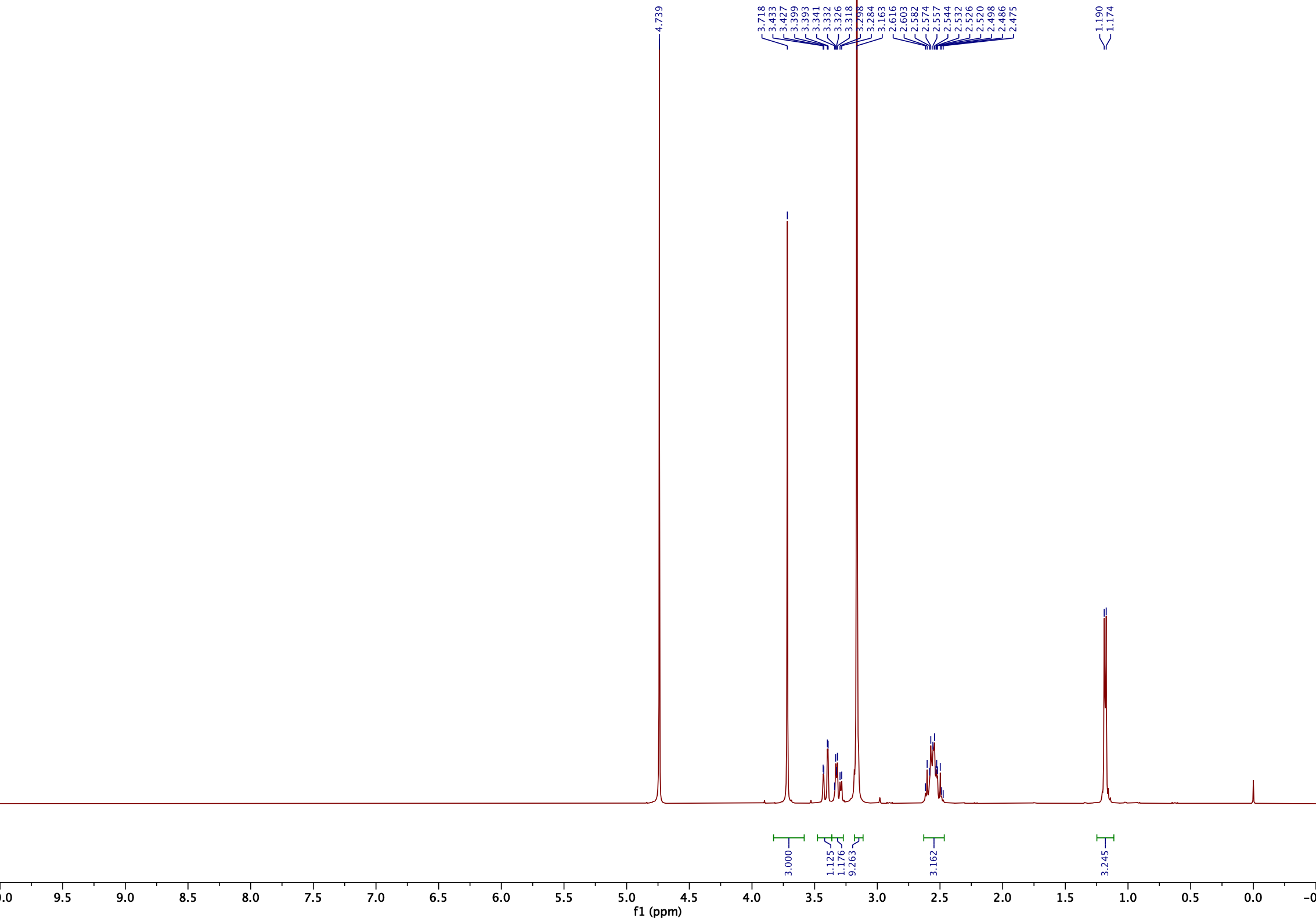

^13^C NMR (101 MHz, 300 K, D_2_O):

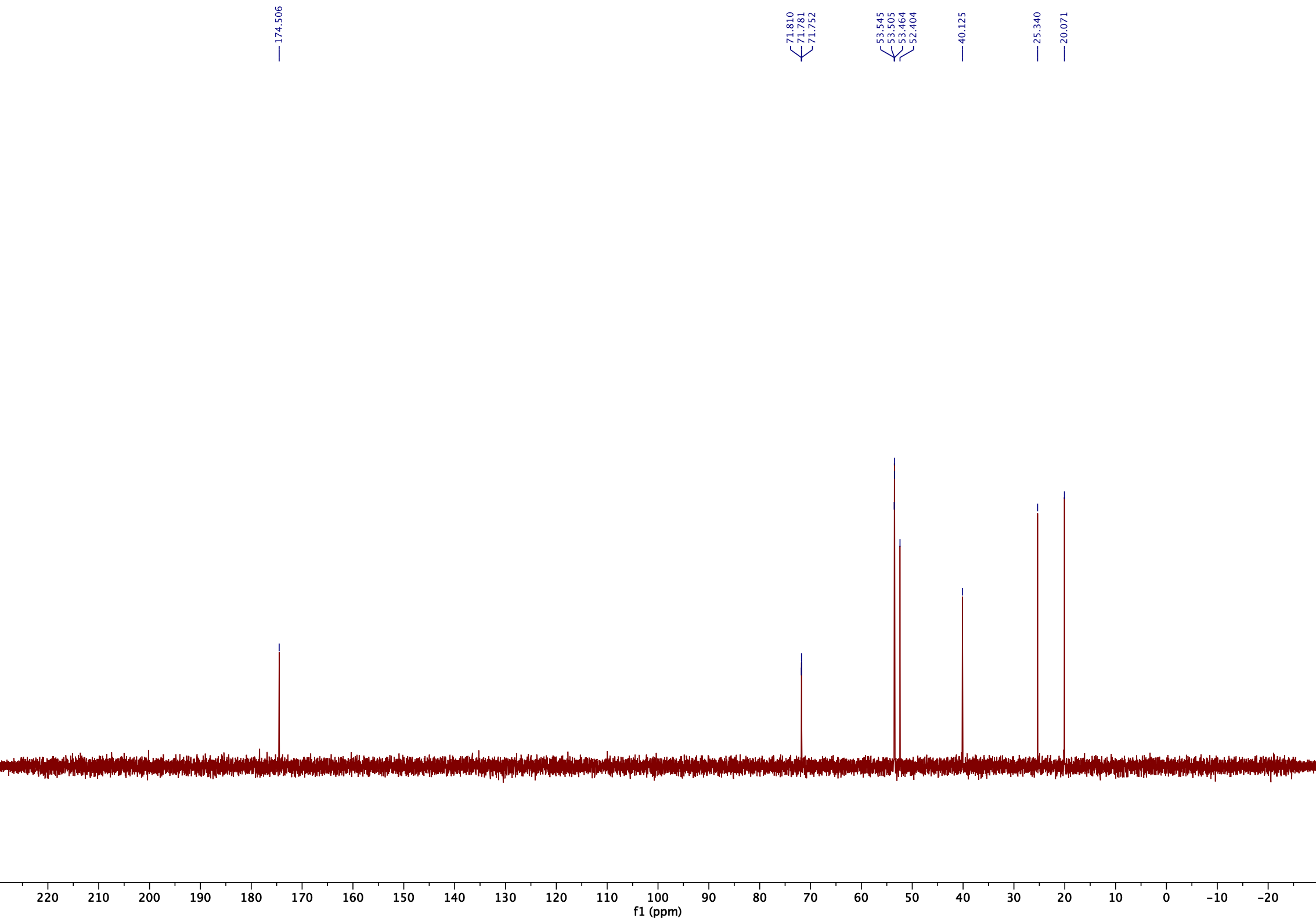

**Figure S25**

**3-Methyl-4-(trimethylammonio)butanoate (3M-4TMAB)**

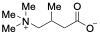

^1^H NMR (600 MHz, 300 K, D_2_O, TSP):

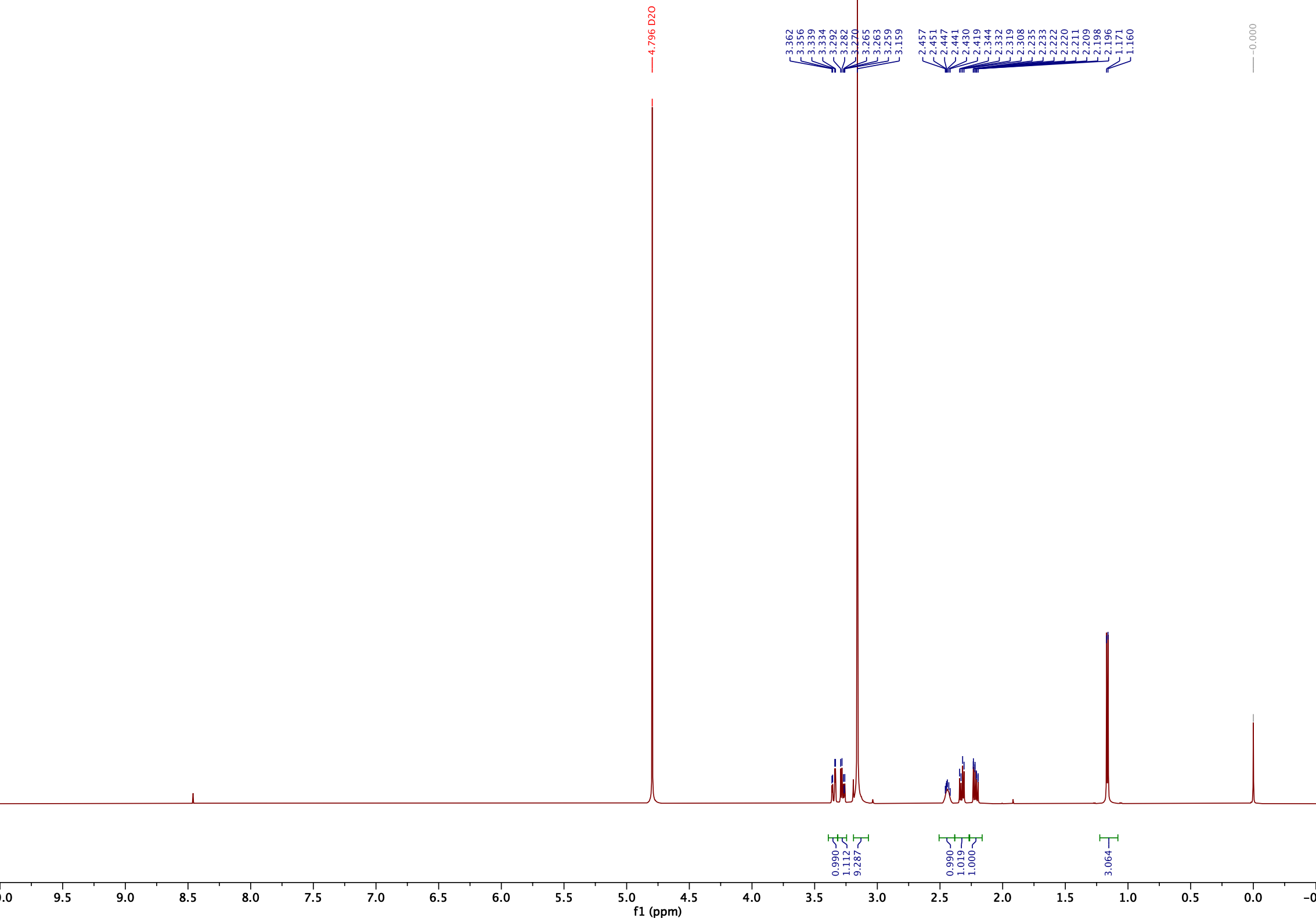

^13^C NMR (151 MHz, 300 K, D_2_O, TSP):

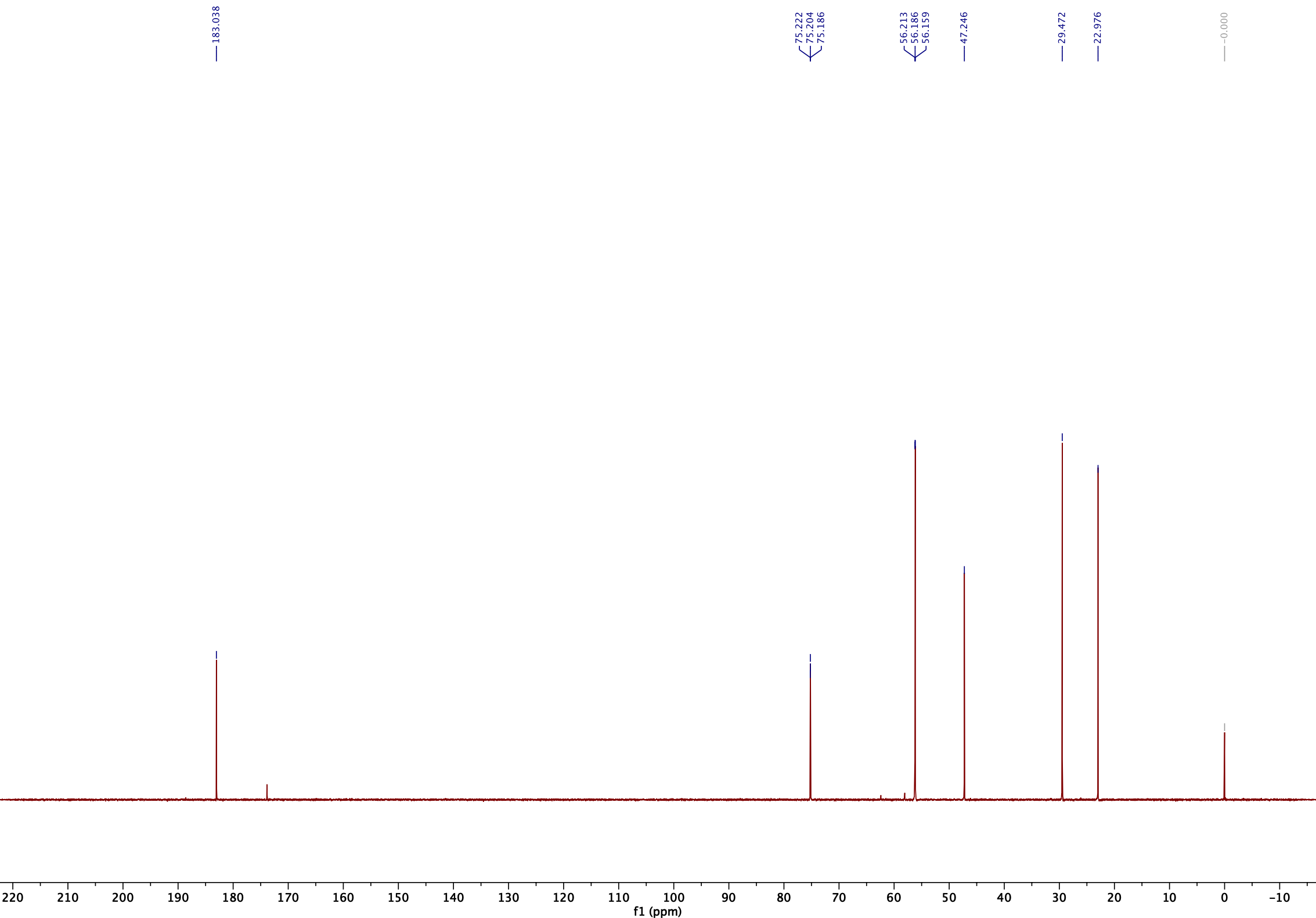

**Figure S26**

**4-Methoxy-2-methyl-4-oxobutan-1-aminium chloride (5)**

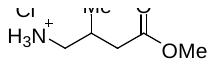

^1^H NMR (600 MHz, 300 K, D_2_O, DSS):

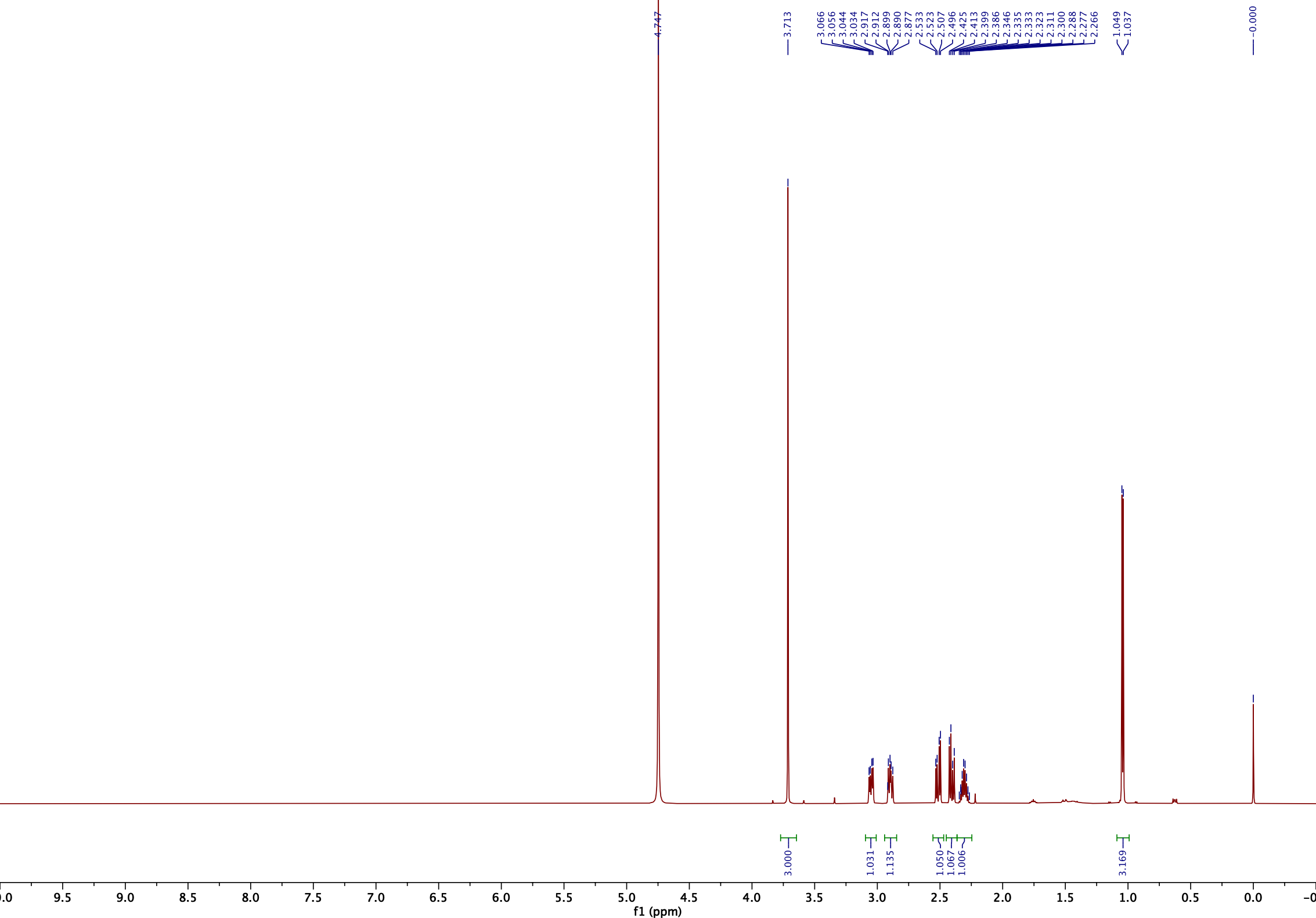

^13^C NMR (151 MHz, 300 K, D_2_O, DSS):

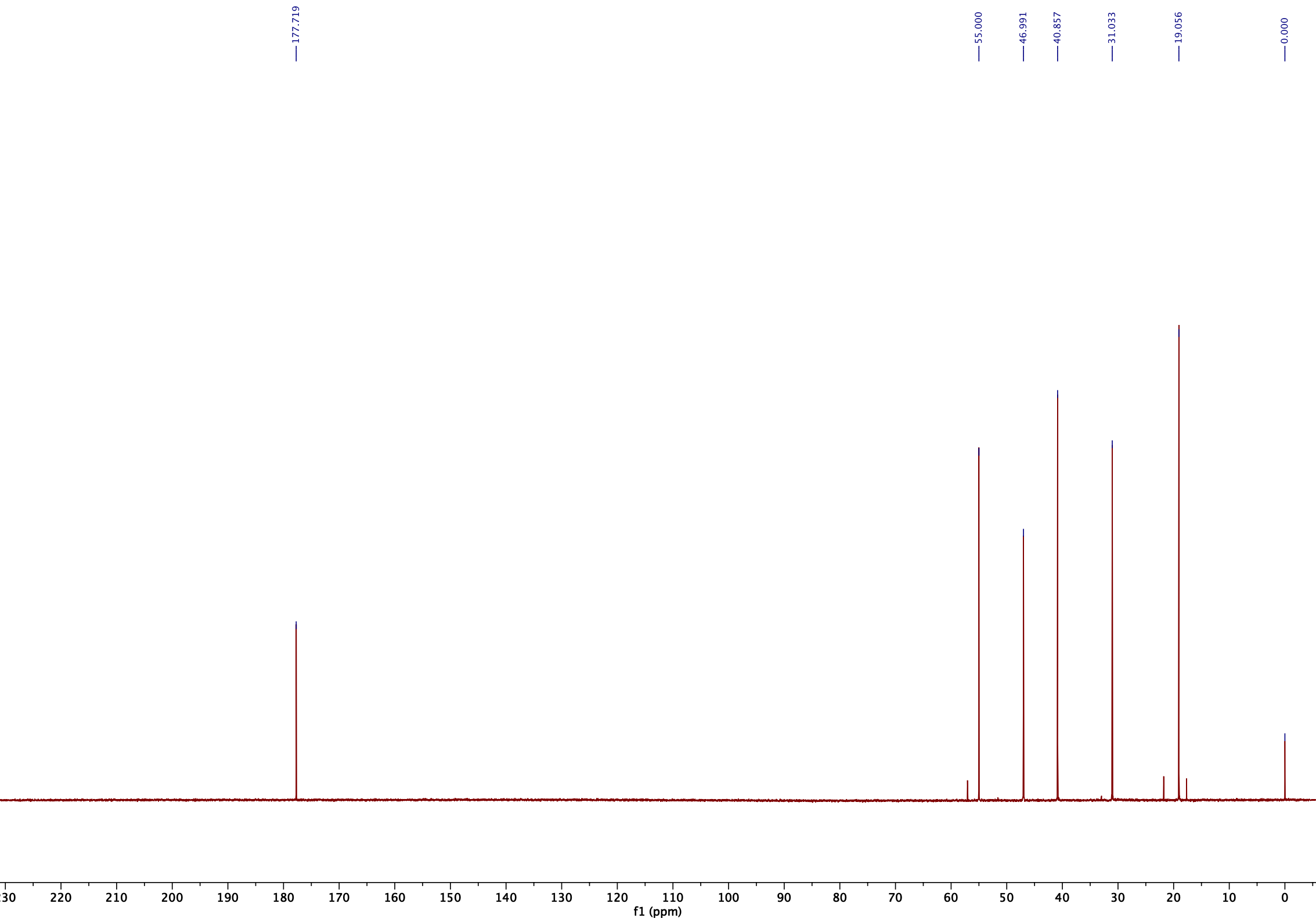

**Figure S27**

**4-Methoxy-2-methyl-*N*,*N*,*N*-tri(methyl-^13^*C*)-4-oxobutan-1-aminium iodide (6)**

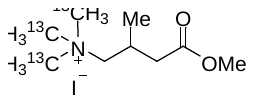

^1^H NMR (500 MHz, 300 K, D_2_O, TSP):

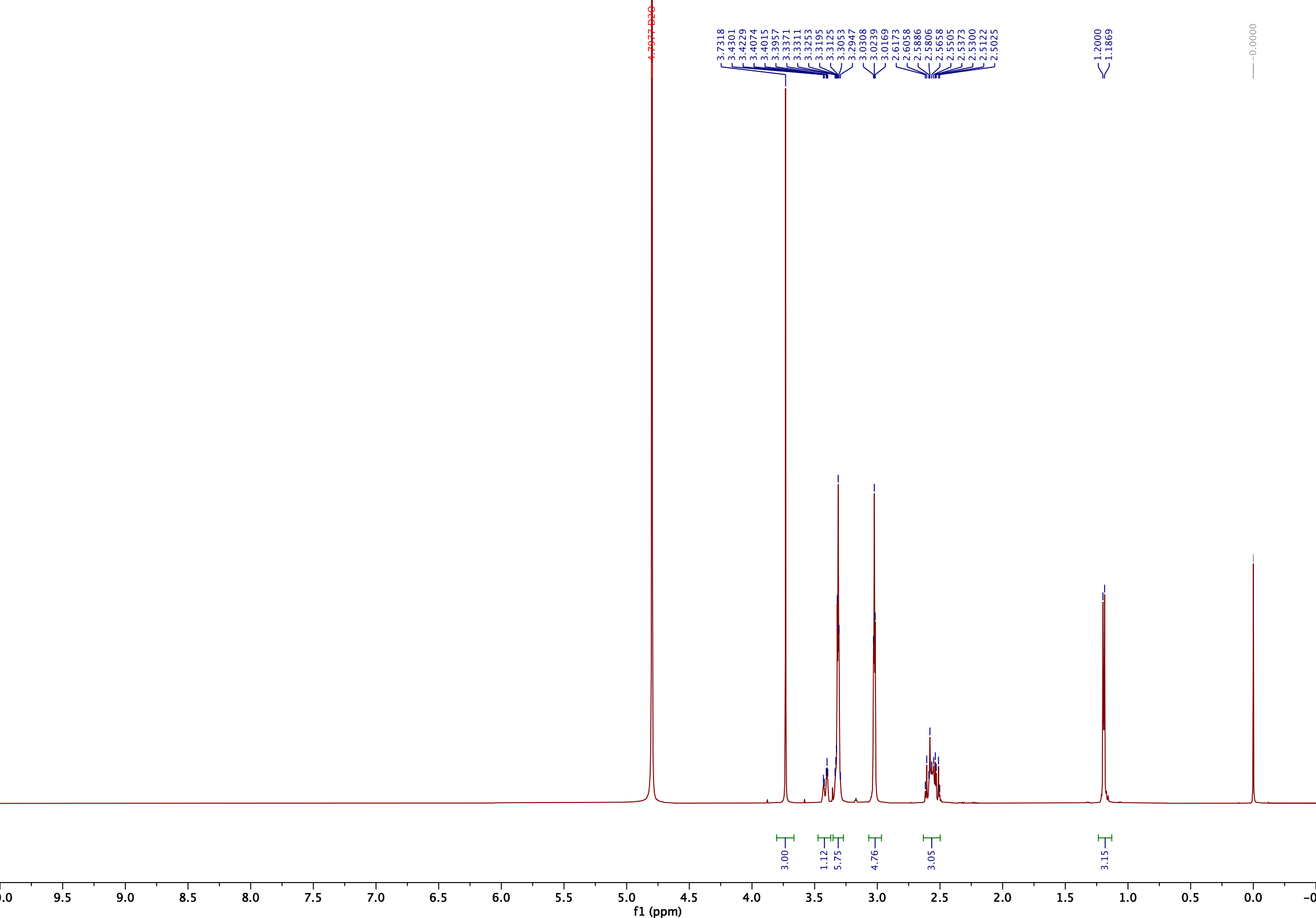

^1^H(^13^C) NMR (500 MHz, 300 K, D_2_O, TSP):

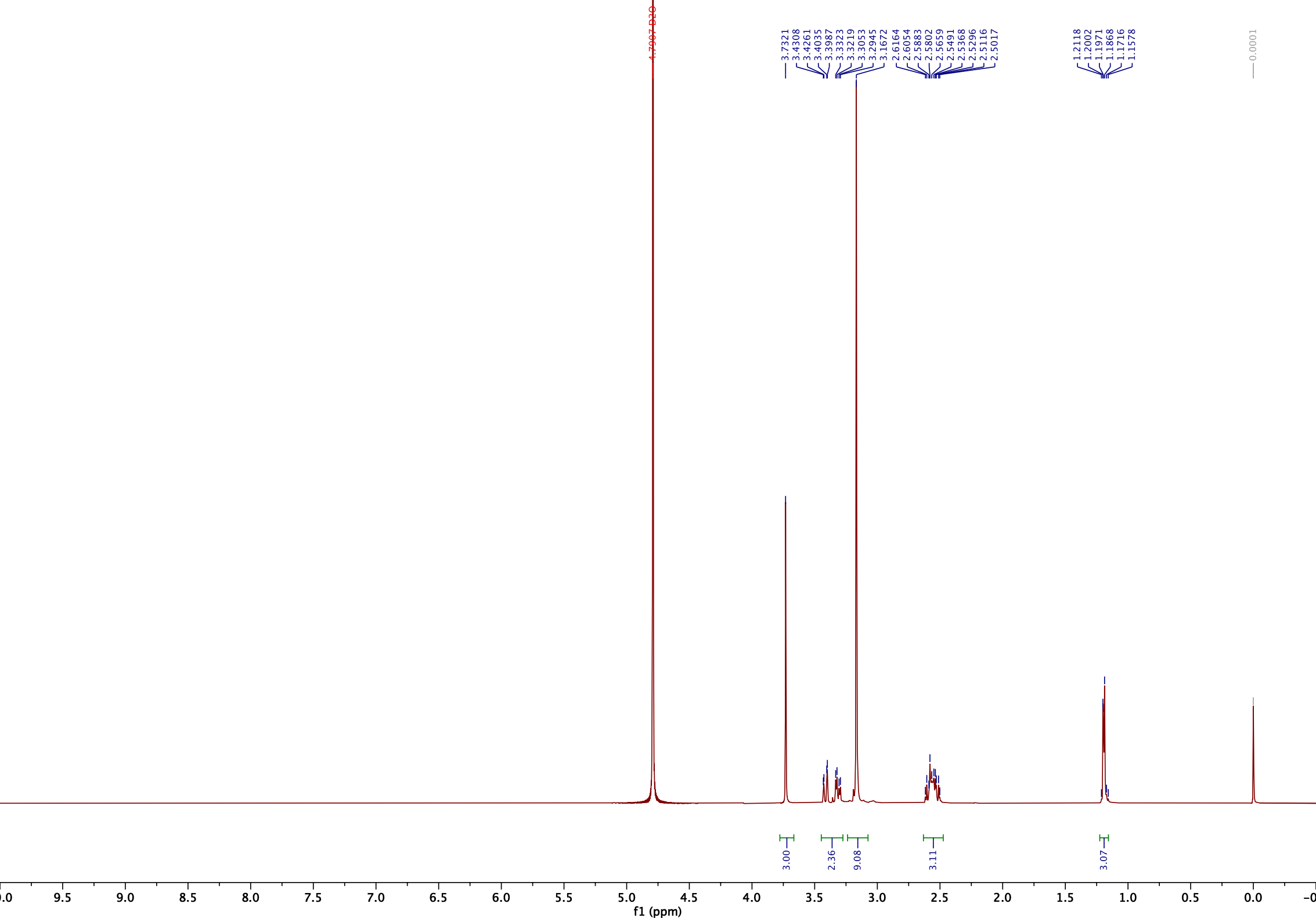

^13^C NMR (151 MHz, 300 K, D_2_O, TSP):

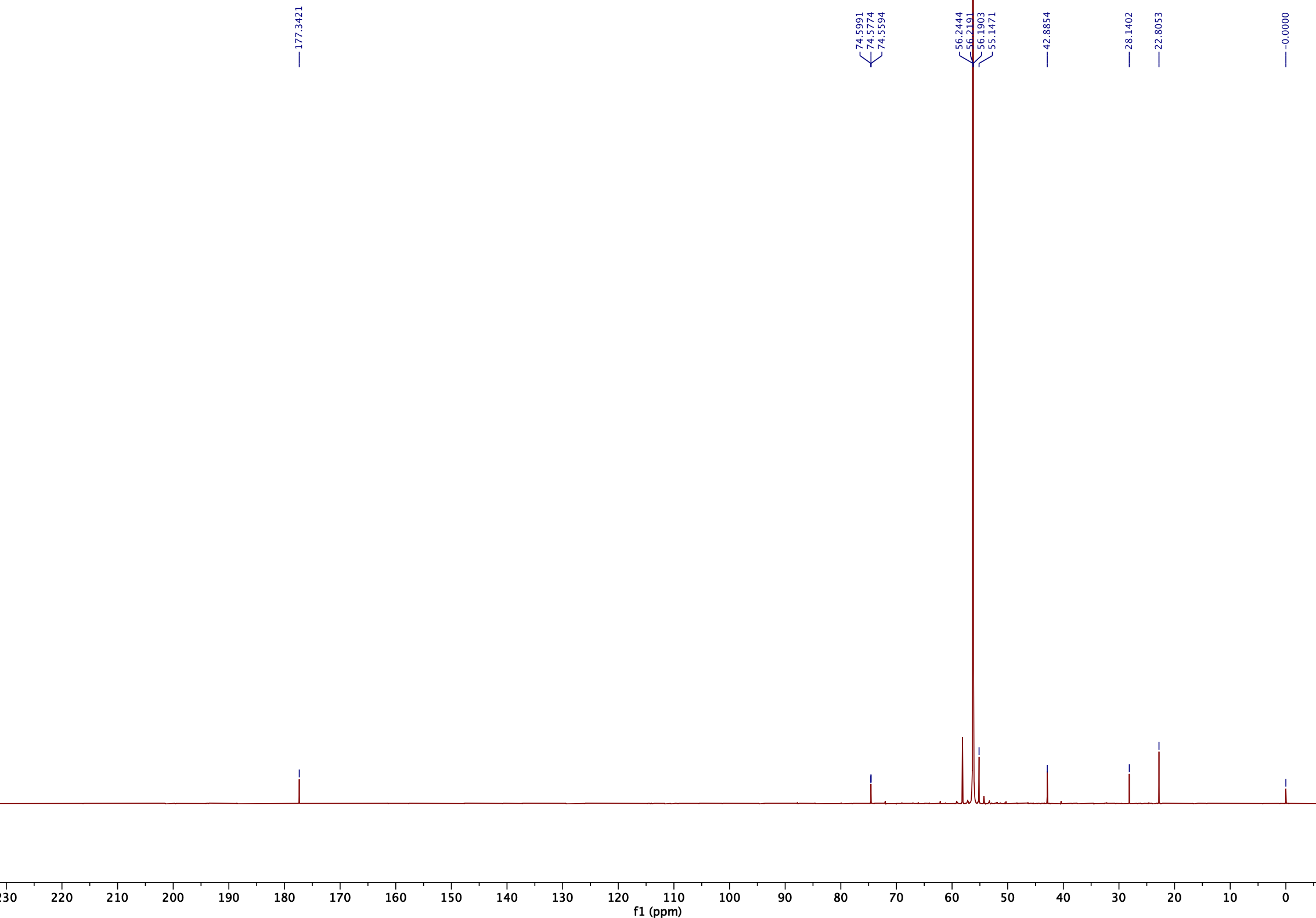

**Figure S28**

**3-Methyl-4-(tri(methyl-^13^*C*)ammonio)butanoate (^13^C-3M-4-TMAB)**

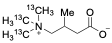

^1^H NMR (600 MHz, 300 K, D_2_O, TSP):

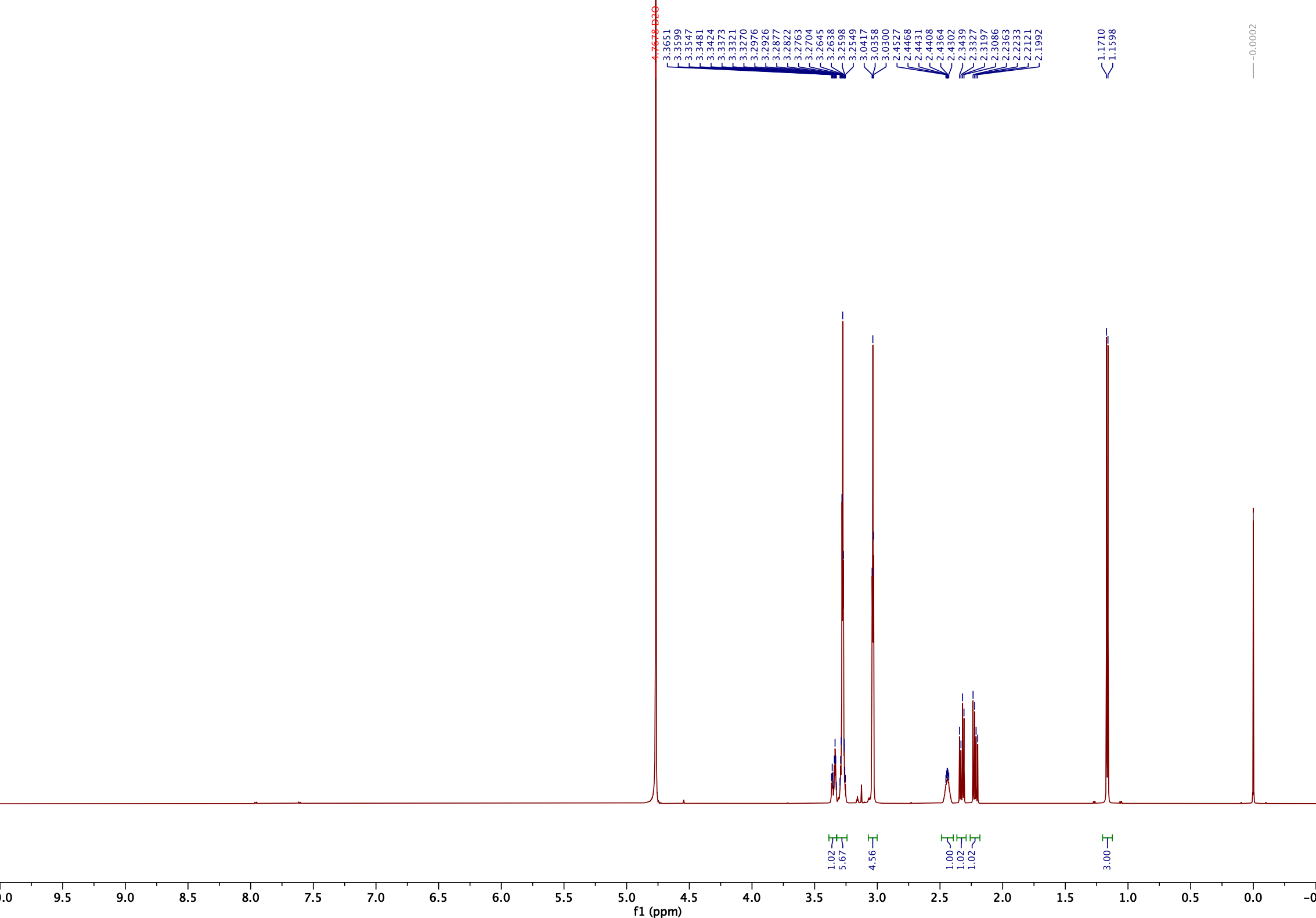

^1^H(^13^C) NMR (600 MHz, 300 K, D_2_O, TSP):

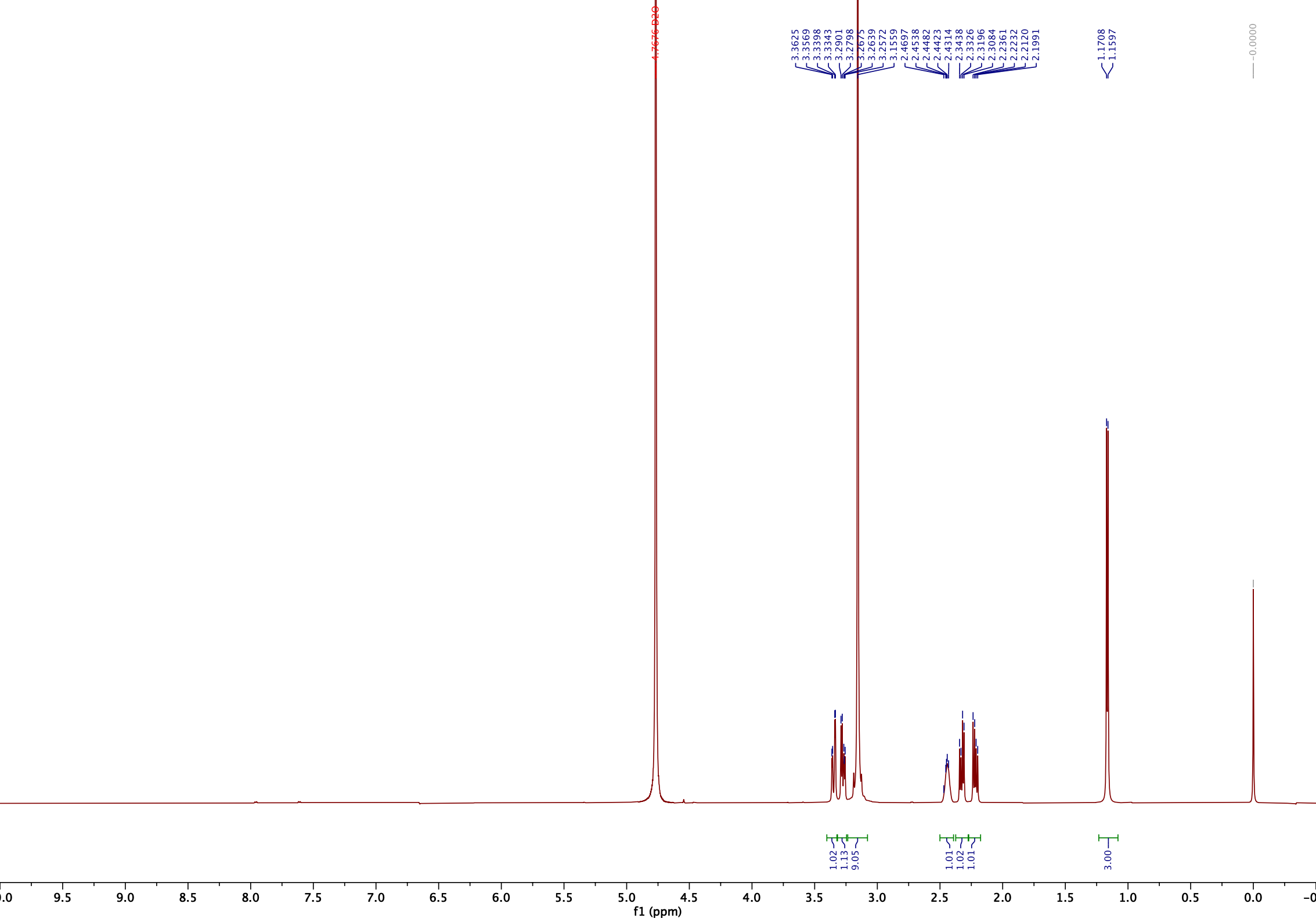

^13^C NMR (151 MHz, 300 K, D_2_O, TSP):

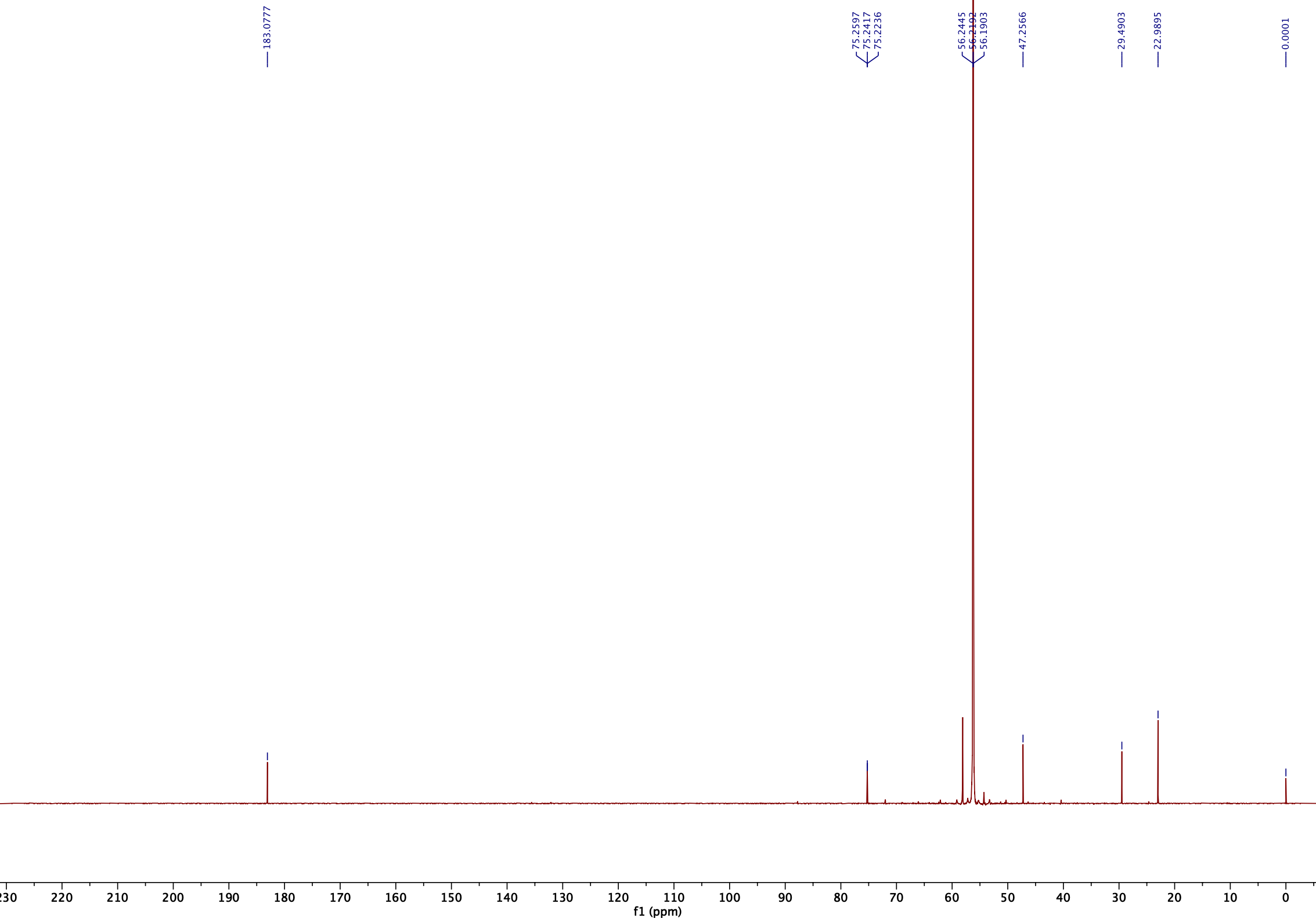

**Figure S29**

**5-(Trimethylammonio)pentanoate (5-AVAB)**

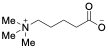

^1^H NMR (600 MHz, 300 K, D_2_O, TSP):

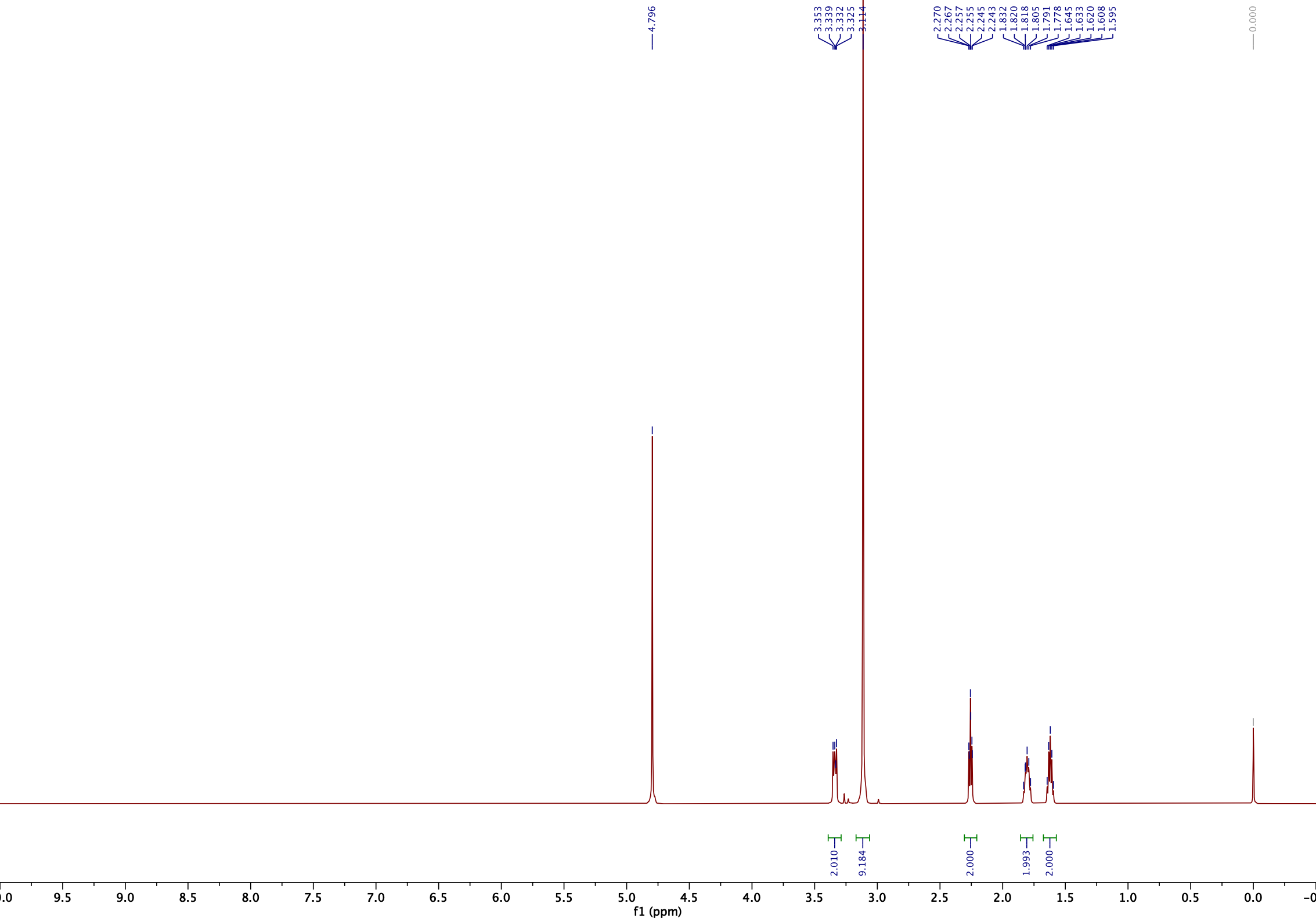

^13^C NMR (151 MHz, 300 K, D_2_O, TSP):

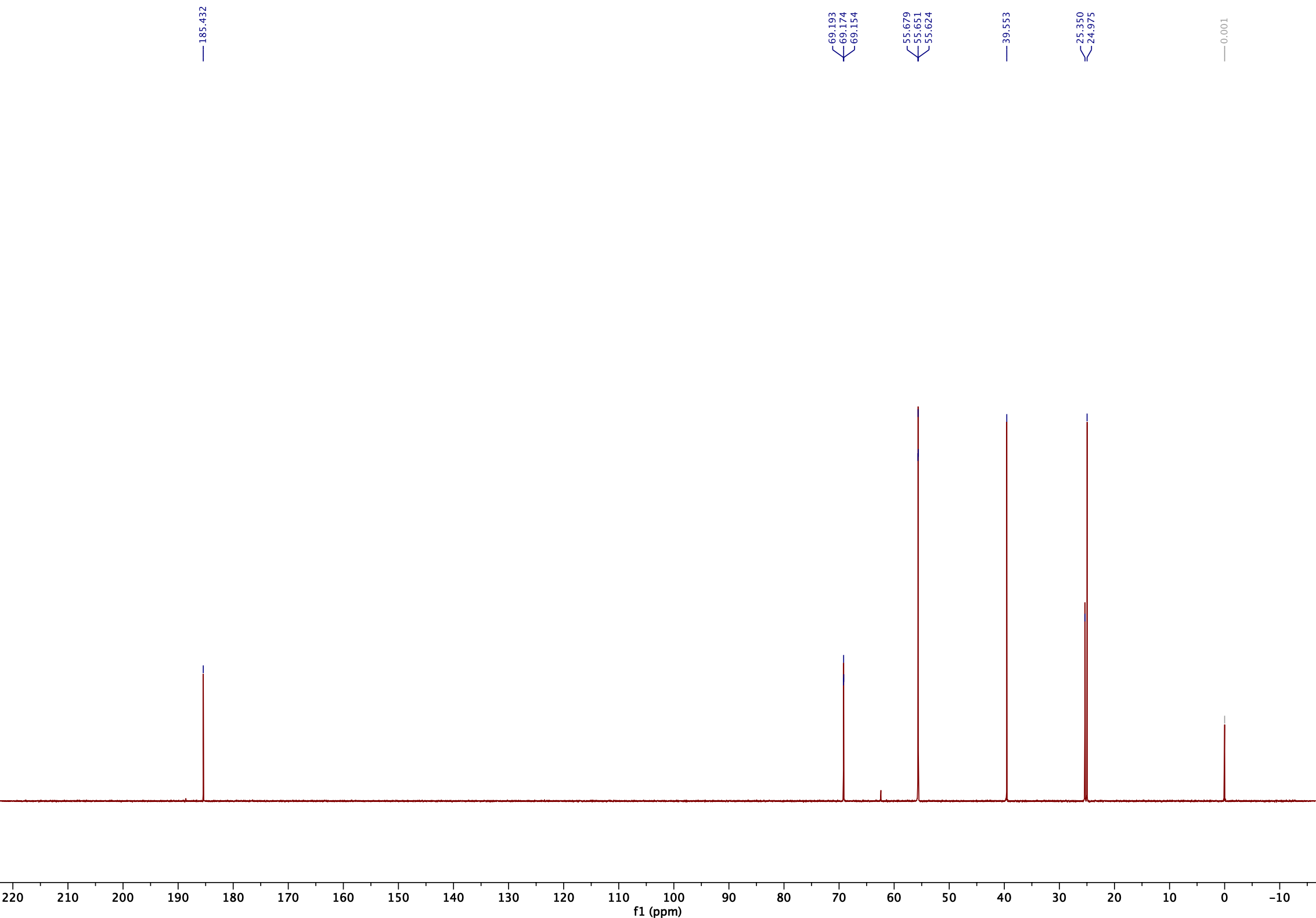
